# Choice-selective sequences dominate in cortical relative to thalamic inputs to nucleus accumbens, providing a potential substrate for credit assignment

**DOI:** 10.1101/725382

**Authors:** Nathan F. Parker, Avinash Baidya, Julia Cox, Laura Haetzel, Anna Zhukovskaya, Malavika Murugan, Ben Engelhard, Mark S. Goldman, Ilana B. Witten

**Affiliations:** Princeton Neuroscience Institute, Princeton University, Princeton NJ 08544; Department of Psychology, Princeton University, Princeton NJ 08544; Center for Neuroscience; Department of Neurobiology, Physiology and Behavior; Department of Ophthalmology and Vision Science, University of California Davis, Davis, CA 95616; Department of Physics and Astronomy, University of California Davis, Davis, CA 95616

## Abstract

How are actions linked with subsequent outcomes to guide choices? The nucleus accumbens, which is implicated in this process, receives glutamatergic inputs from the prelimbic cortex and midline regions of the thalamus. However, little is known about whether and how representations differ across these input pathways. By comparing these inputs during a reinforcement learning task in mice, we discovered that prelimbic cortical inputs preferentially represent actions and choices, whereas midline thalamic inputs preferentially represent cues. Choice-selective activity in the prelimbic cortical inputs is organized in sequences that persist beyond the outcome. Through computational modeling, we demonstrate that these sequences can support the neural implementation of reinforcement learning algorithms, both in a circuit model based on synaptic plasticity, and one based on neural dynamics. Finally, we test and confirm predictions of our circuit models by direct manipulation of nucleus accumbens input neurons. Thus, we integrate experiment and modeling to suggest neural solutions for credit assignment.

## Introduction

Multiple lines of experimental evidence indicate that the nucleus accumbens (NAc, part of the ventral striatum) is critical to reward-based learning and decision-making (Apicella et al., 1991; Cador et al., 1989; Carelli et al., 1993; Cox and Witten, 2019; Di Ciano et al., 2001; Everitt et al., 1991; Parkinson et al., 1999; Phillips et al., 1993, 1994; Robbins et al., 1989; Roitman et al., 2005; Setlow et al., 2003; Stuber et al., 2011; Taylor and Robbins, 1986). The NAc is a site of convergence of glutamatergic inputs from a variety of regions, including the prefrontal cortex and the midline thalamus, along with dense dopaminergic inputs from the midbrain (Brog et al., 1993; Do-Monte et al., 2017; Groenewegen et al., 1980; Hunnicutt et al., 2016; Otis et al., 2017; Phillipson and Griffiths, 1985; Poulin et al., 2018; Reed et al., 2018; Swanson, 1982; Wright and Groenewegen, 1995; Zhu et al., 2016).

An important mechanism underlying reward-based learning and decision-making is thought to be dopamine-dependent synaptic plasticity of glutamatergic inputs to the NAc that are co-active with a reward prediction error (RPE) in dopamine neurons (Fisher et al., 2017; Gerfen and Surmeier, 2011; Reynolds and Wickens, 2002; Russo et al., 2010). Such strengthening of glutamatergic inputs is thought to be central to learning, allowing actions that are followed by a rewarding outcome to be more likely to be repeated in the future (Britt et al., 2012; MacAskill et al., 2014; Steinberg et al., 2013; Tsai et al., 2009; Witten et al., 2011).

A central question in reinforcement learning is how actions and outcomes become associated with each other, even when they are separated in time (Asaad et al., 2017; Gersch et al., 2014; Sutton, 1988; Wörgötter and Porr, 2005). A possible mechanism that could contribute to solving this problem of temporal credit assignment in the brain is that neural activity in the glutamatergic inputs to the NAc provide a neural memory trace of previous actions. This could allow action representations from glutamatergic inputs and outcome information from dopaminergic inputs to overlap in time.

Whether glutamatergic inputs to the NAc indeed represent memories of previous actions is unclear. More broadly, what information is carried by glutamatergic inputs to the NAc during reinforcement learning, and whether different inputs provide overlapping or distinct streams of information, has not been examined systematically. To date, there have been relatively few recordings of cellular-resolution activity of glutamatergic inputs to the NAc during reinforcement learning, nor comparison of multiple inputs within the same task, nor examination of the timescale with which information is represented within and across trials. Furthermore, if glutamatergic inputs do indeed provide memories of previous actions, construction of a neurally plausible instantiation of an algorithm for credit assignment based on the measured signals remains to be demonstrated (for review of biological instantiation of reinforcement learning algorithms, see Joel et al., 2002).

To address these gaps, we recorded from glutamatergic inputs to the NAc during a probabilistic reversal learning task that we previously demonstrated was dopamine-dependent. In this task, dopamine neurons that project to the NAc encode RPE, and inhibition of dopamine neurons substitutes for a negative RPE (Parker et al., 2016). To compare activity in major cortical and thalamic input to the NAc core, here we combined a retrograde viral targeting strategy with cellular-resolution imaging to examine the input from prelimbic cortex (“PL-NAc”, part of medial prefrontal cortex) and that from the midline regions of the thalamus (“mTH-NAc”). We found that PL-NAc neurons preferentially encode actions and choices relative to mTH-NAc neurons, with choice-selective sequential activity that persists until the start of the subsequent trial. The long timescale through which a prior action is encoded in cortical inputs to the NAc provides the information to bridge actions, outcomes, and the subsequent choice. In addition, we demonstrated with computational modeling that these choice-selective sequences can support neural instantiations of reinforcement learning algorithms, either through dopamine-dependent changes in downstream synaptic weights, or dopamine-dependent changes in neural dynamics. Finally, we test and confirm a prediction of our models through direct optogenetic manipulation of PL-NAc neurons. Thus, by recording and manipulating glutamatergic inputs to the NAc and integrating these data with computational modeling, we provide specific proposals for how reinforcement learning could be implemented by neural circuitry.

## Results

### Cellular resolution imaging of glutamatergic inputs to the NAc during a probabilistic reversal learning task

Mice performed a probabilistic reversal learning task while inputs from thalamus or cortex were imaged (**Figure 1a**). A trial was initiated when the mouse entered a central nose poke, which prompted the presentation of a lever on either side. Each lever had either a high (70%) or low (10%) reward probability, with the identity of the high and low probability levers reversing in an unsignaled manner after a variable number of trials (see Methods for block reversal probabilities). After a variable delay (0-1s), either a sound (CS+) was presented at the same time as a reward was delivered to a central reward port, or another sound (CS−) was presented that signaled the absence of reward.

**Figure 1.**
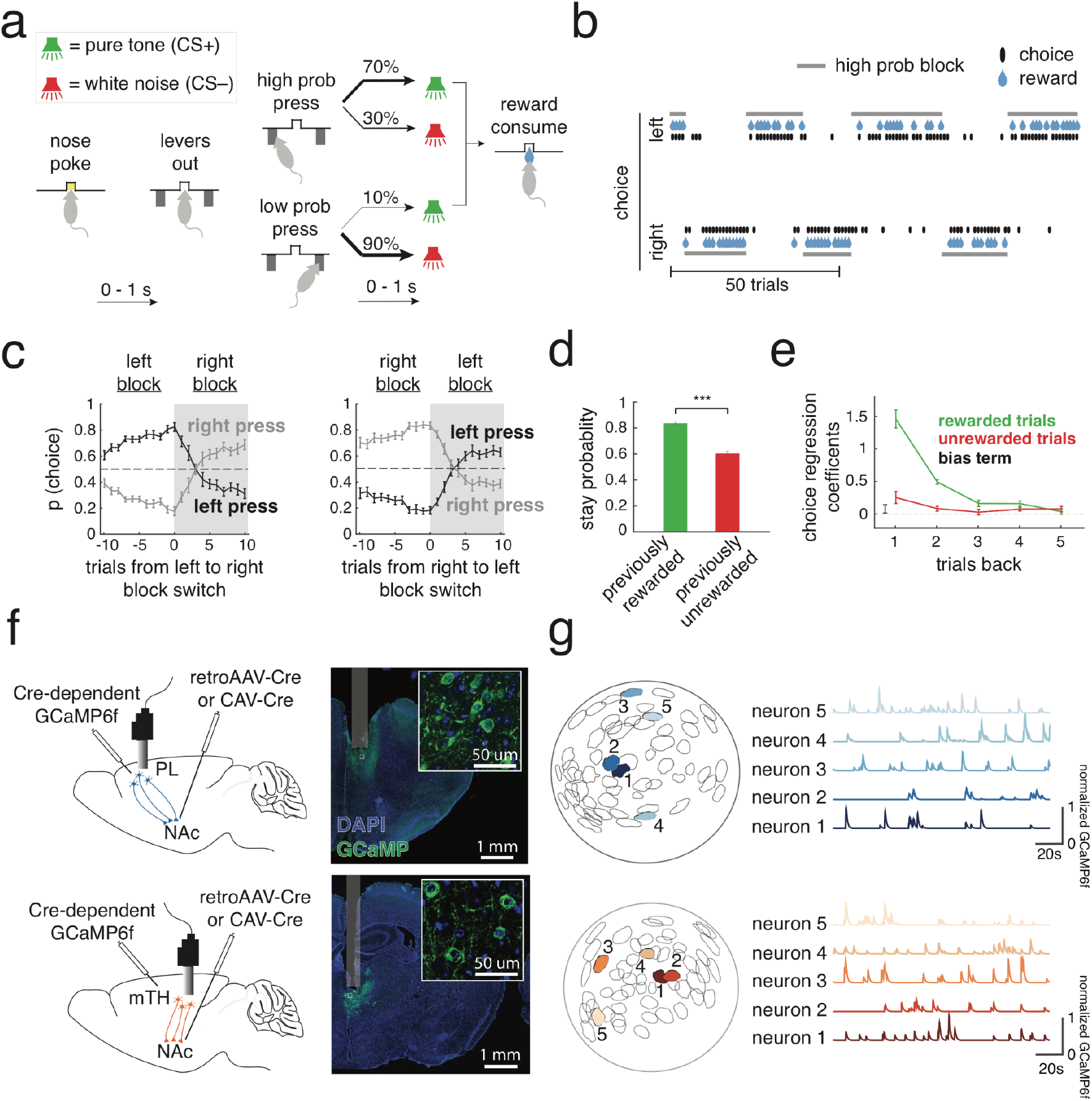
Cellular-resolution imaging of PL and mTH neurons that project to the NAc in mice performing a reinforcement learning task. **(a)** Schematic of probabilistic reversal learning task. Mice began a trial by entering a central nose poke (‘nose poke’) which resulted in the presentation of two levers (‘levers out’). A lever press resulted in either a rewarded (CS+, reward consumption) or unrewarded (CS-) outcome, with the probability of a given outcome dependent on whether the trial was in a right or left high-probability block. A high probability press (i.e. a left press in a left block, or a right press in a right block) led to reward on 70% of trials, while a low-probability press led to reward on 10%. **(b)** Example behavior of a mouse performing 120 trials during a recording session. The choice of the mouse (black marks) follows the identity of the higher probability lever as it alternates between a left and right block (horizontal grey bars). **(c)** Left, Probability of the mice choosing either the left or right lever 10 trials before and after a reversal from a left to right high-probability block. Right, same as left except choice probabilities following a right to left high-probability block. **(d)** Mice had a significantly higher stay probability following a rewarded versus unrewarded trial (*** p=5×10^-9^, two-tailed t-test, n=16 mice). **(e)** Coefficients from a logistic regression that uses choice and outcome from the previous five trials to predict choice on the current trial. The regression used two sets of predictors: (i) ‘Rewarded trials’ (green) which identify a previous trial as a rewarded right press (+1), rewarded left press (−1) or unrewarded press (0) and (ii) ‘Unrewarded trials’ (red), which identify a previous trial as an unrewarded right press (+1), unrewarded left press (−1) or rewarded trial (0). Positive regression coefficients correspond to a greater likelihood of the mouse making the same choice as that of the previous trial. Error bars in **c,d,e** represent s.e.m. across mice (n=16). **(f)** Left, surgical schematic for PL-NAc (top) and mTH-NAc (bottom) recordings showing the injection site and optical lens implant with miniature head-mounted microscope attached. Right, Coronal section from a PL-NAc (top) and mTH-NAc (bottom) mouse showing GCaMP6f expression in each respective recording site. Inset: confocal image showing GCaMP6f expression in individual neurons. **(g)** Left, example field of view from a recording in PL-NAc (top, blue) and mTH-NAc (bottom, orange) with five representative regions of interest (ROIs). Right, normalized GCaMP6f fluorescence traces from the five ROIs on the left. For visualization purposes, each trace was normalized by the peak fluorescence across the hour-long session.

As expected, mice switched the lever they were more likely to press following block reversals (**Figure 1b,c**). Similarly, mice were significantly more likely to return to the the previously chosen lever (i.e. stay) following rewarded, as opposed to unrewarded, trials (**Figure 1d**; p<0.0001: paired, two-tailed t-test between stay probabilities following rewarded and unrewarded trials across mice, n=16 mice), meaning that, as expected, mice were using previous choices and outcomes to guide behavior. A logistic regression to predict choice based on previous choices and outcomes indicated that mice relied on ~3 previous trials to guide their choices (**Figure 1e**; see Methods for choice regression details).

To image activity of glutamatergic input neurons to the NAc during this behavior, we injected a retroAAV or CAV2 to express Cre-recombinase in the NAc as well as an AAV2/5 to Cre-dependently express GCaMP6f in either the PL or mTH (**Figure 1f**). A gradient refractive index (GRIN) lens was implanted above either the PL or mTH (see **Supplementary Figure 1** for implant locations), and a head-mounted miniature microscope was used to image activity in these populations during behavior (**Figure 1f**, n=278 neurons in PL-NAc from n=7 mice, n=256 neurons in mTH-NAc from n=9 mice). An example field of view from a single recording session is shown for both PL-NAc neurons as well as mTH-NAc neurons (**Figure 1g**). Behavior between mice in the PL-NAc versus mTH-NAc cohorts was similar (**Supplementary Figure 2**).

### Actions are preferentially represented by PL-NAc neurons, while reward-predicting stimuli are preferentially represented by mTH-NAc neurons

Individual PL-NAc and mTH-NAc neurons displayed elevated activity when time-locked to specific behavioral events in the task (**Figure 2a**). Given the correlation between the timing of task events, as well as the temporal proximity of events relative to the time-course of GCaMP6f, we built a linear encoding model to properly relate neural activity to each event (Engelhard et al., 2019; Krumin et al., 2018; Lovett-Barron et al., 2019; Musall et al., 2019; Park et al., 2014; Parker et al., 2016; Pinto and Dan, 2015; Sabatini, 2019; Steinmetz et al., 2019). Briefly, time-lagged versions of each behavioral event (nosepoke, lever press, etc) were used to predict the GCaMP6f fluorescence in each neuron using a linear regression. This allowed us to obtain “response kernels”, which related each event to the GCaMP6f fluorescence in each neuron, while removing the potentially confounding (linear) contributions of correlated task events (**Figure 2b**; see Methods for details).

**Figure 2.**
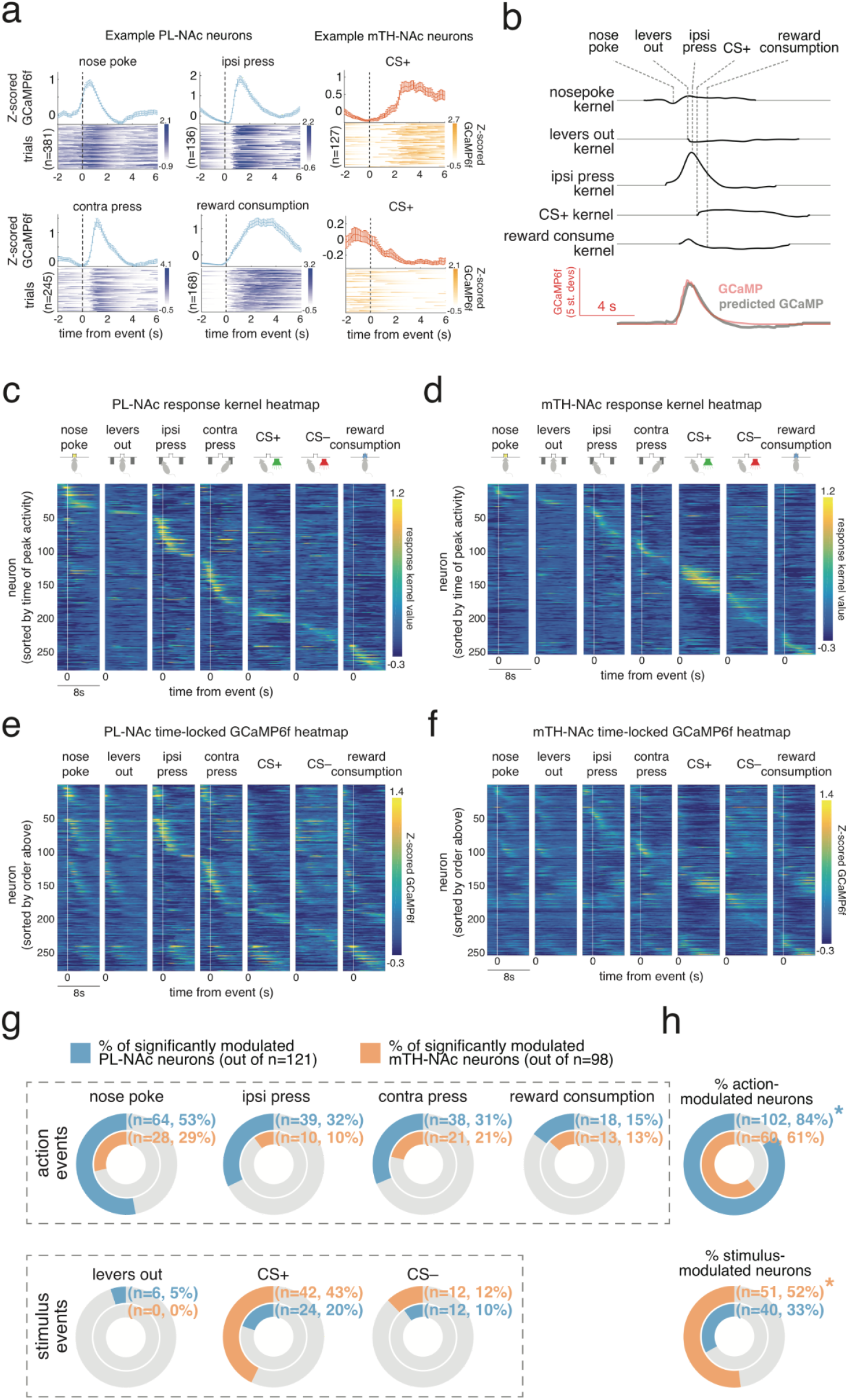
PL-NAc preferentially represents action events while mTH-NAc preferentially represents the CS+. **(a)** Time-locked responses of individual PL-NAc (blue) and mTH-NAc (orange) neurons to task events. **(b)** Kernels representing the response to each of the task events for an example neuron, generated from the encoding model. The predicted GCaMP trace in the model is the sum of the individual response kernels, aligned to the behavioral event times (see Methods). **(c)** Heatmap of response kernels generated from the encoding model in **b** from PL-NAc neurons. Heatmap is ordered by the time of the peak response across all behavioral events (n=278 neurons from 7 mice). **(d)** Same as **c** except heatmap is of response kernels from mTH-NAc recordings. (n=256 neurons from 9 mice) **(e)** Heatmap of mean Z-scored GCaMP6f fluorescence from PL-NAc neurons aligned to the time of each event in the task. Neurons are sorted by the order established in **c. (f)** Same as **e** except heatmaps are composed of GCaMP6f fluorescence from mTH-NAc neuron recordings. **(g)** Top row, fraction of neurons significantly modulated by action events (‘nose poke’, ‘ipsilateral lever press’, ‘contralateral lever press’, ‘reward’) in the PL-NAc (blue) and mTH-NAc (orange). For all action events, PL-NAc had a larger fraction of significantly modulated neurons compared with mTH-NAc. Bottom row, fraction of neurons in PL-NAc (blue) and mTH-NAc (orange) significantly modulated by stimulus events (‘levers out’, ‘CS+’ and ‘CS-’). In contrast to action events, two out of three stimulus events had a larger fraction of significantly modulated neurons in mTH-NAc compared with PL-NAc. Significance was determined using the linear model used to generate response kernels in **b**, see Methods for additional model details. **(h)** Top, a significantly larger fraction of event-modulated PL-NAc neurons encode at least one action event (P=0.0001: two-proportion Z-test comparing fraction of action-modulated PL-NAc and mTH-NAc neurons). Bottom, same as top except a significantly larger fraction of mTH-NAc neurons encode a stimulus event compared with PL-NAc (P=0.005: two-proportion Z-test comparing fraction of stimulus-modulated neurons between PL-NAc and mTH-NAc).

To visualize the response kernels, we plotted them as a heatmap, where each row was the response kernel for a particular neuron associated with each behavioral event. This heatmap was then ordered by the time of peak kernel value across all behavioral events. Visual observation revealed a clear difference between the PL-NAc and mTH-NAc populations – PL-NAc neurons were robustly modulated by the action-events in our task (**Figure 2c**; kernel values associated with ‘nose poke, ‘ipsilateral lever press’, ‘contralateral lever press’ and ‘reward consumption’) while mTH-NAc neurons appeared to be most strongly modulated by the stimulus-events, specifically the positive reward auditory cue (**Figure 2d**, kernel values associated with ‘CS+’).

Examination of the GCaMP6f fluorescence time-locked to each behavioral event (rather than the encoding model-derived response kernels) revealed similar observations of action encoding in PL-NAc and CS+ encoding in mTH-NAc (**Figure 2e,f**). While this time-locked GCaMP6f heatmap displays neurons which appear to respond to multiple events (**Figure 2e**, see neurons approximately 50-100 that show elevated activity to ‘ipsilateral lever press’, ‘levers out’ and ‘nose poke’), this impression is likely a result of the temporal correlation between neighboring behavioral events, which our encoding model accounts for. To illustrate this, we applied our encoding model on a population of simulated neurons that responded only to the lever press events. In this simulated data, we observed a similar multi-peak heatmap when simply time-locking the simulated GCaMP6f fluorescence, but this multi-peak effect is eliminated by the use of our encoding model, which recovers the true relationship between GCaMP6f fluorescence and behavior in the simulated data (**Supplementary Figure 3**).

This encoding model was used to identify neurons in the PL-NAc and mTH-NAc populations that were significantly modulated by each event in our task (significance was assessed by comparing the encoding model with and without each task event, see Methods). We found that a similar fraction of PL-NAc and mTH-NAc neurons were modulated by at least one task event (PL-NAc: n=121/278 neurons from 7 mice; mTH-NAc: n=98/256 neurons from 9 mice). Of these neurons that were selective to at least one task event, the selectivity for actions versus sensory stimuli differed between the two populations (**Figure 2g,h**). In particular, more PL-NAc neurons were modulated by at least one *action* event (nose poke, ipsilateral lever press, contralateral lever press and reward consumption; 102/121 PL-NAc neurons; 60/98 mTH-NAc neurons; P=0.0001: two-proportion Z-test comparing fraction of action-modulated neurons between PL-NAc and mTH-NAc). By contrast, a significantly larger fraction of mTH-NAc neurons were modulated by at least one stimulus cue (levers out, CS+ and CS-; 51/98 mTH-NAc neurons; 40/121 PL-NAc neurons; P=0.005, two-proportion Z-test comparing fraction of stimulus-modulated neurons between PL-NAc and mTH-NAc).

### PL-NAc neurons preferentially encode choice relative to mTH-NAc neurons

This preferential representation of actions in PL-NAc relative to mTH-NAc suggests that lever choice (contralateral versus ipsilateral to the recording site) could also be preferentially encoded in PL-NAc. Indeed, a significantly larger fraction of neurons were choice-selective in PL-NAc compared with mTH-NAc (**Figure 3a**; PL-NAc: 92/278 (33%); mTH-NAc: 42/256 (16%); P=9.9×10^-6^: two-proportion Z-test; significant choice-selectivity was determined with a nested comparison of the encoding model with and without choice information, see Methods). A logistic regression population decoder supported this observation of preferential choice-selectivity in PL-NAc relative to mTH-NAc. Choice decoding using neural activity of simultaneously recorded PL-NAc neurons was significantly more accurate compared with decoding using mTH-NAc activity (**Figure 3b**; PL-NAc: 72±3%, mTH-NAc: 60±2%, choice decoding accuracy from a logistic regression with activity from multiple, random selections of 10 simultaneously imaged neurons, mean±s.e.m. across mice; P=0.0065: unpaired, two-tailed t-test comparing peak decoding accuracy across mice between PL-NAc, n=6 mice, and mTH-NAc, n=9 mice).

**Figure 3.**
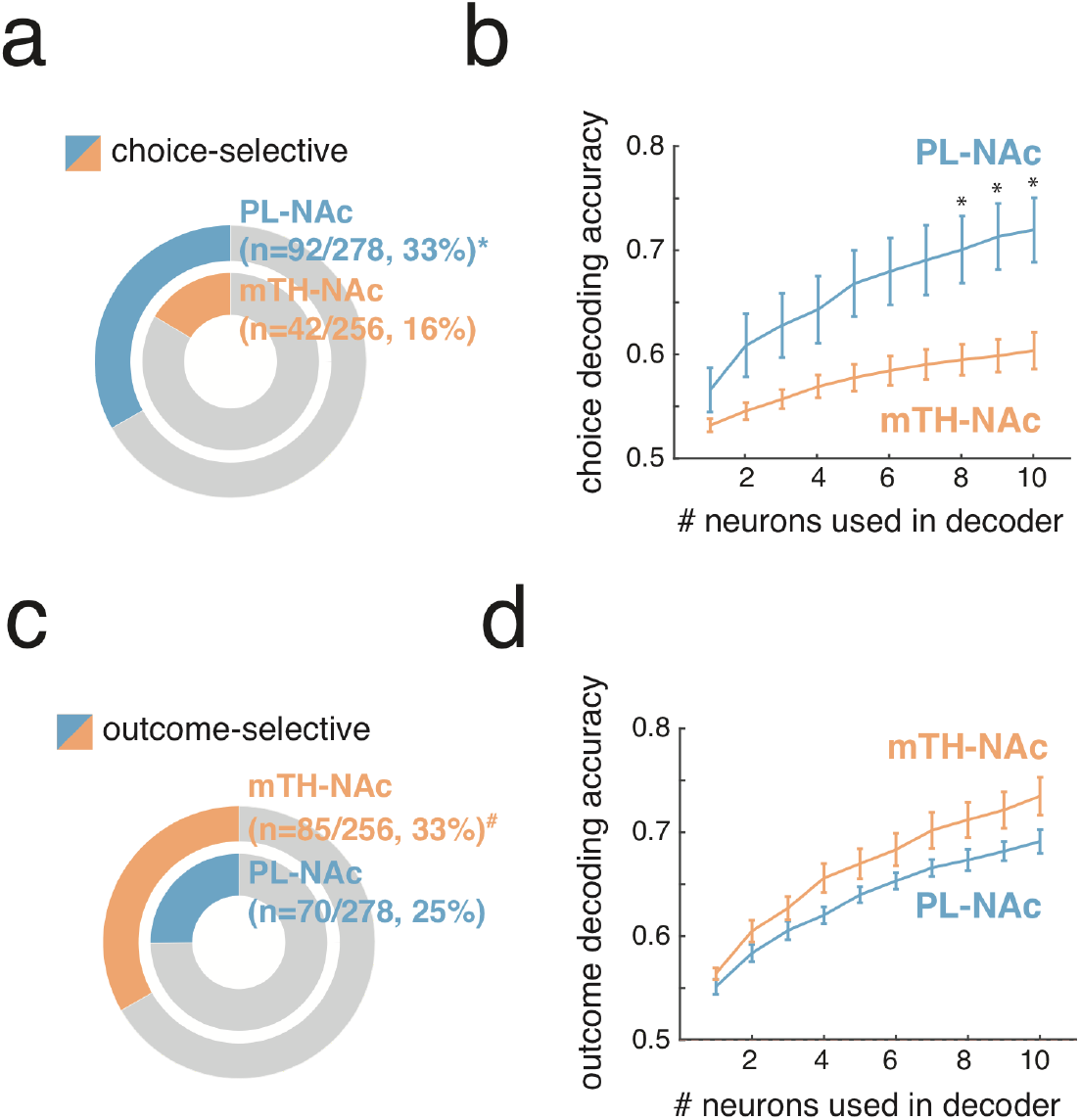
PL-NAc preferentially represents choice but not outcome relative to mTH-NAc. **(a)** Fraction of choice-selective neurons in PL-NAc (n=92 out of 278 neurons from 7 mice) and mTH-NAc (n=42 out of 256 neurons from 9 mice). A significantly larger fraction of PL-NAc neurons were choice-selective (P=9.9×10^-6^: two-proportion Z-test comparing fraction of choice-selective PL-NAc and mTH-NAc neurons). **(b)** Choice decoding accuracy using neural activity from one to ten randomly-selected, simultaneously imaged neurons around the time of the lever press. The PL-NAc population (n=6 mice) more accurately decodes the choice of the trial compared with mTH-NAc (n=9 mice; peak decoding accuracy of 72+/-3% for PL-NAc and 60+/-2% for mTH-NAc). Asterisks indicate P<0.05, unpaired two-tailed t-test comparing decoding accuracy between PL-NAc (n=6) and mTH-NAc (n=9) mice. **(c)** Fraction of outcome-selective neurons in mTH-NAc (n=85 out of 256 neurons, n=9 mice) and PL-NAc (n=70 out of 278 neurons, n=7 mice). A significantly larger fraction of mTH-NAc neurons were outcome-selective (P=0.038: two-proportion Z-test comparing fraction of outcome-selective PL-NAc and mTH-NAc neurons). **(d)** Outcome decoding accuracy using neural activity after the time of the CS from one to ten randomly-selected, simultaneously imaged neurons in mTH-NAc in orange (n=9 mice, peak decoding accuracy of 74+/-2%) and PL-NAc in blue (n=6 mice, peak decoding accuracy of 69+/-1%). Error bars in **b,d** indicate s.e.m. across mice.

In contrast to the preferential representation of choice in PL-NAc compared to mTH-NAc, there was a larger fraction of neurons in mTH-NAc that encoded outcome (CS identity or reward consumption) compared to PL-NAc (**Figure 3c**; mTH-NAc: 85/256 (33%), PL-NAc: 70/278 (25%); P=0.038: two-proportion Z-test; significant outcome-selectivity was determined using a nested comparison of the encoding model with and without CS identity and reward consumption information, see Methods). However, while outcome decoding accuracy in mTH-NAc was slightly better relative to PL-NAc (**Figure 3d**; mTH-NAc: 73±2%, PL-NAc: 69±1%), this difference was not statistically significant (P=0.11: unpaired, two-tailed t-test comparing peak decoding accuracy across mice between PL-NAc (n=6 mice) and mTH-NAc (n=9 mice)). These results suggest that, unlike the preferential choice representation observed in PL-NAc over mTH-NAc, outcome was more similarly represented between these two populations. This is presumably due to the fact that both CS+ and reward consumption responses contribute to outcome representation, and although more neurons encoded CS+ in mTH-NAc, the opposite was true for reward consumption (**Figure 2c,d**).

To determine if choice or outcome decoding in either population depended on recording location, we aligned GRIN lens tracks from the histology to the Allen atlas (see Methods). We found no obvious relationship between the strength of either choice or outcome decoding and recording location in either PL-NAc or mTH-NAc (**Supplementary Figure 4**).

### PL-NAc neurons display choice-selective sequences that persist into the next trial

We next examined the temporal organization of choice-selective activity in PL-NAc neurons. Across the population, choice-selective PL-NAc neurons displayed sequential activity with respect to the lever press that persisted for >4s after the press (**Figure 4a-c**; see **Supplementary Figure 5** for sequences without peak-normalization). These sequences were visualized by time-locking the GCaMP6f fluorescence of choice-selective neurons with respect to the lever press, rather than with the encoding model from the earlier figures. The robustness of these sequences was confirmed using a cross-validation procedure, in which the order of peak activity across the PL-NAc choice-selective population was first established using half the trials (**Figure 4b**, ‘train’), and then the population heatmap was plotted using the same established ordering and activity from the other half of trials (**Figure 4c**, ‘test’). To quantify the consistency of these sequences, we correlated the neurons’ time of peak activity in the ‘training’ and ‘test’ data, and observed a strong correlation (**Figure 4d**; R^2^=0.81, P=6.4×10^-23^, n = 92 neurons from 7 mice). Additionally, the ridge-to-background ratio, a metric used to confirm the presence of sequences (Akhlaghpour et al., 2016; Harvey et al., 2012; Kondo et al., 2017) was significantly higher when calculated using the PL-NAc choice-selective sequences compared with sequences generated using shuffled data (**Supplementary Figure 6a-c**; P<0.001, t-test between ratio calculated from unshuffled data and 500 iterations of shuffled data).

In contrast, choice-selective sequential activity in the mTH-NAc population was significantly less consistent than in PL-NAc (**Supplementary Figure 7a-d**; Z = 2.34, P=0.0096: Fisher’s Z, comparison of correlation coefficients derived from comparing peak activity between ‘test’ and ‘training’ data from PL-NAc versus mTH-NAc). Additionally, while the ridge-to-background ratio of the sequences generated using mTH-NAc activity was significantly higher than that using shuffled data, (**Supplementary Figure 6d-f**; P<0.001) this ratio was also significantly lower than that obtained from PL-NAc sequences (2.36+/-0.12 mean+/-sem ratio for choice-selective mTH-NAc neurons, n=42, vs 3.01+/-0.12 mean+/-sem ratio for choice-selective PL-NAc, n=92; P=0.001: unpaired, two-tailed t-test comparing ratio between PL-NAc and mTH-NAc neurons). The ridge-to-background ratio of both the PL-NAc and mTH-NAc sequences did not significantly change across either a block or recording session (**Supplementary Figure 8a-d**).

A striking feature of these choice-selective sequences in PL-NAc was that they persisted for seconds after the choice, potentially providing a neural ‘bridge’ between action and outcome. To further quantify the timescale of choice encoding, both within and across trials, we used activity from simultaneously imaged neurons at each timepoint in the trial to predict the mouse’s choice (with a decoder based on a logistic regression using random combinations of 10 simultaneously imaged neurons to predict choice). Choice on the current trial could be decoded above chance for ~7s after the lever press, spanning the entire trial (including the time of reward delivery and consumption), as well as the beginning of the next trial (**Figure 4e**; P<0.01: unpaired, two-tailed t-test of decoding accuracy across mice at the time of the next trial nose poke compared with chance decoding of 0.5, n=6 mice). Choice on the previous or subsequent trial was not represented as strongly as current trial choice (**Figure 4e**; in all cases we corrected for cross-trial choice correlations with a weighted decoder, see Methods) and choice from two trials back could not be decoded above chance at any time point in the trial (**Supplementary Figure 8e**). We also examined the temporal extent of choice encoding in the mTH-NAc population (**Supplementary Figure 7e**). Similar to PL-NAc, we observed that decoding of the mice’s choice persisted up to the start of the next trial. However, the peak decoding accuracy across all time points in the trial was lower in mTH-NAc (59%±0.2%) compared with PL-NAc (73±0.2%).

### Choice-selective sequences in PL-NAc neurons, in combination with known anatomy, can provide a substrate for temporal difference (TD) learning

Thus far, we observed that choice-selective sequences in PL-NAc neurons encoded the identity of the chosen lever for multiple seconds after the lever press. This sequential activity bridged the gap in time between a mouse’s action and reward feedback, and therefore contained information that could be used to solve the task (**Figure 4**). But how could a biologically realistic network use sequences to implement this task? In particular, how could the synaptic outputs of PL-NAc neurons that were active at the *initiation* of the choice-selective sequence be strengthened by an outcome that occurred toward the *end* of the sequence? This represents a temporal credit-assignment problem and may be critical to the ability of the system to learn to generate the appropriate choice-selective sequence for a given trial.

To address this question, we developed a circuit-based computational model that could perform this task based on the observed choice-selective sequences in PL-NAc neurons. This was achieved by a synaptic plasticity model that implemented a temporal difference (TD) reinforcement learning algorithm by combining the recorded choice-selective sequential activity of PL-NAc neurons with the known connectivity of downstream structures (**Figure 5a,b**). The goal of TD learning is for weights to be adjusted in order to predict the sum of future rewards, or “value” (Dayan and Niv, 2008; O’Doherty et al., 2003; Sutton and Barto, 1998; Tsitsiklis and Van Roy, 1997). When this sum of future rewards changes, such as when an unexpected reward is received or an unexpected predictor of reward is experienced, a TD reward prediction error (RPE) occurs and adjusts the weights to reduce this error. The error signal in the TD algorithm closely resembles the RPE signal observed in ventral tegmental area (VTA) dopamine neurons (Parker et al., 2016; Schultz, 1998; Schultz et al., 1997), but how this signal is computed remains an open question.

**Figure 4.**
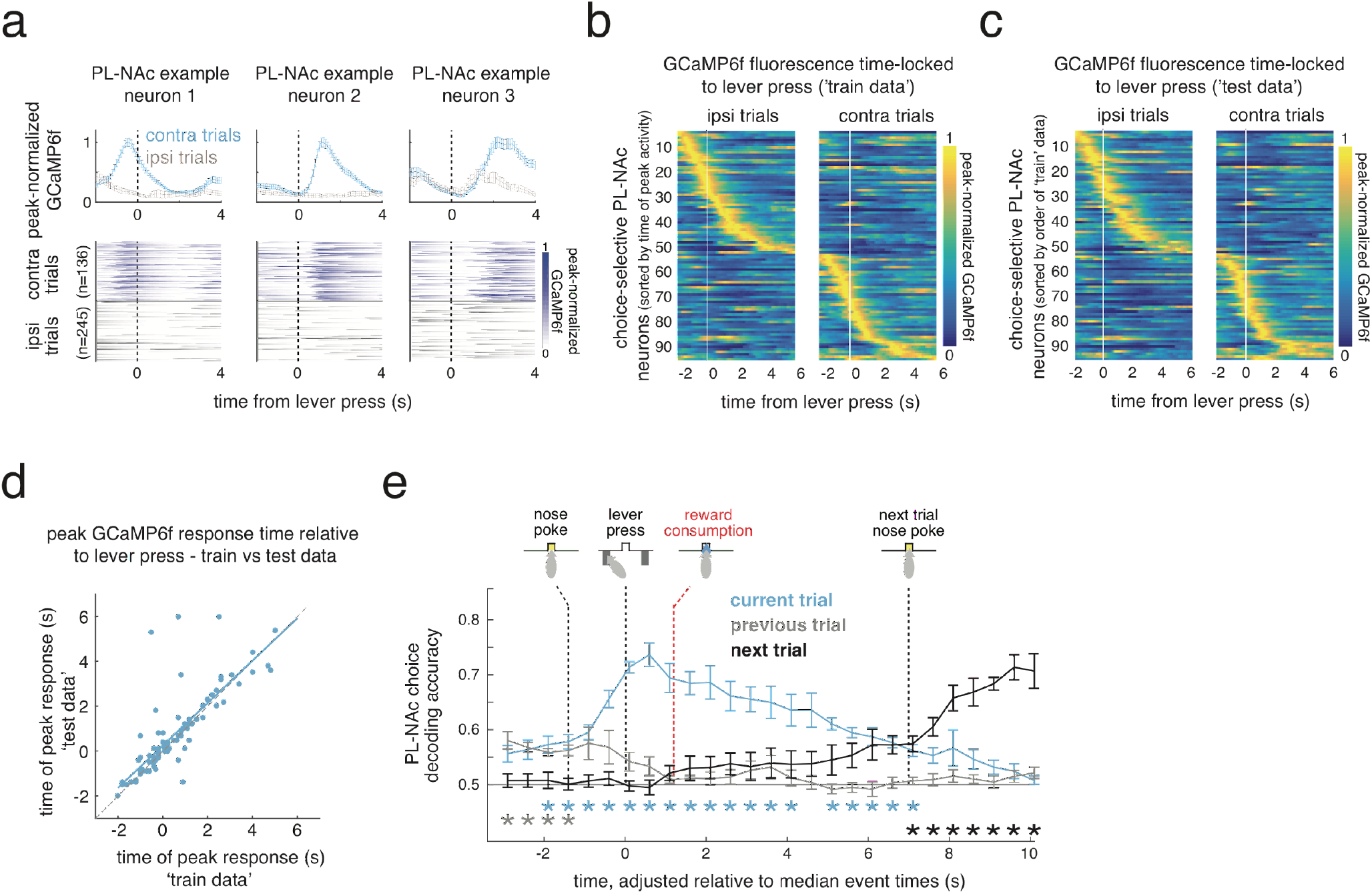
Choice-selective sequences in PL-NAc persist into the subsequent trial, beyond arrival at the central reward port. **(a)** Top, average peak-normalized GCaMP6f fluorescence of three simultaneously imaged PL-NAc choice-selective neurons with different response times relative to the lever press. Error bars are s.e.m across trials. Bottom, heatmaps of GCaMP6f fluorescence across trials relative to an ipsilateral (blue) and contralateral (grey) press. **(b,c)** Heatmaps demonstrating sequential activation of choice-selective PL-NAc neurons (n=92/278 neurons). Each row is the average GCaMP6f fluorescence time-locked to the ipsilateral (left column) and contralateral (right column) lever press for a neuron, normalized by the neuron’s peak average fluorescence. In **b** (‘train data’), heatmap is generated using a randomly selected half of trials and ordered by the time of each neurons’ peak activity. In **c** (‘test data’), the peak-normalized, time-locked GCaMP6f fluorescence from the other half of trials was used while maintaining the order from ‘train data’ in **b**. This cross-validation procedure ensures that sequences are not an artifact of the sorting. **(d)** Correlation between time of peak activity using the ‘train’ (horizontal axis) and ‘test’ (vertical axis) trials for choice-selective PL-NAc neurons in response to a contralateral or ipsilateral lever press (R^2^ = 0.81, P = 6.4×10^-23^, n = 92 neurons). **(e)** Average decoding accuracy of the mice’s choices on the current (blue), previous (grey) and next (black) trial as a function of GCaMP6f fluorescence throughout the current trial. GCaMP6f fluorescence is taken from 100 random selections per mouse of 10 simultaneously imaged PL-NAc neurons (each trial’s activity is adjusted in a piecewise linear manner relative to the median time of the nose poke, lever press and next trial nose poke, see Methods for details). Error bars are s.e.m. across mice (n=6 mice). Red dashed line indicates median onset of reward consumption. Asterisks indicate significant decoding accuracy above chance, P<0.01, two-tailed, one-sample t-test across mice.

In our synaptic plasticity model, the PL-NAc sequences enabled the calculation of the RPE in dopamine neurons. Over the course of a series of trials, this error allowed a reward that occurred at the end of the PL-NAc sequence to adjust synaptic weights at the beginning of the sequence (**Figure 5a,b**). Our model generated an RPE in dopamine neurons based on minimal assumptions: i) the choice-selective sequences in PL-NAc neurons that we report here (**Figure 4b**), ii) established (but simplified) anatomical connections between PL, NAc, and VTA neurons (see circuit diagram in **Figure 5a**), and iii) dopamine- and activity-dependent modification of the synapses connecting PL and NAc neurons (with a synaptic eligibility decay time constant of 0.6s, consistent with Gerstner et al., 2018 and Yagishita et al., 2014). In addition to the proposed circuit architecture of **Figure 5a**, we present several variant circuits, which also are able to calculate an RPE signal using the recorded choice-selective PL-NAc sequences (**Supplementary Figure 9**).

**Figure 5.**
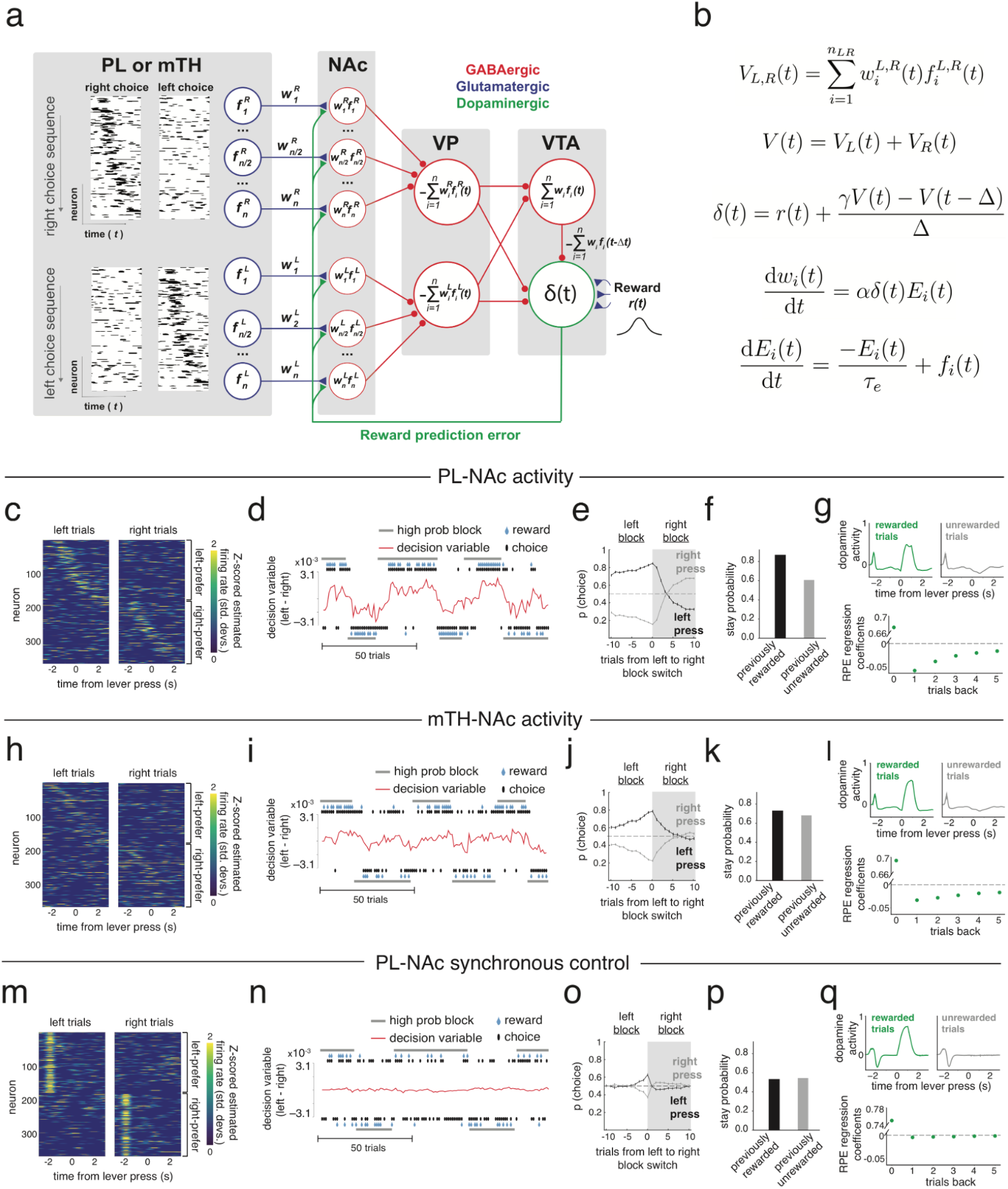
Choice-selective sequences recorded in PL-NAc, in combination with known downstream connectivity, can implement a temporal difference (TD) learning model based on synaptic plasticity. **(a)** Schematic of circuit architecture used in the model. Actual model implementation used single-trial recorded PL-NAc or mTH-NAc responses as input (see Results and Methods for model details). Choice-selective PL-NAc or mTH-NAc neurons, each with a temporal activity pattern defined as 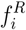 for right-preferring neurons and 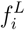 for left-preferring neurons, form synapses with weights 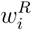 and 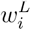 onto GABAergic NAc projection neurons. Dopamine (RPE)-dependent changes in the synaptic strengths 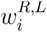, of these inputs enable the NAc neurons to approximately convey the value of a left or right choice at any time. These NAc neurons converge upon DA neurons through a fast (NAc-VP-DA) and slow, delayed by one synaptic connection (NAc-VP-VTA GABA-DA) pathway to form the temporal differencing operation required to generate the RPE of TD learning. VP neurons serve as a site of convergence (Oorschot, 1996; Kimura et al., 1996) and invert the sign of NAc inputs to provide the correct sign for generating an RPE, producing populations representing left or right choice value. DA neurons additionally receive an external reward signal *r*(*t*). See **Supplementary Figure 9** for alternative, mathematically equivalent circuit architectures. **(b)** Equations for the model as instantiated by the circuit schematic in **a**. *α* defines the learning rate, *τ_e_* defines the decay time constant for the PL-NAc synaptic eligibility trace *E*(*t*), Δ is the delay generated by the VTA GABA interneuron, *γ* is the discount in value associated with a time step Δ and *n_L,R_* is the total number of left and right-preferring neurons. The individual sums of left- and right-preferring NAc neuron activity is given by *V_L_* and *V_R_*, respectively, and the sum of the left and right value is *V*. **(c)** Heatmap of single trial PL-NAc estimated firing rates used as input to the model (see Methods for details). **(d)** Behavior of the synaptic plasticity model for 120 example trials spanning a series of high-probability block reversals. The decision variable (red trace, see Methods) along with the predicted choice of the model (black dots) follows the identity of the higher probability lever as it alternates between a left and right block (horizontal grey bars). **(e)** Choice probability of the model for the left (black) and right (grey) presses following a left-to-right block reversal. The model is able to switch to the new lever following a block reversal, similar to the mice’s behavior in **Figure 1c. (f)** Predicted stay probability of the synaptic plasticity model following rewarded and unrewarded trials. Similar to the behavior of mice in **Figure 1d**, the model predicts a higher stay probability following a rewarded trial compared with an unrewarded trial. **(g)** Top, simulated VTA dopamine neuron activity throughout the trial, averaged across rewarded trials (green) and unrewarded trials (grey). Bottom, coefficients from a linear regression that uses outcome of the current and previous five trials to predict dopamine neuron activity following outcome feedback (see Methods for details). Model dopamine activity follows the pattern predicted by the encoding of an RPE as observed in recorded dopamine activity (Bayer and Glimcher, 2005; Parker et al., 2016; see **Supplementary Figure 10** for comparison to experimental recordings). **(h-l)** Same as **c-g** except using the estimated firing rates from mTH-NAc instead of PL-NAc single trial activity. The mTH-NAc model input generates appropriate behavior; however, the performance is worse than that using PL-NAc input, with less and slower modulation of the decision variables, and weaker modulation of DA activity by previous trial outcomes. **(m)** Control model where choice-selective PL-NAc neurons fire synchronously at the approximate onset of the sequence (2s before the lever press), rather than sequentially. **(n-q)** Same as **d-g,** except results from using the control model with synchronous choice-selective PL-NAc activity. Unlike the sequential model, synchronous activity is not able to generate appropriate behavior **(n-p)** nor is it able to produce an RPE signal **(q).** Note that the increase in choice probability just prior to a block reversal in the synchronous case (panel **o**), and also in panels **j** and **e**, is due to the statistical structure of the block reversal criterion, rather than due to learning (see Methods).

In more detail, our synaptic plasticity model took as inputs experimental, single-trial recordings of choice-selective, sequentially active PL neurons (**Figure 5c**, see Methods). These inputs represented temporal basis functions (*f_i_*(*t*) in **Figure 5a**) for computing the estimated value of making a left or right choice. These basis functions are weighted in the NAc by the strength *w_i_* of the PL-NAc synaptic connection and summed together to create a (sign-inverted) representation of the estimated value, at time t, of making a left choice, *V_L_*(*t*), or right choice, *V_R_*(*t*). To create the RPE observed in DA neurons requires that the DA neuron population receive a fast, positive value signal *V*(*t*) and a delayed negative value signal *V*(*t* – Δ), as well as a direct reward signal (**Figure 5b**). In the instantiation of our model shown in **Figure 5a**, the summation of NAc inputs and sign-inversion occurs in the ventral pallidum (VP), so that the fast value signal is due to direct VP to DA input. The delayed negative signal to the DA population is due to a slower, disynaptic pathway that converges first upon the VTA GABA neurons, so that these neurons encode a value signal as observed experimentally (Eshel et al., 2015). The temporal discounting factor 7 is implemented through different strengths of the two pathways to the VTA DA neurons (**Figure 5b**).

Learning is achieved through DA-dependent modification of the PL-NAc synaptic strengths. We assume that PL-NAc neuronal activity leads to an exponentially decaying “eligibility trace” (Gerstner et al., 2018; Sutton and Barto, 1998). The correlation of this presynaptically driven eligibility trace with DA input then drives learning (**Figure 5b**). Altogether, this circuit architecture (as well as the mathematically equivalent architectures shown in **Supplementary Figure 9**) realizes a temporal difference learning algorithm for generating value representations in the ventral basal ganglia, providing a substrate for the selection of proper choice based on previous trial outcomes. The synaptic plasticity model was able to correctly perform the task and recapitulate the mice’s behavior, achieving a comparable rate of reward (47.5% for the model versus 47.6% for the mice). Similar to mice, the model alternated choice following block reversals (**Figure 5d,e**; compare to **Figure 1b,c**; choice based upon a probabilistic readout of the difference between right and left values at the start of the sequence, see Methods) and had a higher stay probability following rewarded trials relative to unrewarded trials (**Figure 5f**; compare to **Figure 1d**).

Appropriate task performance was achieved through the calculation of an RPE signal in VTA dopamine neurons. The RPE signal was evident within a trial, based on the positive response to rewarded outcomes and negative response to unrewarded outcomes (**Figure 5g, top**). The RPE signal was also evident across trials, based on the negative modulation of the dopamine outcome signal by previous trial outcomes (**Figure 5g, bottom**, multiple linear regression similar to Bayer & Glimcher, 2005 and Parker et al., 2016).

Several features of the model neuron responses resembled those previously observed experimentally. The model dopamine responses were similar to results obtained from recorded dopamine activity in the same task (**Supplementary Figure 10**; Parker et al., 2016). The VTA GABA interneuron had a sustained value signal, due to the converging input of the transient, sequential value signals from NAc/VP (**Supplementary Figure 11**), replicating the sustained value signal in VTA GABA interneurons recently found in monosynaptic inputs to the VTA dopamine neurons (Cohen et al., 2012). We note that, alternatively, the VP neurons shown in **Figure 5a** could project to a second set of VP neurons that functionally take the place of the VTA GABA interneurons (**Supplementary Figure 9a,c,f**), leading to sustained positive value encoding VP neurons as were recently observed in VTA-projecting VP neurons (Tian et al., 2016). Additionally, the convergence of the right-choice and left-choice neurons could occur in VP rather than VTA, so that there is no explicit encoding of *V_L_* and *V_R_* (**Supplementary Figure 9a**).

We next asked how the same model using single-trial activity from choice-selective mTH-NAc neurons would perform (**Figure 5h**). In line with the less consistent sequential choice-selective activity in mTH-NAc relative to PL-NAc (**Supplementary Figure 7**; compare with **Figure 4**), we observed a substantial reduction in performance when using mTH-NAc activity as input (**Figure 5i-k**). In fact, the mTH-NAc model performed at chance levels (39.7% reward rate, compared with chance reward rate of 40%). Additionally, relative to the PL-NAc model, using mTH-NAc activity resulted in a reduction in the negative modulation of dopamine signal following an unrewarded outcome (**Figure 5l, top**; compare with **Figure 5g, top**), and less effect of previous trial outcome on dopamine response (**Figure 5l, bottom**; compare with **Figure 5g, bottom**). Compared with PL-NAc, using mTH-NAc as input also resulted in a reduction in the speed at which the correct value is learned within the NAc and VTA GABA neurons (**Supplementary Figure 11c,d**; compare with **Supplementary Figure 11a,b**).

The choice-selective sequences in PL-NAc neurons were critical to model performance, as they allowed the backpropagation of the RPE signal across trials. This was verified by comparing performance to a model in which PL-NAc activity was shifted to be synchronously active at the trial onset (**Figure 5m**). Unlike the sequential data, synchronous choice-selective PL-NAc activity was unable to correctly modulate lever value following block reversals, and therefore did not lead to correct choices, resulting in model performance near chance (**Figure 5n-p**; synchronous model reward rate: 39.8%, compared with chance reward rate of 40%). This was due to the fact that the synchronous model was unable to generate an RPE signal that was modulated properly within or across trials. Within a trial, negative outcomes did not result in the expected decrease in the dopamine signal (**Figure 5q, top**), while across trials, the influence of the previous trial’s outcome on the dopamine signal was disrupted relative to both the sequential model and recorded dopamine activity (**Figure 5q, bottom**, compare to **Figure 5g** and recorded dopamine activity in NAc in **Supplementary Figure 10**). Without a properly calculated RPE signal, the synchronous model was unable to generate value signals that correlated with the identity of the high probability lever in either NAc or the VTA GABA interneuron (**Supplementary Figure 11e,f**).

### Stimulation of PL-NAc (but not mTH-NAc) neurons decreases the effect of previous trial outcomes on subsequent choice in both the model and the mice

We next sought to generate experimentally testable predictions from our synaptic plasticity model by examining the effect of disruption of these sequences on behavioral performance. Towards this end, we simulated optogenetic-like neural stimulation of this projection by replacing the PL-NAc sequential activity in the model with constant, population-wide and choice-independent activity across the population on a subset of trials (on 10% of trials, 65% of neurons were stimulated; **Figure 6a,b**). This generated a decrease in the probability of staying with the previously chosen lever following rewarded trials and an increase following unrewarded trials relative to unstimulated trials (**Figure 6c**). In other words, the effect of previous outcome on choice was reduced when PL-NAc activity was disrupted. This effect persists for multiple trials, as revealed by including terms that account for the interaction between previous rewarded and unrewarded choices and stimulation in the logistic choice regression introduced in **Figure 1e** (**Figure 6d**; see Methods for details). This occurs because stimulation disrupts the choice-selectivity that is observed in the recorded PL-NAc sequences, thus allowing dopamine to indiscriminately adjust the synaptic weights (i.e. value) of *both* the right and left PL-NAc synapses following rewarded or unrewarded outcomes. The effect of stimulation is observed multiple trials back because the incorrect weight changes persist for multiple trials in the synaptic plasticity model. In contrast, stimulation did not result in a difference in stay probability on the trial with stimulation, as choice was determined by spontaneous activity before the sequence initiates in PL-NAc neurons, combined with the synaptic weights between PL and NAc neurons (**Figure 6c**).

**Figure 6.**
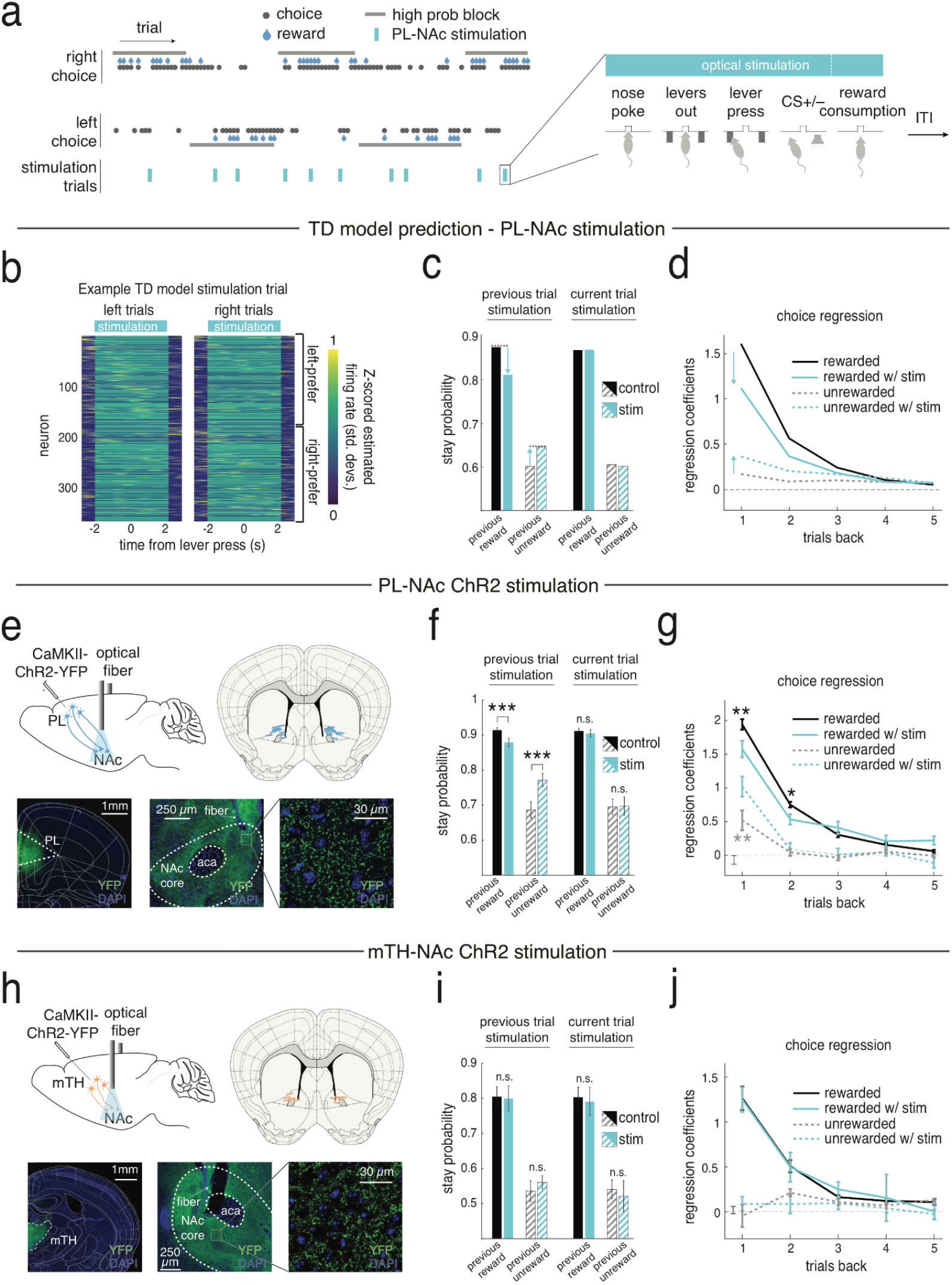
Stimulation of PL-NAc neurons disrupts the influence of previous trial outcomes on subsequent choice in both the synaptic plasticity model and in mice. **(a)** In the synaptic plasticity model and in mice, PL-NAc neurons were stimulated for the full duration of the trial on a randomly selected 10% of trials, disrupting the endogenous choice-selective sequential activity. For the experimental data, ChR2 stimulation (447 nm, 20 Hz stimulation, 5 ms pulse duration, 1-3 mW) began when the mouse entered the central nose poke and continued until either 2s after the time of the CS or the end of reward consumption, depending on the outcome of the trial and the experimental cohort (see Methods and **Supplementary Figure 13**). **(b)** Heatmap representing single-trial activity of PL-NAc input to the synaptic plasticity model on a stimulation trial. Optogenetic-like stimulation begins at the simulated time of nose poke and ends 2s after CS presentation. **(c)** Effect of stimulating the PL-NAc input in the synaptic plasticity model. Left, stimulation on the previous trial reduced the stay probability following a stimulation+rewarded trial but increased the stay probability following a stimulation+unrewarded trial. Right, in contrast to the effect of previous trial stimulation, stimulation on the current trial had no effect on the model’s stay probability following either rewarded or unrewarded trials. **(d)** Logistic choice regression showing dependence of the current choice on previously rewarded choices, and unrewarded choices, with and without stimulation (see Methods). Higher coefficients indicate a higher probability of staying with the previously chosen lever. The model predicts that previous trial stimulation will decrease the stay probability of rewarded trials but increase the stay probability of unrewarded trials across multiple previous trials. **(e)** Top left, surgical schematic illustrating injection site of an AAV2/5 expressing CaMKII-ChR2-eYFP in the PL (black needle) and optical fiber implant in the NAc core terminals. Top right, location of optical fiber tips of PL-NAc ChR2 cohort. Bottom left, coronal section showing ChR2-YFP expression in PL. Bottom middle, ChR2-YFP terminal expression in the NAc-core. Bottom right, ChR2-eYFP expression in PL terminals in the NAc core. **(f)** Left, as predicted by the model (panel **c**), PL-NAc ChR2 stimulation on the previous trial significantly reduced the mice’s stay probability following a rewarded trial (P = 0.002: paired, two-tailed t-test across mice, n=14) while increasing the mice’s stay probability following an unrewarded trial (P = 0.0005: paired, two-tailed t-test across mice, n=14). Right, also consistent with the model prediction, stimulation on the current trial had no significant effect on stay probability following rewarded (P = 0.62: paired, two-tailed t-test across mice, n=14) or unrewarded (P=0.91: paired, two-tailed t-test across mice, n=14) trials. **(g)** Same as **d**, but for PL-NAc stimulation in mice instead of synaptic plasticity model. Similar to the model, PL-NAc ChR2 stimulation decreased the weight of rewarded choices one- and two-trials back (P=0.002: one-trial back; P=0.023: two-trial back: one-sample, two-tailed t-test across mice, n=14) and increased the weight of unrewarded choices one-trial back (P=5.4×10^-6^, one-sample, two-tailed t-test across mice, n=14). **(h-j)** Same as **e-g** but for mTH-NAc ChR2 stimulation. Unlike PL-NAc ChR2 stimulation, mTH-NAc stimulation had no significant effect on stay probability following either rewarded or unrewarded choices on the previous trial (**i**, P>0.05, paired t-test across mice) and multiple trials back (**j**, P>0.05 for all trials back, t-test across laser x choice interaction term coefficients)

We tested these model predictions experimentally by performing an analogous manipulation in mice, which involved stimulating PL-NAc axon terminals with ChR2 on 10% of trials (**Figure 6e**). In close agreement with our model, mice had a significant decrease in their stay probability following a rewarded trial that was paired with stimulation (**Figure 6f**; P=0.001: paired, two-tailed t-test across mice, n=14, comparison between stay probability following rewarded trials with and without stimulation), while they were more likely to stay following an unrewarded trial paired with stimulation (**Figure 6f**; P=0.0005: paired, two-tailed t-test across mice, n=14, comparison between stay probability following unrewarded trials with and without stimulation). Similar to the synaptic plasticity model (**Figure 6d**), the effect of stimulation on the mouse’s choice persisted for multiple trials. Mice had a significant decrease in their stay probability following PL-NAc stimulation on rewarded choices one and two trials back (**Figure 6g**; P=0.001 for one trial back; P=0.02 for two trials back; one-sample, two-tailed t-test across mice of regression coefficients corresponding to the interaction term between rewarded choice and optical stimulation, n=14 mice).

Also similar to the model, and in contrast to these effects on the next trial, stimulation on the current trial had no significant effect on choice following either rewarded or unrewarded trials (**Figure 6f**; P>0.5: paired, two-tailed t-test across mice, n=14, comparison between stay probability on trials with and without stimulation following both rewarded and unrewarded trials).

We also observed an increase in the probability of mice abandoning the trials with stimulation compared with those trials without (P=0.0006: paired, two-tailed t-test comparing percentage of abandoned trials on stimulated versus non-stimulated trials; 12.2+/-2.5% for stimulated trials, 0.9+/-0.2% for non-stimulated trials). Relatedly, we found an increase in the latency to initiate a trial following PL-NAc stimulation (**Supplementary Figure 12a,b**). Together, these results suggest that this manipulation had some influence on the mouse’s motivation to perform the task.

Given the relatively weak choice encoding in mTH-NAc compared to PL-NAc (**Figure 3a,b**), and the fact that the mTH-NAc did not support effective trial-by-trial learning in our model (**Figure 5h-k**), we hypothesized that optogenetic stimulation of the mTH-NAc projection might not impact choice (**Figure 6h**). Indeed, in contrast to PL-NAc stimulation, mTH-NAc stimulation had no significant effect on the mice’s stay probability on the subsequent trial, following either rewarded or unrewarded stimulation trials (**Figure 6i**; P=0.84: paired, two-tailed t-test across mice, n=8, comparison of stay probability following rewarded trials with and without stimulation; P=0.40: paired two-tailed t-test, comparison of stay probability following unrewarded trials with and without stimulation). Similarly, inclusion of mTH-NAc stimulation into our choice regression model from **Figure 1e** revealed no significant effect of stimulation on rewarded or unrewarded choices (**Figure 6j**; P>0.05 for all trials back: one-sample, two-tailed t-test of regression coefficients corresponding to the interaction term between rewarded or unrewarded choice and optical stimulation across mice, n=8 mice). Additionally, there was no effect on the mice’s stay probability for current trial stimulation (**Figure 6i**; P=0.59: paired, two-tailed t-test across mice, n=8, comparison of stay probability following rewarded trials with and without stimulation; P=0.50: paired, two-tailed t-test, comparison of stay probability following unrewarded trials with and without stimulation). Similar to PL-NAc stimulation, mTH-NAc stimulation generated an increase in the probability of abandoning a trial on stimulation trials compared with control trials (P=0.032: paired, two-tailed t-test comparing percentage of abandoned trials on stimulated versus non-stimulated trials; 22.1+/-7.9% for stimulated trials, 6.4+/-3.1% for non-stimulated trials) as well as an increase in the latency to initiate the next trial (**Supplementary Figure 12c**), indicating that laser stimulation of the mTH-NAc projection may affect motivation to perform the task while not affecting the mouse’s choice.

To control for non-specific effects of optogenetic stimulation, we ran a control cohort of mice that received identical stimulation but did not express the opsin (**Supplementary Figure 12e,f**). Stimulation had no effect on the mice’s choice behavior (**Supplementary Figure 12g,h**) nor on the probability of abandoning trials on stimulation versus control trials (P=0.38: paired, two-tailed t-test comparing percentage of abandoned trials on stimulated versus non-stimulated trials; 0.4+/-0.08% for stimulated trials, 0.4+/-0.01% for non-stimulated trials).

### Trial-by-trial modulation in neural dynamics, without changes in synaptic plasticity on the timescale of trials, provides an alternative explanation for the behavioral and neural results

In our synaptic plasticity model (**Figure 5**), plasticity in PL-NAc synapses was used to produce value correlates in the NAc, and the values are compared across the choices for action selection. Thus, this model requires fast, dopamine-mediated synaptic plasticity, on the time scale of a trial, to mediate behavior. Whether plasticity operates in the NAc on this timescale is unclear. An alternative hypothesis, which preserves the same conceptual structure of this model but does not require fast synaptic plasticity, is that the across-trial updating of values and corresponding selection of actions is accomplished through the dynamics of a recurrent neural network rather than the dynamics of synaptic plasticity (Botvinick et al., 2019, 2020; Doshi-Velez and Konidaris, 2016; Song et al., 2017; Wang et al., 2018). The initial learning of the neural network’s synaptic weights is based on a reinforcement learning algorithm, which models slow initial task acquisition, but during task performance, synaptic weights remain fixed and the DA RPE serves only to alter neural dynamics.

We thus developed such a neural dynamics model based on the recorded PL-NAc activity (**Figure 7a**; see Methods for model details). Similar to the synaptic plasticity model, single-trial, experimentally recorded PL-NAc activity was input to a recurrent neural network that modeled NAc and other associated brain regions (the “critic network”) to calculate value. Next, the TD RPE was calculated in the DA neurons from the value signal using the same circuit architecture as the synaptic plasticity model. However, rather than reweighting PL-NAc synapses on the timescale of trials, the TD RPE was input to a second recurrent neural network that modeled dorsomedial striatum (DMS) and other associated brain regions (the “actor network”; Atallah et al., 2007; Lau and Glimcher, 2008; O’Doherty et al., 2004; Richard et al., 2016; Tsutsui et al., 2016). This actor network, which is also a recurrent neural network, used this TD RPE input from the previous timestep, along with the action from the previous timestep and a “temporal context” sequence which may arise from hippocampus or other cortical areas (Howard and Eichenbaum, 2013; Leon and Shadlen, 2003), in order to select the choice (left choice, right choice, or no action). This choice, in turn, determined which PL-NAc choice sequence was initiated.

**Figure 7.**
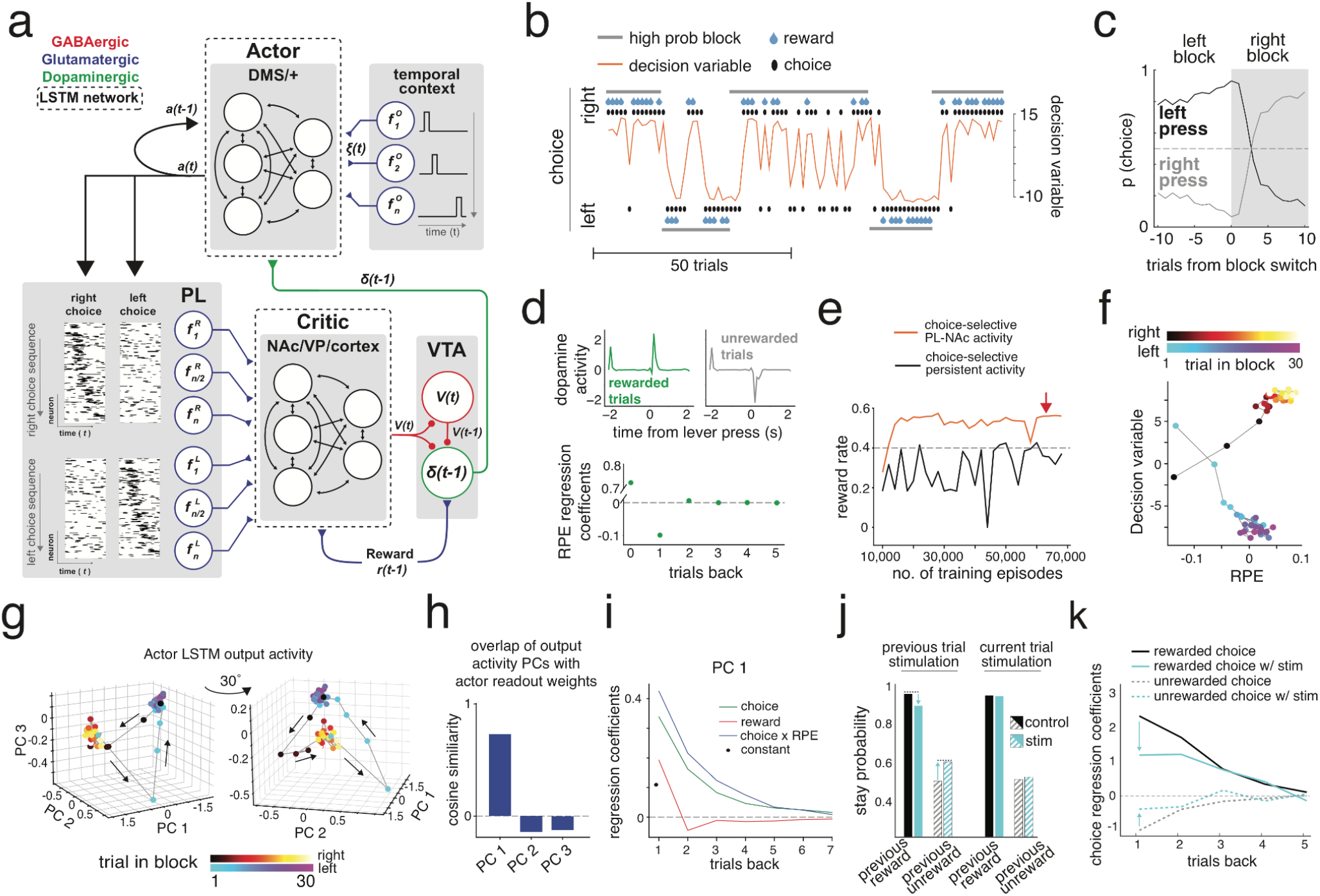
Neural dynamics model, which receives recorded choice-selective PL-NAc activity as input to the critic, performs the task and replicates the experimentally observed effects of PL-NAc stimulation, similarly to synaptic plasticity model. **(a)** Model schematic. The critic is a recurrent neural network which receives choice-selective sequential activity from recorded single trial activity of PL-NAc neurons (see **Figure 5c**) as well as reward input from the previous time step, *r*(*t* – 1). Value output from the critic network, *V*(*t*), is used to compute a temporal difference reward prediction error (RPE) signal, *δ*(*t* – 1), using a temporal delay and sign inversion provided by a GABAergic VTA interneuron, similar to the synaptic plasticity model in **Figure 5a**. An actor LSTM network within the dorsomedial striatum (DMS) and other associated brain regions (+) receives *δ*(*t* – 1) along with a temporal context signal, *ξ*(*t*), and the action on the previous time step *a*(*t* – 1). The output of the actor network is then passed through a logit function to compute the action at the current time step, *a*(*t*). The weights of LSTM units within the critic and actor networks are learned during the training stage and then frozen for the testing phase. All results shown are from the testing phase where all synaptic weight values are frozen and thus the modulation in model behavior shown are a result of changes in network dynamics alone. See Methods for additional details. **(b)** Example behavior from the neural dynamics model. Similar to the synaptic plasticity model in **Figure 5**, the decision variable (orange) and choice (black dot) of the model follows the identity of the high probability lever (grey bars). The decision variable is defined as the difference in the log odds (logit) of choosing right versus choosing left at the time of decision. **(c)** Choice probability of the neural dynamics model around the time of a block reversal. The model is able to rapidly modulate its choice following a block reversal. **(d)** Similar to the synaptic plasticity model, dopamine activity in the neural dynamics model encodes a quantitative reward prediction error signal. Top, dopamine activity displays a transient increase or decrease in activity on rewarded and unrewarded trials, respectively. Bottom, previous trial outcome modulates the magnitude of dopamine response to reward in a manner consistent with a reward prediction error signal. **(e)** Reward rate as a function of the number of training episodes for the model with recorded PL-NAc input to the critic (red) and an alternative model with persistent choice-selective input (instead of PL-NAc input) to the critic (black). Red arrow corresponds to the training duration of the neural dynamics model with PL-NAc input used in generating all other panels. The model with sequential PL-NAc input was able to learn the task whereas the model with persistent choice-selective activity failed to learn the task, achieving a reward rate near chance (0.4, grey dashed trace). **(f)** Relationship between the decision variable used to select the choice on the next trial and the calculated RPE across right and left blocks. The actor responds oppositely to similar RPE inputs in the two blocks - causing an increase in the probability of selecting the right lever for the right block while causing a decrease in the probability of choosing the right lever in the left block. The change in probability occurs within the first few trials of a block, corresponding to the rapid choice reversal observed in panels **b-c**. The RPE signal shown is calculated as an average over all time points past the time of lever press (0-2s) and is averaged across blocks. The decision variable is also averaged across blocks. **(g)** Evolution of the principal components of the output of the actor LSTM units across trials within a right and left block. The network activity shown is taken from the first time point in a trial (where the choice decision is made) and is averaged across all blocks. The network neural dynamics at this time point exhibits separate representations for the two blocks, with network activity changing rapidly at block reversals before settling into an approximate steady state. The first three components (out of 128 total outputs) accounted for 71.0%, 16.5%, and 6.6% of the total variance, respectively. **(h)** Cosine of the angle between the actor network’s readout weight vector and the vectors corresponding to the three principal components. Network activity in the PC1 direction (but not PC2 or PC3) aligns with the network readout weights. **(i)** Coefficients from a linear regression that uses choice and RPE information from the previous 7 trials to predict the value of PC1 on the current trial. The regression used three sets of predictors: (i) ‘Choice’ (green), which identifies a previous trial as a right press (+1) or a left press (−1), (ii) ‘RPE’ (red), which is given by the average RPE over all time points following the time of lever press (0-2s) on that trial and (iii) ‘Choice x RPE’ which is given by a product of the ‘Choice’ and ‘RPE’ predictors. PC1 is modulated by both choice and RPE history multiple trials back. **(j)** Stimulation of PL-NAc input to the neural dynamics model reproduces the reduction in stay probability following rewarded trials and increase in stay probability following unrewarded trials observed with optogenetic manipulations (**Figure 6f**). **(k)** Similar to optogenetic experiments (**Figure 6g**), PL-NAc stimulation in the neural dynamics model modulates the effect of previously rewarded choices multiple trials back.

Similar to the synaptic plasticity model (**Figure 5d,e**), our neural dynamics model was able to rapidly modulate choice following a reversal in the identity of the high probability lever (**Figure 7b,c**). In addition, the neural dynamics model generated RPE signals in VTA dopamine neurons that resemble previous experimental recordings (**Figure 7d**; **Supplementary Figure 10**). By contrast, when we replaced the choice-selective sequences to the NAc by choice-selective persistent activity the model failed to train within the same number of training episodes (**Figure 7e**). This suggests that temporal structure in this input is beneficial for efficient task learning.

Given the symmetry in the block structure, the average RPE signal will be statistically similar as a function of trial number within each block. However, the model must make different choices across blocks to achieve high performance, given that the choice that is most likely to lead to reward changes across blocks. Therefore, across blocks, the actor network needs to respond differently to similar RPE inputs. Consistent with this, the decision variable for a given RPE was different for left versus right blocks (**Figure 7f**).

To further understand how the model is able to appropriately modulate its choices, we analyzed the evolution of the actor network’s activity across trials. Towards this, we performed dimensionality reduction on the actor network’s activity at the first time step in a trial (i.e., the time of decision) using principal components analysis. The actor network’s dynamics were low-dimensional, with the first three principal components explaining ~95% of the variance. The dynamics were different for the two blocks (**Figure 7g**), consistent with the model responding differently to similar RPE inputs across blocks (**Figure 7f**). At a block reversal, for example from a left block to a right block, the network activity rapidly transitioned from the approximately steady-state representation of the left block (cluster of blue-purple points in **Figure 7g**) to the approximately steady-state representation of the right block (cluster of red-yellow points). Furthermore, the model learned to align the first principal component of activity along the direction of the network readout weights that determine the actor’s choice *a*(*t*) (**Figure 7h**). Thus, the actor learned to generate an explicit representation of the decision variable in the first principal component of its activity.

To solve the reversal learning task, the network needs to use its past history of choices and rewards to accumulate evidence for whether the current block is a left block or a right block. Rewarded left-side choices, or unrewarded right-side choices, represent evidence that the current block is a left block, while the converse represents evidence for a right block. In the synaptic plasticity model (**Figure 5b**), this accumulation occurs in the PL-NAc synaptic weights, which accumulate the product of RPE (representing whether a choice was rewarded) and the eligibility trace (which, due to the choice-selectivity of the PL-NAc activity, represents the current choice). To analyze whether the actor network uses a similar accumulation of evidence to solve the task, we linearly regressed the first principal component of actor activity (PC1, which correlated strongly with the decision variable as described above) against the past history of choices and RPEs, which serve as inputs to the network, as well as the product of these (‘Choice x RPE’), which represents evidence. PC1 most strongly depended upon the ‘Choice x RPE’ predictor, with coefficients that decayed on a timescale of approximately 3 trials, suggesting that the actor used a leaky accumulation of evidence over this timescale to solve the task (**Figure 7i**). In addition, like the mice and the synaptic plasticity model, the neural dynamics model tended to stay with its previous choices as evident from the positive coefficients corresponding to the previous choice regressors in **Figure 7i**. Therefore, both the synaptic plasticity model and the neural dynamics model follow the same principle of accumulating evidence across trials to perform fast reversal learning in addition to having a tendency to repeat their previous choices.

To determine if the neural dynamics model, similar to the synaptic plasticity model, is able to recapitulate our optogenetic experiments, we simulated optogenetic stimulation of a fraction of PL-NAc neurons in the same manner as described for the synaptic plasticity model. Like the synaptic plasticity model and optogenetic experiments, relative to unstimulated trials, stimulation decreased the probability of staying with the previously chosen lever following rewarded trials and increased the probability following unrewarded trials (**Figure 7j**). Also like the synaptic plasticity model, this effect persisted for multiple trials (**Figure 7k**). In the case of the neural dynamics model, this disruption was due to a cascade of events: PL-NAc stimulation disrupted the activity of the critic (NAc and associated regions), thus disrupting the RPE signal that is transmitted to the actor. This disrupted RPE then leads to the observed disruption in behavior. Thus, both the synaptic plasticity and neural dynamics models exhibit disrupted trial-by-trial learning when PL-NAc activity is stimulated in a manner mimicking our experimental optogenetic perturbations.

## Discussion

This work provides both experimental and computational insights into how the NAc and associated regions could contribute to reinforcement learning. Experimentally, we found that mTH-NAc neurons were preferentially modulated by a reward-predictive cue, while PL-NAc neurons more strongly encoded actions (e.g. nose poke, lever press). In addition, PL-NAc neurons display choice-selective sequential activity which persists for several seconds after the lever press action, beyond the time the mice receive reward feedback. Computationally, we demonstrate that the choice-selective and sequential nature of PL-NAc activity can contribute to performance of a choice task by implementing a circuit-based version of reinforcement learning (Sutton and Barto, 1998; Tesauro, 1992), based on either synaptic plasticity or neural dynamics. The models are able to i) perform the task, ii) replicate previous recordings in VTA dopamine and GABA neurons, and iii) make new predictions that we have experimentally tested regarding the effect of perturbing PL-NAc or mTH-NAc activity on trial-by-trial learning. Thus, this work suggests a computational role of choice-selective sequences, a form of neural dynamics whose ubiquity is being increasingly appreciated (Kawai et al., 2015; Kim et al., 2017; Long et al., 2010; Ölveczky et al., 2011; Picardo et al., 2016; Sakata et al., 2008).

### Relationship to previous neural recordings in the NAc and associated regions

To our knowledge, a direct comparison, at cellular resolution, of activity across multiple glutamatergic inputs to the NAc has not previously been conducted. This is a significant gap, given that these inputs are thought to contribute critically to reinforcement learning by providing the information to the NAc that dopaminergic inputs can modulate (Centonze et al., 2001; Nestler, 2001; Nicola et al., 2000; Shen et al., 2008; Wilson, 2004; Xiong et al., 2015). The differences in the representations in the two populations that we report were apparent due to the use of a behavioral task with both actions and sensory stimuli.

The preferential representations of actions relative to sensory stimuli in PL-NAc is somewhat surprising, given that previous studies have focused on sensory representations in this projection (Otis et al., 2017), and also given that the NAc is heavily implicated in Pavlovian conditioning (Day and Carelli, 2007; Day et al., 2006; Di Ciano et al., 2001; Parkinson et al., 1999; Roitman et al., 2005; Wan and Peoples, 2006). On the other hand, there is extensive previous evidence of action correlates in PFC (Cameron et al., 2019; Genovesio et al., 2006; Luk and Wallis, 2013; Siniscalchi et al., 2019; Sul et al., 2010), and NAc is implicated in operant conditioning in addition to Pavlovian conditioning (Atallah et al., 2007; Cardinal and Cheung, 2005; Collins et al., 2019; Hernandez et al., 2002; Kelley et al., 1997; Kim et al., 2009; Salamone et al., 1991).

Our finding of sustained choice-encoding in PL-NAc neurons is in agreement with previous work recording from medial prefrontal cortex (mPFC) neurons during a different reinforcement learning task (Maggi and Humphries, 2019; Maggi et al., 2018). Additionally, other papers have reported choice-selective sequences in other regions of cortex, as well as in the hippocampus (Harvey et al., 2012; Pastalkova et al., 2008; Terada et al., 2017). In fact, given previous reports of choice-selective (or outcome-selective) sequences in multiple brain regions and species (Kawai et al., 2015; Kim et al., 2017; Long et al., 2010; Ölveczky et al., 2011; Picardo et al., 2016; Sakata et al., 2008), the relative absence of sequences in mTH-NAc neurons may be more surprising than the presence in PL-NAc.

Our observation of prolonged representation of the CS+ in mTH-NAc (**Figure 2d,f**) is in line with previous observations of pronounced and prolonged encoding of task-related stimuli in the primate thalamus during a Pavlovian conditioning task (Matsumoto et al., 2001). Together with our data, this suggests that the thalamus is contributing information about task-relevant stimuli to the striatum, which could bridge the gap between a CS and US in a Pavlovian trace conditioning task (Campus et al., 2019; Do-Monte et al., 2017; Otis et al., 2019; Zhu et al., 2018). Consistent with the observation that mTH-NAc displayed weaker choice representation relative to PL-NAc, disruption of this mTH projection did not have a causal effect on our choice task **(Figure 3a,b; Figure 6h-j**). We also demonstrate that mTH-NAc stimulation results in both an increase in the number of abandoned trials as well as the latency to initiate the subsequent trial (**Supplementary Figure 12c**), suggesting that this population may play a role in motivation as opposed to action selection. This is in line with recent work demonstrating a role of thalamostriatal projections in the initiation and execution, but not learning of a motor task (Wolff et al., 2019).

### Choice-selective sequences implement TD learning in a neural circuit synaptic plasticity model of task performance

Given the widespread observation of choice-selective sequences across multiple behaviors and brain regions, including the PL-NAc neurons that we record from in this study, a fundamental question is what is the computational function of such sequences. Here, we suggest that these sequences may contribute to the neural implementation of TD learning, by providing a temporal basis set that bridges the gap in time between actions and outcomes. Specifically, neurons active in a sequence that ultimately results in reward enable the backpropagation in time of the dopaminergic RPE signal, due to the fact that the earlier neurons in the sequence predict the activity of the later neurons in the sequence, which themselves overlap with reward. This causes synaptic weights onto NAc of these earlier neurons to be strengthened, which in turn biases the choice towards that represented by those neurons.

While TD learning based on a stimulus representation of sequentially active neurons has previously been proposed for learning in the context of sequential behaviors (Fee and Goldberg, 2011; Jin et al., 2009), and for learning the timing of a CS-US relationship (Aggarwal et al., 2012; Carrillo-Reid et al., 2008; Gershman et al., 2014; Ponzi and Wickens, 2010), here we extend these ideas in several important ways:

First, we link these theoretical ideas directly to data, by demonstrating that choice-selective sequential activity in the NAc is provided primarily by PL-NAc (as opposed to mTH-NAc) input neurons, and that perturbation of the PL-NAc (but not mTH-NAc) projection disrupts action-outcome pairing consistent with model predictions. Specifically, in our synaptic plasticity model, overwriting the choice-selective sequential activity with full trial optogenetic-like stimulation of PL-NAc neurons disrupts the model’s ability to link choice and outcome on one trial so as to guide choice on the subsequent trials (**Figure 6b-d**). In contrast to the effect of stimulation on subsequent trials’ choice, stimulation caused no disruption of choice on the stimulated trial, in either the synaptic plasticity model or the experimental results (**Figure 6c,f**). This is true in the synaptic plasticity model because choice is determined by the PL-NAc weights at the beginning of the trial, which are determined by previous trials’ choices and outcomes. Thus, the synaptic plasticity model provides a mechanistic explanation of a puzzling experimental finding: that optogenetic manipulation of PL-NAc neurons affects subsequent choices but not the choice on the stimulation trial itself and that this stimulation creates oppositely directed effects following rewarded versus unrewarded trials.

Second, we extend these ideas to the performance of a choice task. For appropriate learning in a choice task, an RPE must update the value of the chosen action only. Given that dopamine neurons diffusely project to striatum, and given that the dopamine RPE signal is not action-specific (Lee et al., 2019), it is not immediately obvious how such specificity would be achieved. In our synaptic plasticity model, the specificity of the value updating by dopamine to the chosen action is possible because the PL-NAc neurons that are active for one choice are silent for the other choice. Similarly, even though dopamine neurons receive convergent inputs from both NAc/VP neurons that encode right-side value as well as others that encode left-side value (given, once again, that dopamine RPE signals are not action-specific), the model produces dopamine neuron activity that is dependent on the value of the chosen action (rather than the sum of left and right action values) by virtue of the fact that PL-NAc inputs that are active for one choice are silent for the other.

Third, our synaptic plasticity model replicates numerous experimental findings in the circuitry downstream of PL-NAc. The most obvious is the calculation of an RPE signal in dopamine neurons (Bayer and Glimcher, 2005; Parker et al., 2016), which allows proper value estimation and, thus, task performance. We had previously confirmed an RPE signal in NAc-projecting dopamine neurons in this task (Parker et al., 2016). In addition, GABA interneurons encode value (Cohen et al., 2012; Tian et al., 2016), as predicted by our model, and their activation inhibits dopamine neurons, again consistent with our model (Eshel et al., 2015). This value representation in the GABA interneurons is key to our model producing an RPE signal in dopamine neurons, as it produces the temporally delayed, sign inverted signals required for the calculation of a temporally differenced RPE (**Figure 5a**). Although previous work had suggested other neural architectures for this temporal differencing operation (Aggarwal et al., 2012; Carrillo-Reid et al., 2008; Doya, 2002; Hazy et al., 2010; Ito and Doya, 2015; Joel et al., 2002; Pan et al., 2005; Suri and Schultz, 1998, 1999), these models have not been revisited in light of recent cell-type and projection-specific recordings in the circuit. Consistent with our synaptic plasticity model, electrical stimulation of VP generates both immediate inhibition of dopamine neurons, and delayed excitation, as required by our model (Chen et al., 2019).

Our specific proposal for temporal differencing by the VTA GABA interneuron is attractive in that it could provide a *generalizable* mechanism for calculating RPE: it could extend to any input that projects both to the dopamine and GABA neurons in the VTA, and that also receives a dopaminergic input that can modify synaptic weights.

### Limitations of the synaptic plasticity model and trial-by-trial changes in neural dynamics as an alternative model of the neural implementation of reinforcement learning

While the synaptic plasticity model replicates multiple features of neurons throughout the circuit, it does not replicate some findings on the relative timing and sign of the value correlates in the NAc versus the VP (Chen et al., 2019; Ottenheimer et al., 2018; Tian et al., 2016), likely because of anatomical connections that we omitted from our circuit for simplicity (e.g. VP→ NAc; Ottenheimer et al., 2018; Wei et al., 2016; also see **Supplementary Figure 9** for discussion of feasible model variants that produce a greater diversity of signals in VP). Similarly, our synaptic plasticity model does not produce previously observed responses to the conditioned stimuli in VP (Ottenheimer et al., 2019; Stephenson-Jones et al., 2020; Tian et al., 2016; Tindell et al., 2004), although inclusion of cue-responsive mTH-NAc inputs into our model would remedy this. Relatedly, our model does not take into account other glutamatergic inputs to the NAc, such as those from the basolateral amygdala and ventral hippocampus (Bagot et al., 2015; French and Totterdell, 2003). Further investigations into what information these inputs encode will be elucidating.

In the synaptic plasticity model, appropriate trial-and-error learning is mediated by a dopamine-dependent synaptic plasticity mechanism that operates on the timescale of trials (**Figure 5b**). There is extensive evidence to support a key role for dopamine in this task on this timescale. For example, we had previously demonstrated that dopamine neurons that project to the NAc encode an RPE signal in this task, and that inhibiting these neurons during single trials serves as a negative prediction error signal, consistent with a reinforcement learning mechanism (Parker et al., 2016). Similar evidence for dopamine serving as a reinforcement learning signal has been obtained in other related paradigms in both rodents (Hamid et al., 2016; Kwak et al., 2014; Lak et al., 2020) and human subjects (Rutledge et al., 2009). In addition, other studies have used pharmacology or ablation to implicate dopamine-receptor pathways and NMDA-receptors in NAc in reversal learning (Boulougouris et al., 2009; Ding et al., 2014; Izquierdo et al., 2006; Kruzich and Grandy, 2004; O’Neill and Brown, 2007; Taghzouti et al., 1985).

However, whether dopamine-mediated synaptic plasticity operates on a fast enough timescale to mediate plasticity during trial-and-error learning is not clear (but see Shonesy et al., 2020 and Young and Yang, 2005 for some evidence of fast dopamine-mediated synaptic plasticity in cortex and striatum). Furthermore, model-free TD learning cannot take advantage of additional task-structure information such as the reward probabilities within a block (Collins and Cockburn, 2020; Doll et al., 2012, but see Supplementary Figure 14 for challenges in identifying this ability within tasks like ours). For these reasons, we presented an alternative model based on trial-by-trial modulation by dopamine of neural dynamics instead of synaptic plasticity (Figure 7; see related work by Botvinick et al., 2019, 2020; Doshi-Velez and Konidaris, 2016; Nagabandi et al., 2018; Song et al., 2017; Wang et al., 2018; Sæmundsson et al. 2018; Finn et al. 2017; Duan et al. 2016; Rakelly et al. 2019). In this model, dopamine affected choices by modifying dynamics in a recurrent neural network, while synaptic weights were fixed after initial training of the weights with an RPE. Thus, in the neural dynamics model, the DA RPE played a similar role during initial learning of the task as it contributed during the task in our synaptic plasticity model. Similar to our synaptic plasticity model, the choice-selective sequences were input to the value-calculating network. Eliminating the temporal structure of this input while preserving the choice-selectivity made training of the network less efficient (Figure 7e), suggesting that temporal structure is beneficial in the calculation of value. Similar to our synaptic plasticity model, the neural dynamics model also replicates our key experimental observations. Both models also generate a value signal and an RPE signal using single-trial recorded PL-NAc activity and a reward signal as input. Additionally, in both models, the calculation of the reward prediction error signal is accomplished using similar VTA circuit architecture downstream of the value calculation (compare Figures 5a and 7a). Finally, we showed that, analogous to the accumulation of past history of rewarded choices by the synaptic weights in the synaptic plasticity model (Figure 5b), the neural dynamics model appears to accumulate a similar variable in the output dynamics of its actor network (Figure 7i). Given the similarity in outputs of these two models, future experiments will need to be designed and executed to test predictions that distinguish the synaptic plasticity and neural dynamics models.

## Methods

### Mice

46 male C57BL/6J mice from The Jackson Laboratory (strain 000664) were used for these experiments. Prior to surgery, mice were group-housed with 3-5 mice/cage. All mice were >6 weeks of age prior to surgery and/or behavioral training. To prevent mice from damaging the implant of cagemates, all mice used in imaging experiments were single housed post-surgery. All mice were kept on a 12-h on/ 12-h off light schedule. All experiments and surgeries were performed during the light off time. All experimental procedures and animal care was performed in accordance with the guidelines set forth by the National Institutes of Health and were approved by the Princeton University Institutional Animal Care and Use Committee.

### Probabilistic reversal learning task

Beginning three days prior to the first day of training, mice were placed on water restriction and given per diem water to maintain >80% original body weight throughout training. Mice performed the task in a 21 x 18 cm operant behavior box (MED associates, ENV-307W). A shaping protocol of three stages was used to enable training and discourage a bias from forming to the right or left lever. In all stages of training, the start of a trial was indicated by illumination of a central nose poke port. After completing a nose poke, the mouse was presented with both the right and left lever after a temporal delay drawn from a random distribution from 0 to 1s in 100ms intervals. The probability of reward of these two levers varied based on the stage of training (see below for details). After the mouse successfully pressed one of the two levers, both retracted and, after a temporal delay drawn from the same uniform distribution, the mice were presented with one of two auditory cues for 500ms indicating whether the mouse was rewarded (CS+, 5 kHz pure tone) or not rewarded (CS−, white noise). Concurrent with the CS+ presentation, the mouse was presented with 6μl of 10% sucrose reward in a dish located equidistantly between the two levers, just interior to the central nose poke. The start time of reward consumption was defined as the moment the mouse first made contact with the central reward port spout following the delivery of the reward. The end of the reward consumption period (i.e. reward exit) was defined as the first moment at which the mouse was disengaged from the reward port for >100ms. In all stages of training, trials were separated by a 2s intertrial interval, which began either at the end of CS on unrewarded trials or at the end of reward consumption on rewarded trials.

In the first stage of training (“100-100 debias”), during a two hour session, mice could make a central nose poke and be presented with both the right and left levers, each with a 100% probability of reward. However, to ensure that mice did not form a bias during this stage, after five successive presses of either lever the mouse was required to press the opposite lever to receive a reward. In this case, a single successful switch to the opposite lever returned both levers to a rewarded state. Once a mouse received >100 rewards in a single session they were moved to the second stage (“100-0”) where only one of the two levers would result in a reward. The identity of the rewarded lever reversed after 10 rewarded trials plus a random number of trials drawn from the geometric distribution:

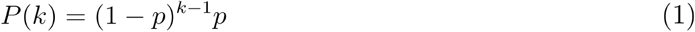

where *P*(*k*) is the probability of a block reversal k trials into a block and p is the success probability of a reversal for each trial, which in our case was 0.4. After 3 successive days of receiving >100 total rewards, the mice were moved to the final stage of training (“70-10”), during which on any given trial pressing one lever had a 70% probability of leading to reward (high-probability lever) while pressing the opposite lever had only a 10% reward probability (low-probability lever). The identity of the higher probability lever reversed using the same geometric distribution as the 100-0 training stage. On average, there were 23.23 +/- 7.93 trials per block and 9.67 +/- 3.66 blocks per session (mean +/- std. dev.). In this final stage, the mice were required to press either lever within 10s of their presentation; otherwise, the trial was considered an ‘abandoned trial’ and the levers retracted. All experimental data shown was collected while mice performed this final “70-10” stage.

### Logistic choice regression

For the logistic choice regressions shown in **Figure 1e** and **Supplementary Figure 2a**, we modeled the choice of the mouse on trial *i* based on lever choice and reward outcome information from the previous n trials using the following logistic regression model:

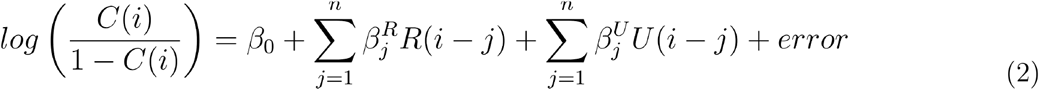

Where *C*(*i*) is the probability of choosing the right lever on trial *i*, and *R*(*i – J*) and *U*(*i – j*) are the choice of the mouse *j* trials back from the *i*-th trial for either rewarded or unrewarded trials, respectively. *R*(*i – j*) was defined as +1 when the *j*-th trial back was both rewarded and a right press, −1 when the *j*-th trial back was rewarded and a left press and 0 when it was unrewarded. Similarly, *U*(*i–j*) was defined as +1 when the *j*-th trial back was both unrewarded and a right press, −1 when the *j*-th trial back was unrewarded and a left press and 0 when it was rewarded. The calculated regression coefficients, 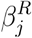 and 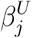, reflect the strength of the relationship between the identity of the chosen lever on a previously rewarded or unrewarded trial, respectively, and the lever chosen on the current trial.

To examine the effect of optogenetic stimulation from multiple trials back on the mouse’s choice (**Figure 6d,g,j; Figure 7k; Supplementary Figure 12h & Supplementary Figure 13c-d**), we expanded our behavioral logistic regression model to include the identity of those trials with optical stimulation, as well as the interaction between rewarded and unrewarded choice predictors and stimulation:

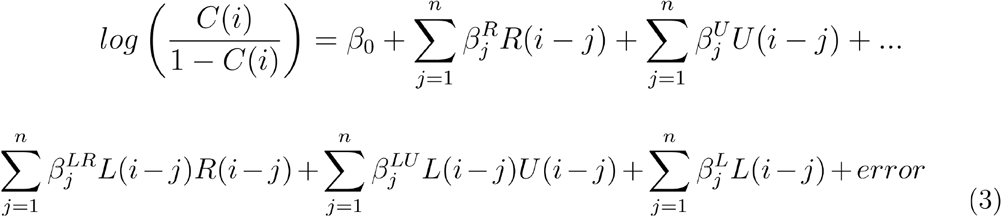

where *L*(*i*) represents optical stimulation on the *i*^th^ trial (1 for optical stimulation, 0 for control trials), 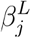 represents the coefficient corresponding to the effect of stimulation on choice j trials back and 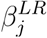 and 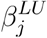 represent the coefficients corresponding to the interaction between rewarded choice x optical stimulation and unrewarded choice x stimulation, respectively.

To visualize the relative influence of stimulation on the mice’s choices compared with unstimulated trials, in **Figure 6d,g,j; Figure 7k; Supplementary Figure 12h & Supplementary Figure 13c-d**, the solid blue trace represents the sum of the rewarded choice coefficients (represented by the black trace) and rewarded choice x stimulation coefficients 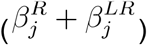. Similarly, the dashed blue trace represents the sum of the unrewarded choice coefficients (grey trace) and unrewarded choice x stimulation coefficients 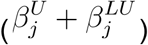. For all choice regressions, the coefficients for each mouse were fit using the *glmfit* function in MATLAB and error bars reflect means and s.e.m. across mice.

### Cellular-resolution calcium imaging

To selectively image from neurons which project to the NAc, we utilized a combinatorial virus strategy to image cortical and thalamic neurons which send projections to the NAc. 16 mice (7 PL-NAc, 9 mTH-NAc) previously trained on the probabilistic reversal learning task were unilaterally injected with 500nl of a retrogradely transporting virus to express Cre-recombinase (CAV2-cre, IGMM vector core, France, injected at ~2.5 × 10^12^ parts/ml or retroAAV-EF1a-Cre-WPRE-hGHpA, PNI vector core, injected at ~6.0 x 10^13^) in either the right or left NAc core (1.2 mm A/P, +/-1.0 mm M/L, −4.7 D/V) along with 600nl of a virus to express GCaMP6f in a Cre-dependent manner (AAV2/5-CAG-Flex-GCaMP6f-WPRE-SV40, UPenn vector core, injected at ~1.27 x 10^13^ parts/ml) in either the mTH (−0.3 & −0.8 A/P, +/- 0.4 M/L, −3.7 D/V) or PL (1.5 & 2.0 A/P, +/- 0.4 M/L, −2.5 D/V) of the same hemisphere. 154 of 278 (55%, n=5 mice) PL-NAc neurons and 95 out of 256 (37%, n=5 mice) mTH-NAc neurons were labeled using the CAV2-Cre virus, the remainder were labeled using the retroAAV-Cre virus. In this same surgery, mice were implanted with a 500 μ*m* diameter gradient refractive index (GRIN) lens (GLP-0561, Inscopix) in the same region as the GCaMP6f injection - either the PL (1.7 A/P, +/- 0.4 M/L, −2.35 D/V) or mTH (−0.5 A/P, +/- 0.3 M/L, −3.6 D/V). 2-3 weeks after this initial surgery, mice were implanted with a base plate attached to a miniature, head-mountable, one-photon microscope (nVISTA HD v2, Inscopix) above the top of the implanted lens at a distance which focused the field of view. All coordinates are relative to bregma using *Paxinos and Franklin’s the Mouse Brain in Stereotaxic Coordinates, 2nd edition* (Paxinos and Franklin, 2004). GRIN lens location was imaged using the Nanozoomer S60 Digital Slide Scanner (Hamamatsu) (location of implants shown in **Supplementary Figure 1**). The subsequent image of the coronal section determined to be the center of the lens implant was then aligned to the Allen Brain Atlas (Allen Institute, brain-map.org) using the *Wholebrain* software package (wholebrainsoftware.org; Fürth et al., 2018).

Post-surgery, mice with visible calcium transients were then retrained on the task while habituating to carrying a dummy microscope attached to the implanted baseplate. After the mice acclimated to the dummy microscope, they performed the task while images of the recording field of view were acquired at 10 Hz using the Mosaic acquisition software (Inscopix). To synchronize imaging data with behavioral events, pulses from the microscope and behavioral acquisition software were recorded using either a data acquisition card (USB-201, Measurement computing) or, when LED tracking (see below for details) was performed, an RZ5D BioAmp processor from Tucker-Davis Technologies. Acquired videos were then pre-processed using the Mosaic software and spatially downsampled by a factor of 4. Subsequent down-sampled videos then went through two rounds of motion-correction. First, rigid motion in the video was corrected using the translational motion correction algorithm based on Thévenaz et al. (1998) included in the Mosaic software (Inscopix, motion correction parameters: translation only, reference image: the mean image, speed/accuracy balance: 0.1, subtract spatial mean [r = 20 pixels], invert, and apply spatial mean [r = 5 pixels]). The video then went through multiple rounds of non-rigid motion correction using the NormCore motion correction algorithm (Pnevmatikakis and Giovannucci, 2017) NormCore parameters: gSig=7, gSiz=17, grid size and grid overlap ranged from 12-36 and 8-16 pixels, respectively, based on the individual motion of each video. Videos underwent multiple (no greater than 3) iterations of NormCore until non-rigid motion was no longer visible). Following motion correction, the CNMFe algorithm (Zhou et al., 2018) was used to extract the fluorescence traces (referred to as ‘GCaMP6f’ throughout the text) as well as an estimated firing rate of each neuron (CNMFe parameters: spatial downsample factor=1, temporal downsample=1, gaussian kernel width=4, maximum neuron diameter=20, tau decay=1, tau rise=0.1). Only those neurons with an estimated firing rate of four transients/ minute or higher were considered ‘task-active’ and included in this paper – 278/330 (84%; each mouse contributed 49,57,67,12,6,27,60 neurons, respectively) of neurons recorded from PL-NAc passed this threshold while 256/328 (78%; each mouse contributed 17,28,20,46,47,40,13,13,32 neurons, respectively) passed in mTH-NAc. Across all figures, to normalize the neural activity across different neurons and between mice, we Z-scored each GCaMP6f recording trace using the mean and standard deviation calculated using the entire recording session.

### Encoding model to generate response kernels for behavioral events

To determine the response of each neuron attributable to each of the events in our task, we used a multiple linear encoding model with lasso regularization to generate a response kernel for each behavioral event (example kernels shown in **Figure 2b**). In this model, the dependent variable was the GCaMP6f trace of each neuron recorded during a behavioral session and the independent variables were the times of each behavioral event (‘nose poke’, ‘levers out’, ‘ipsilateral lever press’, ‘contralateral lever press’, ‘CS+’, ‘CS−’ and ‘reward consumption) convolved with a 25 degrees-of-freedom spline basis set that spanned −2 to 6s before and after the time of action events (‘nose poke’, ‘ipsilateral press’, ‘contralateral press’ and ‘reward consumption’) and 0 to 8s from stimulus events (‘levers out’, ‘CS+’ and ‘CS−’). To generate this kernel, we used the following linear regression with lasso regularization using the *lasso* function in MATLAB:

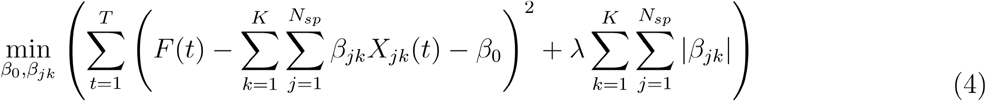

where *F*(*t*) is the Z-scored GCaMP6f fluorescence of a given neuron at time *t, T* is the total time of recording, *K* is the total number of behavioral events used in the model, *N_sp_* is the degrees-of-freedom for the spline basis set (25 in all cases), *β_jk_* is the regression coefficient for the *j^th^* spline basis function and *k^th^* behavioral event, *β*_0_ is the intercept term and λ is the lasso penalty coefficient. The value of lambda was chosen to minimize the mean squared error of the model, as determined by 5-fold cross validation. The predictors in our model, *X_jk_*, were generated by convolving the behavioral events with a spline basis set, to enable temporally delayed versions of the events to predict neural activity:

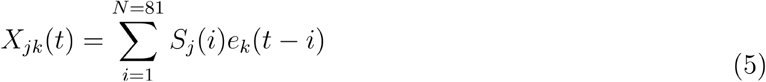

where *S_j_*(*i*) is the *j^th^* spline basis function at time point *i* with a length of 81 time bins (time window of −2 to 6s for action events or 0 to 8s for stimulus events sampled at 10 Hz) and *e_k_* is a binary vector of length *T* representing the time of each behavioral event *k* (1 at each time point where a behavioral event was recorded using the MED associates and TDT software, 0 at all other timepoints).

Using the regression coefficients, *β_jk_*, generated from the above model, we then calculated a ‘response kernel’ for each behavioral event:

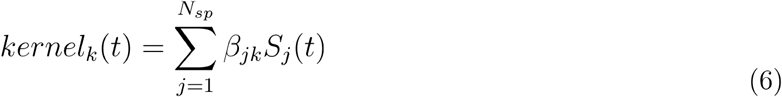

This kernel represents the (linear) response of a neuron to each behavioral event, while accounting for the linear component of the response of this neuron to the other events in the task.

### Quantification of neural modulation to behavioral events

To identify neurons that were significantly modulated by each of the behavioral events in our task (fractions shown in **Figure 2g-h**), we used the encoding model described above, but without the lasso regularization:

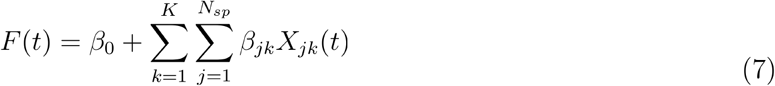

As above, *F*(*t*) is the Z-scored GCaMP6f fluorescence of a given neuron at time *t, K* is the total number of behavioral events used in the model, *N_sp_* is the degrees-of-freedom for the spline basis set (25 in all cases), *β_jk_* is the regression coefficient for the *j^th^* spline basis function and *k^th^* behavioral event and *β*_0_ is the intercept term. To determine the relative contribution of each behavioral event when predicting the activity of a neuron, we compared the full version of this model to a reduced model with the *X* and *β* terms associated with the behavioral event in question excluded. For each behavioral event, we first generated an F-statistic by comparing the fit of a full model containing all event predictors with that of a reduced model that lacks the predictors associated with the event in question. We then calculated this same statistic on 500 instances of shuffled data, where shuffling was performed by circularly shifting the GCaMP6f fluorescence by a random integer. We then compared the F-statistic from the real data to the shuffled distribution to determine whether the removal of an event as a predictor compromised the model significantly more than expected by chance. If the resulting P-value was less than the significance threshold of P=0.01, after accounting for multiple comparison testing of each of the behavioral events by Bonferroni correction, then the event was considered significantly encoded by that neuron.

To determine whether a neuron was significantly selective to the choice or outcome of a trial (‘choice-selective’ and ‘outcome-selective’, fractions of neurons from each population shown in **Figure 3a,c**), we utilized a nested model comparison test similar to that used to determine significant modulation of behavioral events above, where the full model used the following behavioral events as predictors: ‘nose poke’, ‘levers out’, ‘all lever press’, ‘ipsilateral lever press’, ‘all CS’, ‘CS+’ and ‘reward consumption’. For choice-selectivity, an F-statistic was computed for a reduced model lacking the ‘ipsilateral lever press’ predictors and significance was determined by comparing this value with a null distribution generated using shuffled data as described above. For outcome-selectivity, the reduced model used to test for significance lacked the predictors associated with both the ‘CS+’ and ‘reward consumption’ events.

By separating the lever press and outcome-related events into predictors that were either blind to the choice or outcome of the trial (‘all lever press’ and ‘all CS’, respectively) and those which included choice or outcome information (‘ipsilateral lever press’ or ‘CS+’ and ‘reward consumption’, respectively) we were able to determine whether the model was significantly impacted by the removal of either choice or outcome information. Therefore, neurons with significant encoding of the ‘ipsilateral lever press’ event (using the same P-value threshold determined by the shuffled distribution of F-statistics) were considered choice-selective, while those with significant encoding of the ‘CS+/reward consumption’ events were considered outcome-selective.

### Neural decoders

#### Choice decoder

In **Figure 3b**, we quantified how well simultaneously imaged populations of 1 to 10 PL-NAc or mTH-NAc neurons could be used to decode choice using a logistic regression:

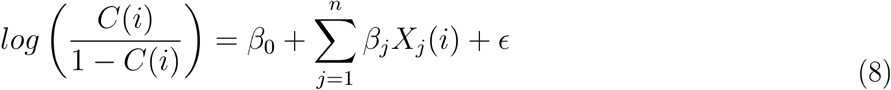

where C(i) is the probability the mouse made an ipsilateral choice on trial i, *β*_0_ is the offset term, n is the number of neurons (between 1 and 10), *β_j_* is the regression weight for each neuron, *X_j_*(*i*) is the mean z-scored GCaMP6f fluorescence from −2s to 6s around the lever press on trial i and ‘ is the error term.

Given that the mice’s choices were correlated across neighboring trials, we weighted the logistic regression based on the frequency of each trial type combination. This was to ensure that choice decoding of a given trial was a reflection of the identity of the lever press on the current trial as opposed to that of the previous or future trial. Thus, we classified each trial as one of eight ‘press sequence types’ based on the following ‘previous-current-future’ press sequences: ipsi-ipsi-ipsi, ipsi-ipsi-contra, ipsi-contra-contra, ipsi-contra-ipsi, contra-contra-contra, contra-contra-ipsi, contra-ipsi-ipsi, contra-ipsi-contra. We then used this classification to equalize the effects of press-sequence type on our decoder by generating weights corresponding to the inverse of the frequency of the press sequence type of that trial. These weights were then used as an input to the *fitglm* function in MATLAB, which was used to fit a weighted version of the logistic regression model above (**Equation 8**).

Decoder performance was evaluated with 5-fold cross-validation by calculating the proportion of correctly classified held-out trials. Predicted ipsilateral press probabilities greater than or equal to 0.5 were decoded as an ipsilateral choice and values less than 0.5 were decoded as a contralateral choice. This was repeated with 100 combinations of randomly selected, simultaneously imaged neurons from each mouse. Reported decoding accuracy is the average accuracy across the 100 runs and 5 combinations of train-test data for each mouse. Note that only 6/7 mice in the PL-NAc cohort were used in the decoder analyses as one mouse had fewer than 10 simultaneously imaged neurons.

#### Outcome decoder

For the outcome decoder in **Figure 3d**, we used the same weighted logistic regression used for choice decoding, except the dependent variable was the outcome of the trial (+1 for a reward, 0 for no reward) and the predictors were the average GCaMP6f fluorescence during the intertrial interval (ITI) of each trial. The ITI was defined as the time between CS presentation and either 1s before the next trial’s nose poke or 8s after the CS, whichever occurred first. This was used in order to avoid including any neural activity attributable to the next trial’s nose poke in our analysis.

To correct for outcome correlations between neighboring trials, we performed a similar weighting of predictors as performed in the choice decoder above using the following eight outcome sequence types: ‘reward-reward-reward’, ‘reward-reward-unreward’, ‘reward-unreward-unreward’, ‘reward-unreward-reward’, ‘unreward-unreward-unreward’, ‘unreward-unreward-reward’, ‘unreward-reward-reward’, ‘unreward-reward-unreward.’

#### Time-course choice decoder

To determine how well activity from PL-NAc and mTH-NAc neurons was able to predict the mouse’s choice as a function of time throughout the trial (**Figure 4e** & **Supplementary Figure 7e**), we trained separate logistic regressions on 500ms bins throughout the trial, using the GCaMP6f fluorescence of 10 simultaneously imaged neurons.

Because of the variability in task timing imposed by the jitter and variability of the mice’s actions, we linearly interpolated the GCaMP6f fluorescence trace of each trial to a uniform length, *t_adjusted_*, relative to behavioral events in our task. Specifically, for each trial, *T*, we divided time into the following four epochs: (i) 2s before nose poke, (ii) time from the nose poke to the lever press, (iii) time from the lever press to the nose poke of the subsequent trial, *T* + 1 and (iv) the 3s following the next trial nosepoke. For epochs ii and iii, *t_adjusted_* was determined by interpolating the GCaMP6f fluorescence trace from each trial to a uniform length defined as the median time between the flanking events across all trials. Thus, *t_adjusted_* within each epoch for each trial, *T*, was defined as:

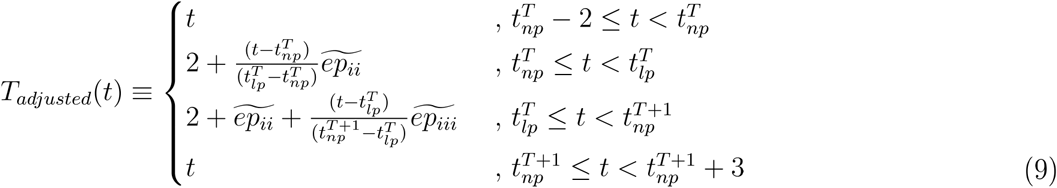

where 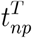, and 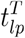 are the times of the nose poke and lever press on the current trial, 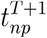 is the time of the nose poke of the subsequent trial and 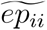 and 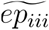 are the median times across trials of epoch ii and iii.

The resulting time-adjusted GCaMP6f traces were divided into 500ms bins. For each bin, we fit the weighted logistic regression described above to predict choice on the current, previous or future trial from the activity of 10 simultaneously imaged neurons. Predictors were weighted based on press sequence type as described above. Decoding accuracy was assessed as described above using 100 combinations of 10 randomly selected neurons and 5-fold cross-validation. To determine if decoding was significantly above chance, which is 0.5, for each timepoint we performed a two-tailed, one-sample t-test.

### Statistics

All t-tests reported in the results were performed using either the *ttest* or *ttest2* function in MATLAB. In all cases, t-tests were two-tailed. In cases where multiple comparisons were performed, we applied a Bonferroni correction to determine the significance threshold. Two-proportion Z-tests (used to compare fractions of significantly modulated/selective neurons, **Figures 2g,h & 3a,c**) and Fisher’s Z (used to compare correlation coefficients, **Figure 4d & Supplementary Figure 7d**) were performed using Vassarstats.net.

For all t-tests in this paper, data distributions were assumed to be normal, but this was not formally tested. No statistical methods were used to predetermine sample sizes, but our sample sizes were similar to those generally employed in the field.

### Synaptic plasticity model

To computationally model how the brain could solve the reversal learning task *using fast dopamine-mediated synaptic plasticity*, we generated a biological instantiation of the TD algorithm for reinforcement learning (Sutton and Barto, 1998) by combining the recorded PL-NAc activity with known circuit connectivity in the NAc and associated regions (Hunnicutt et al., 2016; Kalivas et al., 1993; Otis et al., 2017; Watabe-Uchida et al., 2012). The goal of the model is to solve the “temporal credit assignment problem” by learning the value of each choice at the onset of the choice-selective PL-NAc sequence, when we assume the mouse makes its decision and which is well before the time of reward.

### Synaptic plasticity model description

#### The value function

Our implementation of the TD algorithm seeks to learn an estimate, at any given time, of the total discounted sum of expected future rewards, known as the value function W). To do this, we assume that the value function over time is decomposed into a weighted sum of temporal basis functions (Sutton and Barto, 1998) corresponding to the right-choice and left-choice preferring neurons:

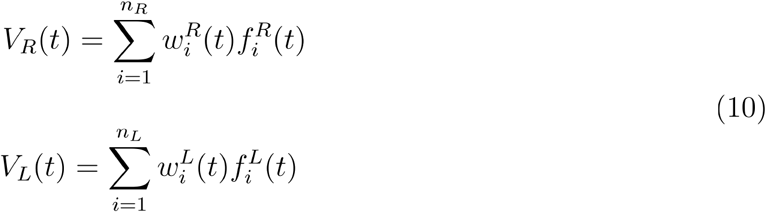

with the total value being given by the sum over both the left and right neurons as

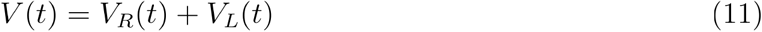

Here, *V_R_*(*t*) and *V_L_*(*t*) are the components of the value functions encoded by the right- and left-preferring neurons respectively, *n_R_* and *n_L_* are the number of right- and left-preferring choice-selective neurons respectively, and 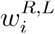 are the weights between the *i^th^* PL neuron and the NAc, which multiply the corresponding basis functions. Thus, each term in *V_R_*(*t*) or *V_L_*(*t*) above corresponds to the activity of one of the striatal neurons in the model (**Figure 5a**). Note that, in our model, the total value *V*(*t*) sums the values associated with the left and right actions and is thus not associated with a particular action. At any given time on a given trial, the choice-selective activity inherent to the recorded PL-NAc neurons results predominantly in activation of the sequence corresponding to the chosen lever compared to the unchosen lever (see **Figure 5c**), so that a single sequence, corresponding to the chosen action, gets reinforced.

#### The reward prediction error (RPE)

TD learning updates the value function iteratively by computing errors in the predicted value function and using these to update the weights *w_i_*. The RPE at each moment of time is calculated from the change in the estimated value function over a time step of size *dt* as follows

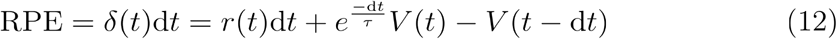

where *δ*(*t*) is the reward prediction error per unit time. Here, the first two terms represent the estimated value at time *t*, which equals the sum of the total reward received at time *t* and the (discounted) expectation of rewards, i.e. value, at all times into the future. This is compared to the previous time step’s estimated value *V*(*t* – *dt*). The coefficient 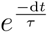 represents the temporal discounting of rewards incurred over the time step d*t*. Here *τ* denotes the timescale of temporal discounting and was chosen to be 0.8*s*.

To translate this continuous time representation of RPE signals to our biological circuit model, we assume that the RPE *δ*(*t*), is carried by dopamine neurons (Montague et al., 1996; Schultz et al., 1997). These dopamine neurons receive three inputs corresponding to the three terms on the right side of the above equation: a reward signal originating from outside the VTA, a discounted estimate of the value function *V*(*t*) that, in **Figure 5a**, represents input from the striatum via the ventral pallidum (Chen et al., 2019; Tian et al., 2016) and an oppositely signed, delayed copy of the value function *V*(*t* – Δ) that converges upon the VTA interneurons (Cohen et al., 2012).

Because the analytical formulation of TD learning in continuous time is defined in terms of the infinitesimal time step *dt*, but a realistic circuit implementation needs to be characterized by a finite delay time for the disynaptic pathway through the VTA interneurons, we rewrite the above equation approximately for small, but finite delay Δ as:

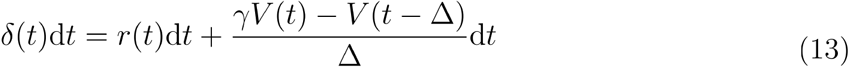

where we have defined 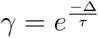 as the discount factor corresponding to one interneuron time delay and, in all simulations, we chose a delay time Δ = 0.01*s*. Note that the discount factor is biologically implemented in different strengths of the weights of the VP inputs to the GABA interneuron and dopaminergic neuron in the VTA.

The proposed circuit architecture of **Figure 5a** can be rearranged into several other, mathematically equivalent architectures (**Supplementary Figure 9**). These architectures are not mutually exclusive, so other more complicated architectures could be generated by superpositions of these architectures.

#### The eligibility trace

The RPE at each time step *δ*(*t*)d*t* was used to update the weights of the recently activated synapses, where the “eligibility” *E_i_*(*t*) of a synapse for updating depends upon an exponentially weighted average of its recent past activity (Gerstner et al., 2018; Sutton and Barto, 1998):

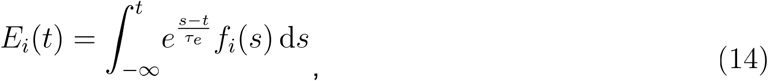

which can be rewritten as

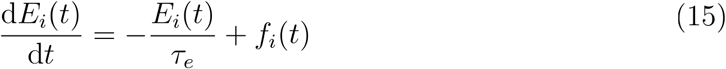

or, in the limit *dt* ≫ 1,

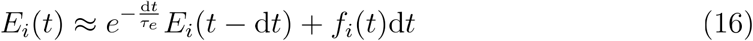

where *τ_e_* defines the time constant of the decay of the eligibility trace, which was chosen to be 0.6s consistent with (Gerstner et al., 2018; Yagishita et al., 2014).

#### Weight Updates

The weight of each PL-NAc synapse, *w_i_*, is updated according to the product of its eligibility *E_i_*(*t*) and the RPE rate *δ*(*t*) at that time using the following update rule (Gerstner et al., 2018; Sutton and Barto, 1998):

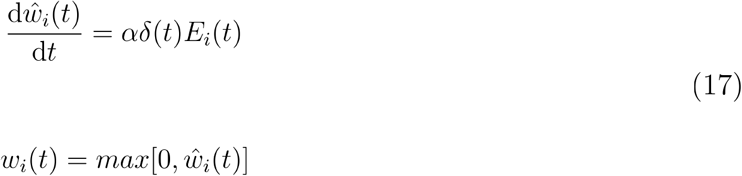

where *α* = 0.009(*spikes/s*)^−1^ was the learning rate. Note that the units of *α* derive from the units of weight being value · (spikes/s)^−1^ The PL-NAc weights used in the model are thresholded to be non-negative so that the weights obey Dale’s principle.

#### Action Selection

In the model, the decision to go left or right is determined by “probing” the relative values of the left versus right actions just prior to the start of the choice-selective sequence. To implement this, we assumed that the choice was read out in a noisy, probabilistic manner from the activity of the first 60 neurons in each (left or right) PL population prior to the start of the sequential activity. This was accomplished by providing a 50 ms long, noisy probe input to each of these PL neurons and reading out the summed activity of the left and the summed activity of the right striatal populations. The difference between these summed activities was then put through a softmax function (given below) to produce the probabilistic decision.

To describe this decision process quantitatively, we define the probability of making a leftward or rightward choice in terms of underlying decision variables *d_left_* and *d_right_* corresponding to the summed activity of the first 60 striatal neurons in each population:

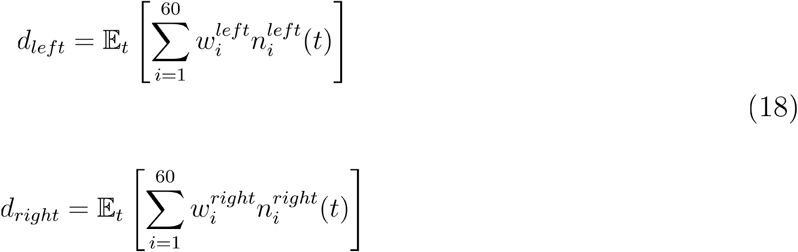

where 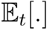 denotes time-averaging over the 50 ms probe period and 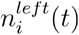 and 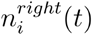 denote the non-negative stochastic probe input, which was chosen independently for each neuron and each time step from a normal distribution (truncated at zero to enforce non-negativity) with mean prior to truncation equal to 0.05 s^−1^ (5% of peak activity) and a standard deviation of 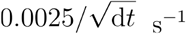. Note that the weights 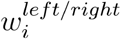 used here correspond to the weights from the end of the previous trial, which we assume are the same as the weights at the beginning of the next trial. The probability of choosing the left or the right lever for a given trial *n* is modeled as a softmax function of these decision variables plus a “stay with the previous choice” term that models the tendency of mice in our study to return to the previously chosen lever irrespective of reward (**Figure 1d**), given by the softmax distribution

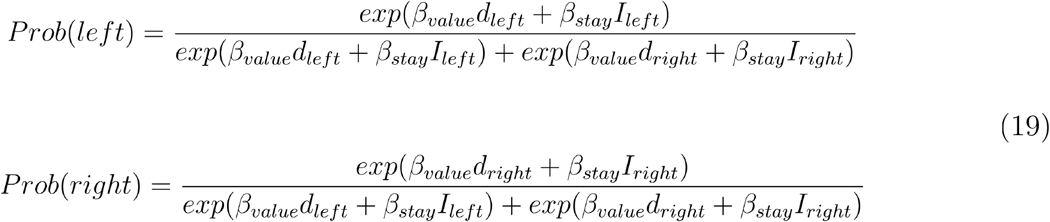

where *I_left/right_* is 1 if that action (i.e. left or right) was chosen on the previous trial and 0 otherwise, and *β_value_* ν 2500 and *β_stay_* = 0.2 are free parameters that define the width of the softmax distribution and the relative weighting of the value-driven versus stay contributions to the choice.

### Synaptic plasticity model implementation

#### Block structure for the model

Block reversals were determined using the same criteria as in the probabilistic reversal learning task performed by the mice – the identity of the rewarded lever reversed after 10 rewarded trials plus a random number of trials drawn from the geometric distribution given by **Equation 1**. The model used p = 0.4 as in the reversal learning experiments. Given the variation in performance across the models that use PL-NAc, mTH-NAc or PL-NAc synchronous activity as input (see **Figure 5**), the average block length for each model varied as well (because block reversals depended upon the number of rewarded trials). The average block length for the single-trial PL-NAc model, single-trial mTH-NAc model and PL-NAc synchronous control were 22.6+/-6.3, 25.7+/-8.5 and 26.5+/-6.6 trials (mean +/- std. dev.), respectively. The PL-NAc model produced a similar block length as that of behaving mice (23.23 +/- 7.93 trials, mean +/- std. dev.). Because a block reversal in our task is dependent on the mice receiving a set number of rewards, the choices just prior to a block reversal are more likely to align with the identity of the block and result in reward (see **Figure 5e,j,o**). Thus, the increase in choice probability observed on trials close to the block reversal (most notably in the synchronous case in **Figure 5o**, but also notable in the mTH-NAc in **Figure 5j** and the PL-NAc in **Figure 5e**) is an artifact of this reversal rule and not reflective of the model learning choice values.

#### PL-NAc inputs to the neural circuit model

To generate the temporal basis functions, *f_i_*(*t*) (example activity shown in **Figure 5c**), we used the choice-selective sequential activity recorded from the PL-NAc neurons shown in **Figure 4b-c**. Spiking activity was inferred from calcium fluorescence using the CNMFe algorithm (Zhou et al., 2018) and choice-selectivity was determined using the nested comparison model used to generate **Figure 3a** (see ***Quantification of neural modulation to behavioral events*** above for details). Model firing rates were generated by Z-scoring the inferred spiking activity of each choice-selective PL-NAc neuron. The resulting model firing rates were interpolated using the *interp* function from Python’s numpy package to match the time step, *dt* = 0.01*s*, and smoothed using a Gaussian kernel with zero mean and a standard deviation of 0.2s using the *gaussian_filter1d* function from the ndimage module in Python’s scikit-learn package.

To generate a large population of model input neurons on each trial, we created a population of 368 choice-selective “pseudoneurons” on each trial. This was done as follows: for each simulated trial, we created 4 copies (pseudoneurons) of each of the 92 recorded choice-selective PL-NAc neurons using that neuron’s inferred spiking activity from 4 different randomly selected trials. The pool of experimentally recorded trials from which pseudoneuron activities were chosen was balanced to have equal numbers of stay and switch trials. This was done because the choices of the mice were strongly positively correlated from trial to trial (i.e., had more stay than switch trials), which (if left uncorrected) potentially could lead to biases in model performance if activity late in a trial was reflective of choice on the next, rather than the present trial. To avoid choice bias in the model, we combined the activity of left- and right-choice-preferring recorded neurons when creating the pool of pseudoneurons. We then randomly selected 184 left-choice-preferring and 184 right-choice-preferring model neurons from this pool of pseudoneurons. An identical procedure, using the 92 most choice-selective mTH-NAc neurons, was followed to create the model mTH-NAc neurons. The identity of these 92 neurons was determined by ranking each neuron’s choice-selectivity using the p-value calculated to determine choice-selectivity (see ***Quantification of neural modulation to behavioral events*** above for details).

To generate the synchronous PL-NAc activity used in **Figure 5m-q**, the temporal basis function of each PL neuron was time-shifted so the peak probability of firing was 2s before the time of the lever press.

To mimic the PL-NAc activity during the optogenetic stimulation of PL-NAc neurons (**Figure 6b**), we set 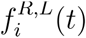 equal to 0.3 for a randomly selected 65% of PL neurons, at all times *t*, from the time of the simulated nosepoke to 2s after the reward presentation. These ‘stimulation trials’ occurred on a random 10% of trials. 65% of PL neurons were activated to mimic the incomplete penetrance of ChR2 viral expression.

#### Reward input to the neural circuit model

The reward input *r*(*t*) to the dopamine neurons was modeled by a truncated Gaussian temporal profile centered at the time of the peak reward:

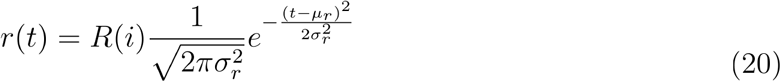

where *R*(*i*) is 1 if trial *i* was rewarded and 0 otherwise, *μ_r_* is the time of peak reward and *σ_r_* defines the width of the Gaussian (0.2s in all cases, width chosen to approximate distribution of dopamine activity in response to reward stimuli observed in previous studies (Matsumoto and Hikosaka, 2009; Schultz et al., 1997). For each trial, a value of *μ_r_* was randomly drawn from a uniform distribution spanning 0.2-1.2s from the time of the lever press. This distribution was chosen to reflect the 1s jitter between lever press and reward used in our behavioral task (see Methods above) as well as the observed delay between reward presentation and peak dopamine release in a variety of studies (Cohen et al., 2012; Matsumoto and Hikosaka, 2009; Parker et al., 2016; Saunders et al., 2018). To ensure that no residual reward response occurred before the time of the lever press, *r_a_*(*t*) was set to 0 for any time *t* that was 0.2s before the time of the peak reward, *μ_r_*.

#### Initial weights

The performance of the model does not depend on the choice of the initial weights as the model learns the correct weights by the end of the first block irrespective of the chosen initial weights. We chose the initial weights to be zero.

#### Weight and eligibility update implementation

We assumed that the weight and eligibility trace updates start at the time of the simulated nose poke. The nose poke time, relative to the time of the lever press, varies due to a variable delay between the nose poke and the lever presentation as well as variation in time between lever presentation and lever press. To account for this, the weight and eligibility trace updates are initiated at time *t* = *t_start_*, where *t_start_* was drawn from a Gaussian distribution with a mean at −2.5s, and a variance of 0.2s, which was approximately both the time of the nose poke and the time at which choice-selective sequences initiated in the experimental recordings. The eligibility trace is reset to zero at the beginning of each trial. We stopped updating the weights at the end of the trial, defined as 3s after the time of lever press. The eligibility traces were updated according to **Equation 16**. The weights were updated by integrating **Equation 17** with a first-order forward Euler routine. In all simulations, we used a simulation time step d*t* = 0.01*s*.

### Neural dynamics model

To computationally model how the brain could solve the reversal learning task *without fast dopamine-mediated synaptic plasticity*, we used an actor-critic network based on the meta-RL framework introduced by Wang et al. (2018). The model actor and critic networks are recurrent neural networks of Long Short Term Memory (LSTM) units whose weights are learned slowly during the training phase of the task. The weights are then frozen during the testing phase so that fast reversal learning occurs only through the activation dynamics of the recurrent actor-critic network. Like the synaptic plasticity model, we input recorded PL-NAc activity to a value-generating “critic” network (conceived of as NAc, VP, and associated cortical regions) to generate appropriate reward prediction error signals in dopamine neurons. Unlike the synaptic plasticity model, the reward prediction error signals in this model are sent to an explicit actor network (conceived of as DMS and associated cortical regions), where they act as an input to help generate appropriate action signals based on reward history.

### Neural dynamics model description

#### LSTM

The model comprises two separate fully connected, gated recurrent neural networks of LSTM units, one each for the actor and critic network. An LSTM unit works by keeping track of a ‘long-term memory’ state (‘memory state’ ***c***(*t*), also known as cell state) and a ‘short-term memory’ state (‘output state’ ***h***(*t*), also known as hidden state) at all times. To regulate the information to be kept or discarded in the memory and output states, LSTMs use three types of gates: the input gate ***i***(*t*) regulates what information that is input to the network, the forget gate ***ϕ***(*t*) regulates what information to forget from the previous memory state, and the output gate ***o***(*t*) (not to be confused with the output state ***h***(*t*)) regulates the output of the network. More precisely, the dynamics of an LSTM is defined by the following equations:

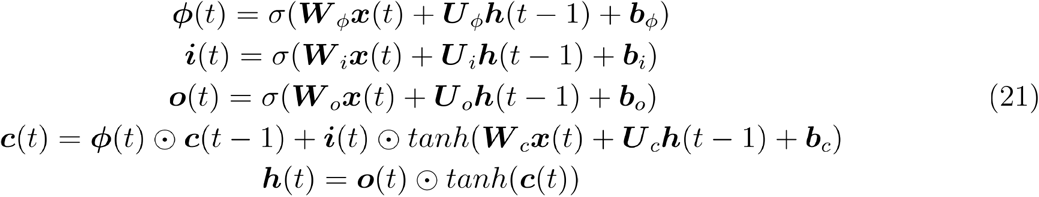

where ***x***(*t*) is the vector of external inputs to the LSTM network at time step *t*, ***W***_*q*_ and ***U***_*q*_ are the weight matrices of the input and recurrent connections, respectively, where the subscript *q* denotes the state or gate being updated, ***b***_*q*_ are the bias vectors, ⊙ denotes element-wise multiplication and *σ* denotes the sigmoid function defined as follows:

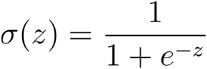

#### Critic network

As in the synaptic plasticity model, the goal of the critic is to learn the value (discounted sum of future rewards) of a given choice at any time in a trial. The learned value signal can then be used to generate the RPE signals that are sent to the actor. The critic is modelled as a network of LSTM units that linearly project through trainable weights to a value readout neuron that represents the estimated value *V*(*t*) at time step *t*. The critic takes as input the reward received at the previous time step *r*(*t* − 1) and the experimentally recorded PL-NAc choice-selective sequential input *C*(*t*). The PL-NAc input provides the critic with a representation of the chosen side on the current trial as well as the time during the trial. This allows the critic to output an appropriately timed value signal (and consequently an appropriately timed RPE signal) corresponding to the chosen side. The reward input acts as a feedback signal to the critic that evaluates the correctness of the generated value signal.

To map the critic to a biological neural circuit, we hypothesize that NAc, together with VP and associated cortical areas, form the critic recurrent neural network (**Figure 7a**; Atallah et al., 2007; Lau and Glimcher, 2008; O’Doherty et al., 2004; Richard et al., 2016; Tsutsui et al., 2016). The choice-selective sequential input *C*(*t*) to the critic is provided by the recorded choice-selective sequential activity in PL-NAc neurons (**Figure 7a**).

#### The reward prediction error (RPE)

As in the synaptic plasticity model (**Figure 5a**), the RPE *δ*(*t*) is computed in the VTA DA neurons based on the value signal from the critic network (**Figure 7a**).

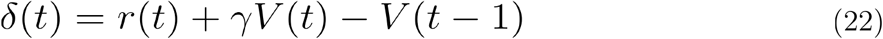

Unlike the synaptic plasticity model, the RPE signal is conveyed by the VTA dopamine neurons to the actor network.

#### Actor network

In contrast to the synaptic plasticity model, in which actions were directly readout from the activity of the value neurons early in the trial, we consider an explicit actor network that generates actions. The actor is modelled as a network of LSTM units that compute the policy, i.e., the probability of choosing an action *a*(*t*) at time step *t* given the current state of the network. The policy is represented by three policy readout neurons, corresponding to choosing left, right or ‘do nothing’, whose activities are given by a (trainable) linear readout of the activities of the actor LSTM units. The actor receives three inputs: (i) an efference copy of the action taken at the previous time step *a*(*t* – 1), (ii) a ‘temporal context’ input *ξ*(*t*), encoded as a vector of all 0s except for a value of 1 in the entry corresponding to the current time point *t*, that provides the actor with a representation of the time within the trial, and (iii) the reward prediction error at the previous time step *δ*(*t* – 1).

To map the actor to a biological neural circuit, we hypothesize that the DMS and associated cortical areas form the actor recurrent neural network (**Figure 7a**; Atallah et al., 2007; O’Doherty et al., 2004; Seo et al., 2012; Tai et al., 2012). The temporal sequence input *ξ*(*t*) to the actor is assumed to originate in the hippocampus or other cortical areas (**Figure 7a**; Hahnloser et al., 2002; Howard and Eichenbaum, 2013; Kozhevnikov and Fee, 2007; Zhou et al., 2020).

#### Training algorithm

To train the recurrent weights of the network, which are then held fixed during task performance, we implement the Advantage Actor-Critic algorithm (Mnih et al., 2016) on a slightly modified version of the reversal learning task (see “Block structure for training” section below). In brief, the weights of the neural network are updated via gradient descent and backpropagation through time. The loss function for the critic network, 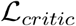, defines the error in the estimated value function V(t). The synaptic weight parameters *θ_υ_* of the critic network are updated through gradient descent on the critic loss function 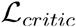:

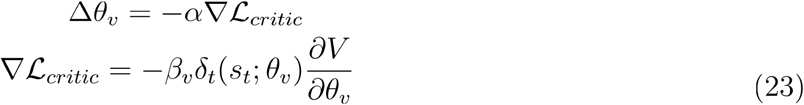

where *α* is the learning rate, *s_t_* is the state at time step *t, V* denotes the value function and *β_υ_* is the scaling factor of the critic loss term. *δ_t_*(*s_t_*; *θ_υ_*) is the k-step return temporal difference error (not to be confused with the RPE input to the actor defined in **Equation 22**) defined as follows:

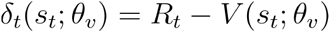

where *R_t_* is the discounted k-step bootstrapped return at time *t*

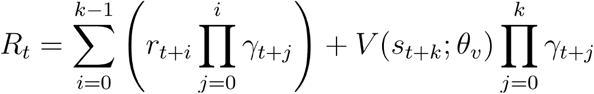

where *r_t_* is the reward received at time step *t, γ_t_* is the discount factor at time step *t* (defined below), and *k* is the number of time steps until the end of an episode.

The loss function for the actor network, 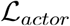, is given by a weighted sum of two terms: a policy gradient loss term, which enables the actor network to learn a policy *π*(*a_t_|s_t_*) that approximately maximizes the estimated sum of future rewards *V*(*s_t_*), and an entropy regularization term that maximize the entropy of the policy *π* to encourage the actor net.work to explore by avoiding premature convergence to suboptimal policies. The gradient of the actor loss function 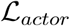 with respect to the synaptic weight parameters of the actor network, *θ*, is given by

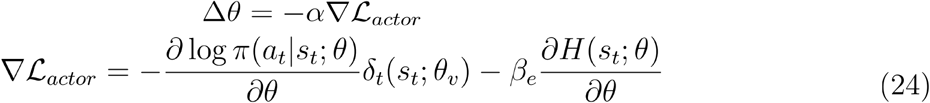

where *a_t_* is the action at time step *t, π* is the policy, *β_e_* is the scaling factor of the entropy regularization term and *H*(*s_t_;θ*) is the entropy of the policy *π*

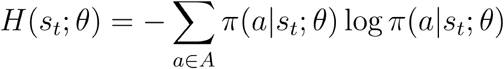

where *A* denotes the space of all possible actions.

### Neural dynamics model implementation

#### LSTM

Both the actor and critic LSTM networks consisted of 128 units each and were implemented using TensorFlow’s Keras API. The weight matrices ***U***_*q*_ were initialized using Keras’s ‘glorot_uniform’ initializer, the weight matrices ***W***_*q*_ were initialized using Keras’s ‘orthogonal’ initializer and the biases ***b*** were initialized to 0. The output and memory states for both LSTM networks were initialized to zero at the beginning of each training or testing episode.

#### PL-NAc inputs to the critic

Input to the critic was identical to the smoothed, single-trial input used for the synaptic plasticity model described above, except i) activity was not interpolated because each time step in this model was equivalent to the sampling rate of the collected data (10 Hz), and ii) we chose to input only the activity from 2s before to 2s after the lever press (as compared to 3s after the lever press for the synaptic plasticity model) in order to reduce the computational complexity of the training process. To reduce episode length, and therefore training time, we also excluded those neurons whose peak activity occurred more than 2s after the lever press, reducing the final number of ‘pseudoneurons’ used as input to 306 (compared with 368 for the synaptic plasticity model).

Optogenetic-like stimulation of the PL-NAc population was performed in a similar manner to the synaptic plasticity model, with activity set to 0.15 for a randomly selected 60% of neurons for the duration of the trial.

#### Trial structure

Each trial was 4s long starting at 2s before lever press and ending at 2s after lever press. At any given time, the model has three different choices - choose left, choose right or do nothing. Similar to the synaptic plasticity model, the model makes its decision to choose left or right at the start of a trial, which then leads to the start of the corresponding choice-selective sequential activity. However, unlike the synaptic plasticity model, the model can also choose ‘do nothing’ at the first time step, in which case an activity pattern of all zeros is input to the critic for the rest of the trial. For all other time steps, the correct response for the model is to ‘do nothing’. Choosing ‘do nothing’ on the first time step or choosing something other than ‘do nothing’ on the subsequent time steps results in a reward *r*(*t*) of −1 at that time. If a left or right choice is made on the first time step, then the current trial is rewarded based on the reward probabilities of the current block (**Figure 1a**) and the reward input *r*(*t*) to the critic is modeled by a truncated Gaussian temporal profile centered at the time of the peak reward (**Equation 20**) with the same parameters as in the synaptic plasticity model.

#### Block structure for training

We used a slightly modified version of the reversal learning task performed by the mice in which the block reversal probabilities were altered in order to make the block reversals unpredictable. This was done to discourage the model from learning the expected times of block reversals and to instead mimic the results of our behavioral regressions (**Figure 1e**) suggesting that the mice use only the previous ~4 trials to make a choice. To make the block reversals unpredictable, the identity of the high-probability lever reversed after a random number of trials drawn from a geometric distribution (**Equation 1**) with *p* = 0.9.

#### Training

Each training episode was chosen to be 15 trials long and the model was trained for 62000 episodes. The values of the training hyperparameters were as follows: the scaling factor of the critic loss term *β_υ_* = 0.05, the scaling factor of the entropy regularization term *β_e_* = 0.05, the learning rate *α* = 0.01*s*^−1^ (*α* = 0.001 per time step), and the timescale of temporal discounting within a trial *τ* = 2.45*s*, leading to a discount factor *γ* = *e^−dt/τ^* = 0.96 for all times except for the last time step of a trial when the discount factor was 0 to denote the end of a trial. The network’s weights and biases were trained using the RMSprop gradient descent optimization algorithm (Hinton et al., 2012) and backpropagation through time, which involved unrolling the LSTM network over an episode (630 time steps).

#### Block structure for testing

Block reversal probabilities for the testing phase were the same as in the probabilistic reversal learning task performed by the mice. The average block length for the PL-NAc neural dynamics model was 19.9+/-5.3 trials (mean+/-std. dev.).

#### Testing

The model’s performance (**Figures 7b-k**) was evaluated in a testing phase during which all network weights were held fixed so that reversal learning was accomplished solely through the neural dynamics of the LSTM networks. The network weights used in the testing phase were the weights learned at the end of the training phase. A testing episode was chosen to be 1500 trials long and the model was run for 120 episodes.

#### Actor network analysis

For **Figures 7f-i**, we tested the model’s performance on a slightly modified version of the reversal learning task in which block lengths were fixed at 30 trials. This facilitated the calculation and interpretation of the block-averaged activity on a given trial of a block. Dimensionality reduction of the actor network activity (**Figure 7g**) was performed using the *PCA* function from the decomposition module in Python’s scikit-learn package.

#### Replacing sequential input to the critic with persistent input

In **Figure 7e**, we analyzed how model performance changed when the temporal structure provided by the choice-selective sequential inputs to the critic were replaced during training by persistent choice-selective input. The persistent choice-selective input was generated by setting the activity of all the left choice-selective neurons to 1 and all the right-choice selective neurons to 0 for all time points on left-choice trials and vice versa on right-choice trials.

### Cross-trial analysis of RPE in dopamine neurons

To generate the regression coefficients in **Figure 5g,l,q, Figure 7d** and **Supplementary Figure 10c,d**, we performed a linear regression analysis adapted from (Bayer and Glimcher, 2005), which uses the mouse’s reward outcome history from the current and previous 5 trials to predict the average dopamine response to reward feedback on a given trial, *i*:

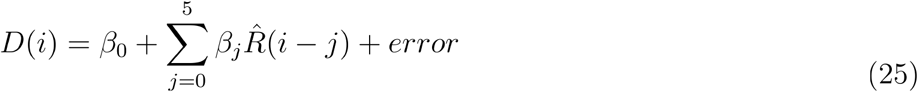

Where *D*(*i*) is the average dopamine activity from 0.2 to 1.2s following reward feedback on trial *i*, 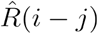 is the reward outcome *j* trials back from trial *i* (1 if *j* trials back is rewarded and 0 if unrewarded) and *β_j_* are the calculated regression coefficients that represent the effect of reward outcome *j* trials back on the strength of the average dopamine activity, *D*(*i*). For the regression coefficients generated from recorded dopamine activity (**Supplementary Figure 10c,d**) we used the Z-scored GCaMP6f fluorescence from VTA-NAc terminal recordings of 11 mice performing the same probabilistic reversal learning task described in this paper (for details see Parker et al., 2016). The regression coefficients for the experimental data as well as the synaptic plasticity and neural dynamics model simulations were fit using the *LinearRegression* function from the linear_model module in Python’s scikit-learn package.

### Optogenetic stimulation of PL-NAc neurons

22 male C57BL/6J mice were bilaterally injected in either the PL (n=14 mice, M–L ± 0.4, A–P 2.0 and D–V −2.5 mm) or mTH (n=8 mice, M–L ± 0.3, A–P −0.7 and D–V −3.6 mm) with 600nl AAV2/5-CaMKIIa-hChR2-EYFP (UPenn vector core, injected 0.6 μl per hemisphere of titer of 9.6 × 10^13^ pp per ml). Optical fibers (300 μm core diameter, 0.37 NA) delivering 1–2 mW of 447 nm laser light (measured at the fiber tip) were implanted bilaterally above the NAc Core at a 10 degree angle (M–L ±1.1, A–P 1.4 and D–V −4.2 mm). An additional cohort of control mice (n=8) were implanted with optical fibers in the NAc without injection of ChR2 and underwent the same stimulation protocol outlined below (**Supplementary Figure 12e-h**). Mice were anesthetized for implant surgeries with isoflurane (3–4% induction and 1–2% maintenance). Mice were given 5 days of recovery after the surgical procedure before behavioral testing.

During behavioral sessions, 5 ms pulses of 1-3 mW, 447 nm blue light was delivered at 20 Hz on a randomly selected 10% of trials beginning when the mouse entered the central nose poke. Light stimulation on unrewarded trials ended 1s after the end of the CS− presentation. On rewarded trials, light administration ended either 1s after CS+ presentation (‘cohort 1’) or at the end of reward consumption, as measured by the mouse not engaging the reward port for 100ms (‘cohort 2’). See **Supplementary Figure 13** for a schematic of stimulation times as well as the behavior of the two cohorts. Mice alternated between sessions with and without stimulation – sessions without stimulation were excluded from analysis. Anatomical targeting was confirmed as successful in all mice through histology after the experiment, and therefore no mice were excluded from this data set.

To quantify the effect of laser stimulation on latency times shown in **Supplementary Figure 12a-d**, we ran a mixed effects linear model using the *fitglme* package in MATLAB. In this model, the median latency to initiate a trial of a mouse, defined as the time between illumination of the central nose poke (i.e., trial start) and the mouse initiating a trial via nose poke, was predicted using i) opsin identity (PL-NAc CaMKII-ChR2, mTH-NAc CaMKII-ChR2 or no-opsin controls), ii) laser stimulation on the current trial, iii) laser stimulation on the previous trial, iv) the interaction between opsin identity and laser stimulation on the current trial and v) the interaction between opsin and laser stimulation on the previous trial. To account for individual variation between mice, a random effect of mouse ID was included.

### Simulation of model-free versus model-based task performance

#### Overview

In order to identify possible RPE signatures that distinguish ideal observer (“model-based”) versus Q-learning (“model-free”) behavior in this task (**Supplementary Figure 14**), we simulated choices using the two models. Based on the dopaminergic signature of block reversal inference reported in Bromberg-Martin et al. (2010), we first confirmed that our ideal observer and Q-learning models gave rise to distinct dopamine signatures when performing the task used in Bromberg-Martin et al. (2010). In this task, reward probabilities were 100% and 0% for the “high probability” and “low probability” choices, respectively, and the reward probabilities reversed with a 5% probability on each trial. Next, we applied the same framework to our task, to determine if we could observe similar distinctions between the models. In this case, the reward probabilities were 70% and 10%, as in the task studied in this paper, and blocks reversed with a 5% probability on each trial, which resulted in block lengths comparable to those observed in our experiments.

#### Ideal observer model

The ideal observer model was provided with knowledge of the reward probabilities associated with each block and the probability of block reversal on each trial. The 5% block reversal probability on each trial can be written in terms of the block state transition probabilities as

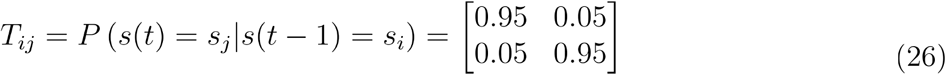

where *T_ij_* is defined as the transition probability between block state *s_i_* on trial *t* and block state *s_j_* on trial *t* + 1. Here, ‘block state’ refers to whether the current block has a higher probability of left or right choices being rewarded. The reward probabilities for each block were as follows

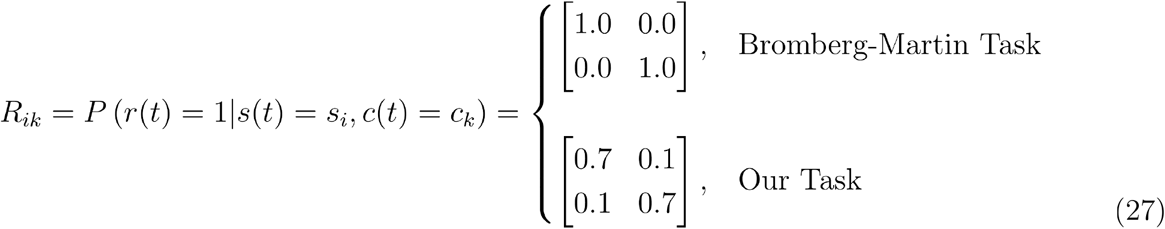

where *R_ik_* is defined as the probability of reward for block state *s_i_* and choice *c_k_*.

On each trial, the ideal observer model selects the choice with the highest expectation of reward based on its belief about the current block state given the choice and reward history. The expectation of reward *ρ_l_*(*t* + 1) for choice *l* on trial *t* + 1, given the entire reward history *r*(1: *t*) and choice history *c*(1: *t*) up until trial *t* is given by

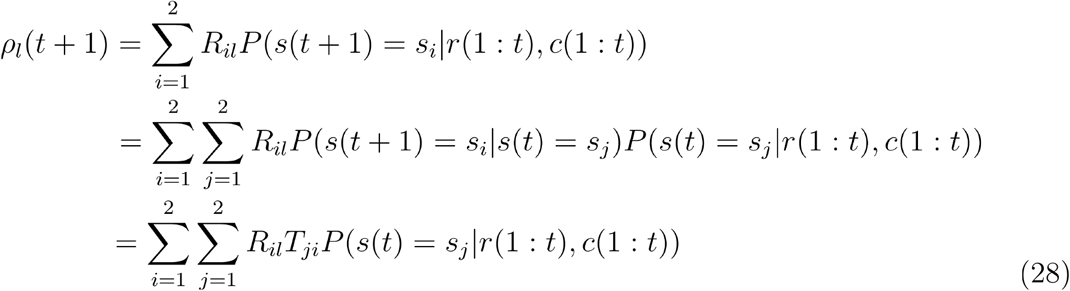

where *l* can be either 1 (left choice) or 2 (right choice) and *P*(*s*(*t*) = *s_j_|r*(1: *t*), *c*(1: *t*)) is the probability of being in block state *s_j_* on trial *t* given the entire reward and choice history up to and including trial *t*. **Equation 28** tells us that estimating the block state probability *P*(*s*(*t*) = *s_j_|r*(1: *t*), *c*(1: *t*) will provide us with an estimate of the expected reward for a given choice on trial *t* + 1 as *R_il_* and *T_ji_* are already known. Using Bayes’ theorem, we can estimate the block state probability as

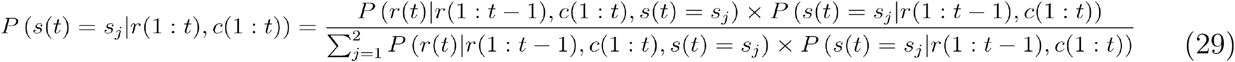

The first term in the numerator of the right hand side of **Equation 29**, *P*(*r*(*t*)|*r*(1: *t* – 1), *c*(1: *t*), *s*(*t*) = *s_j_*), is the probability of receiving reward *r*(*t*) (1 if rewarded and 0 if unrewarded) on trial *t* given the current choice *c*(*t*) = *c_k_*, the block state *s_j_*, and the reward history *r*(1: *t* – 1) and the choice history *c*(1: *t* – 1) up to trial *t* – 1. Because the past history of rewards and choices does not affect the reward probability once the block state is known, this can be rewritten as

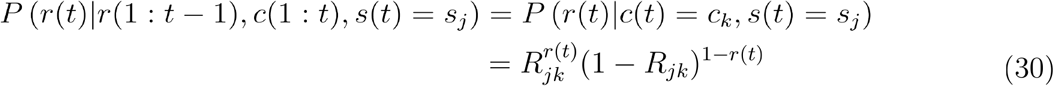

The second term in the numerator of the right hand side of **Equation 29**, *P*(*s*(*t*) = *s_j_*|*r*(1: *t* – 1), *c*(1: *t*)), is the probability that the current block state is *s_j_* given the reward and choice history. This can be rewritten as

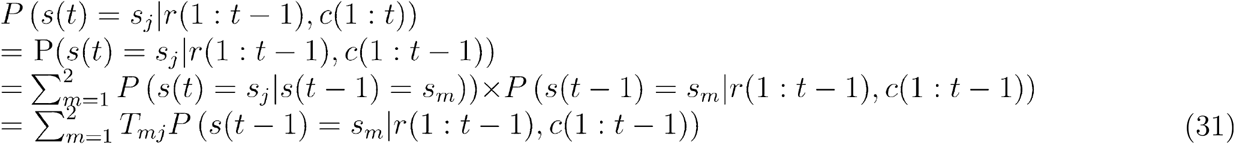

In the second line above, the dependence on c(t) has been removed because the choice on the current trial, in the absence of reward information on the current trial, does not provide any additional information about the current state beyond that provided by the past reward and choice history. Combining **Equations 29–31**, the block state probability on the current trial *t* can be written in terms of the known reward probabilities, known state transition probabilities and the previous block state probability as

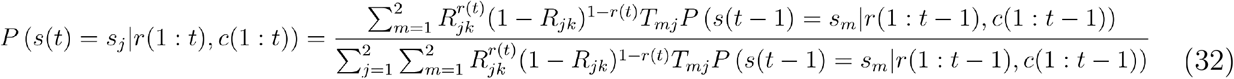

The above equation allows us to estimate the current trial block state probability *P*(*s*(*t*) = *s_j_|r*(1: *t*), *c*(1: *t*)) recursively, since it can be expressed in terms of the previous trial block state probability *P*(*s*(*t* – 1) = *s_m_|r*(1: *t* – 1), *c*(1: *t* – 1)) and other known constant terms. This combined with the known reward and block transition probabilities allows the model to select the optimal choice according to **Equation 28**.

#### Q-Learning model

To simulate trial-by-trial, model-free performance of the tasks, we used a Q-learning model in which the value of the chosen action is updated on each trial as follows:

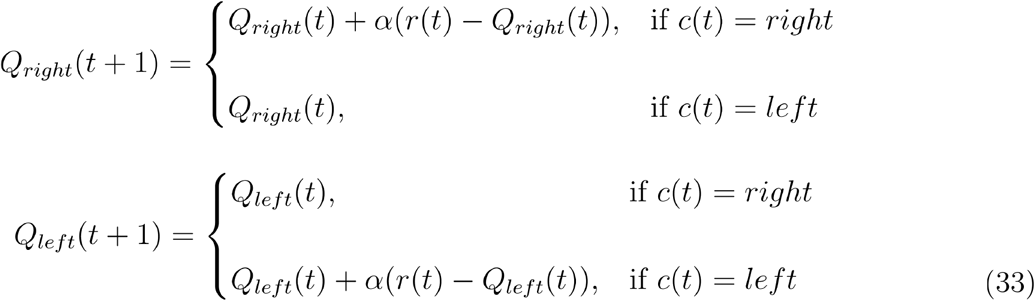

where *Q_right_* is the value for the right choice and *Q_left_* is the value for the left choice *t* is the current trial and *α* is the learning rate, which was set to 0.612 per trial. *r*(*t*) is the outcome of trial *t* (1 for reward, 0 for no reward). Q-values for each choice were initialized to 0. The outcome *r*(*t*) was determined based on the reward probability for choice *c*(*t*) given the block. Choice was simulated using a softmax equation such that the probability of choosing right or left is given by,

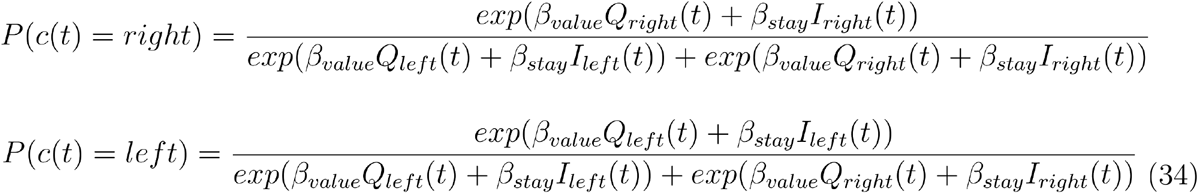

where *β_value_* is the inverse temperature parameter, which was set to 0.99. *β_stay_* is a parameter accounting for how likely mice were to repeat their previous choice, which was set to 0.95. *I_left/right_* is 1 if that action (i.e. left or right) was chosen on the previous trial and 0 otherwise. Parameters for the Q-learning model were fit in Lee et al. (2019) to the behavior of mice in which dopamine neuron activity was recorded in Parker et al. (2016).

#### Comparison of RPE at block reversals

RPE for both the ideal-observer model and the Q-learning model (**Supplementary Figure 14**) was defined as the difference between the experienced reward *r*(*t*) and the expected reward for the chosen action (*ρ_chosen_*(*t*) for the ideal-observer model or *Q_chosen_*(*t*) for the Q-learning model) as follows:

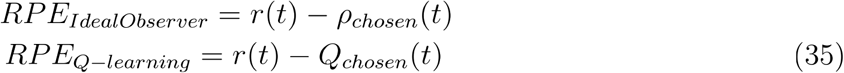

To identify RPE signatures of model free versus model based performance of the two tasks, we compared the RPE from the ideal-observer model and the Q-learning model on trials around block reversals. Specifically, we compared the RPE from the two models on the first trial of a block with the RPE on the second trial of a block when the choice on trial 1 was different from the choice on trial 2. This means that any changes in RPE from trial 1 to trial 2 were inferred because the new action-outcome relationship for the choice made on trial 2 had not been explicitly experienced in the new block.

## Supplemental Figures

**Supplementary Figure 1.**
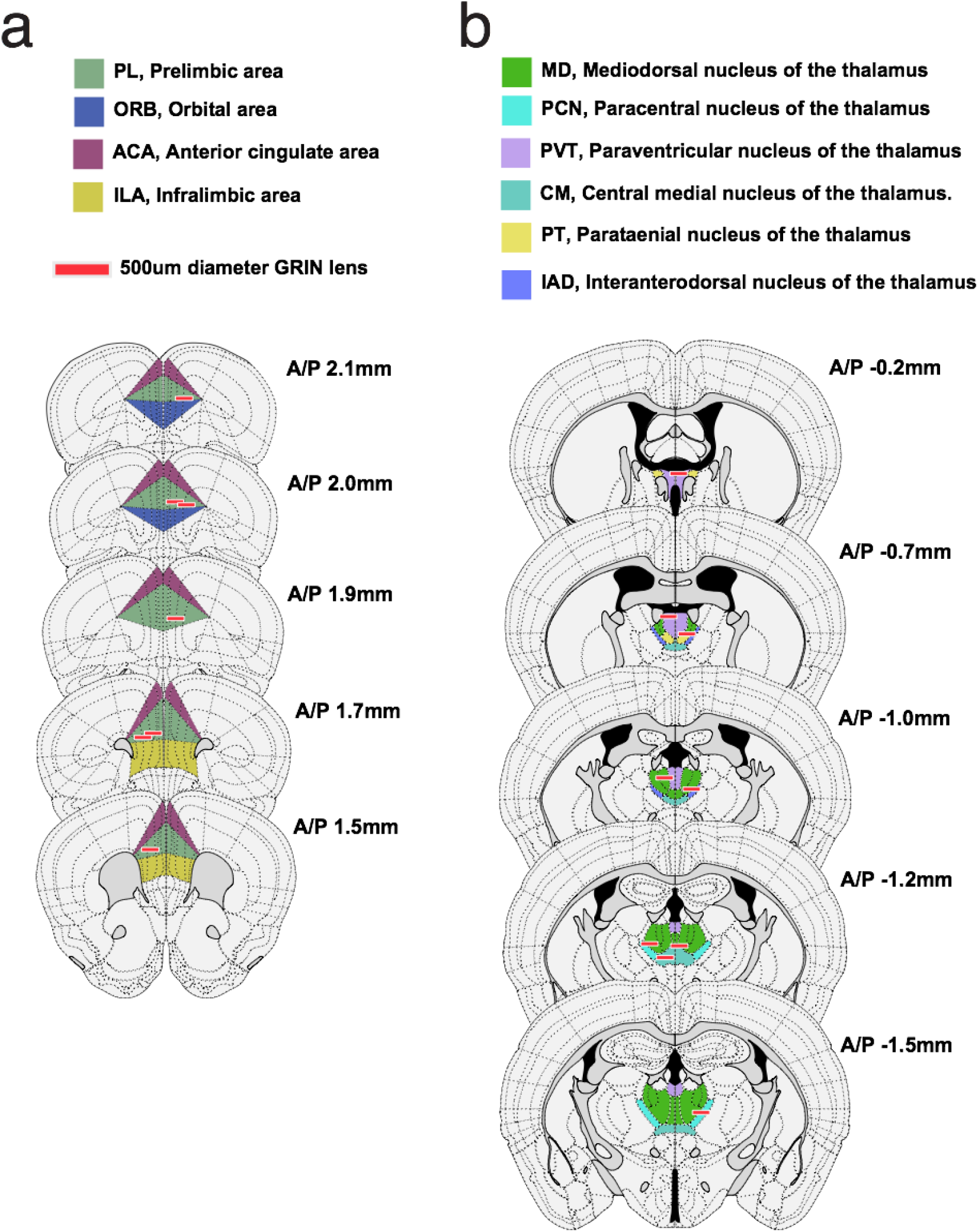
Locations of GRIN lens implants. **(a)** Schematic of coronal sections along the anterior/posterior axis (A/P, numbers relative to bregma) with recording locations of 7 PL-NAc mice. Red lines indicate bottom of lens implant. **(b)** Same as **a** except location of 9 mTH-NAc recordings.

**Supplementary Figure 2.**
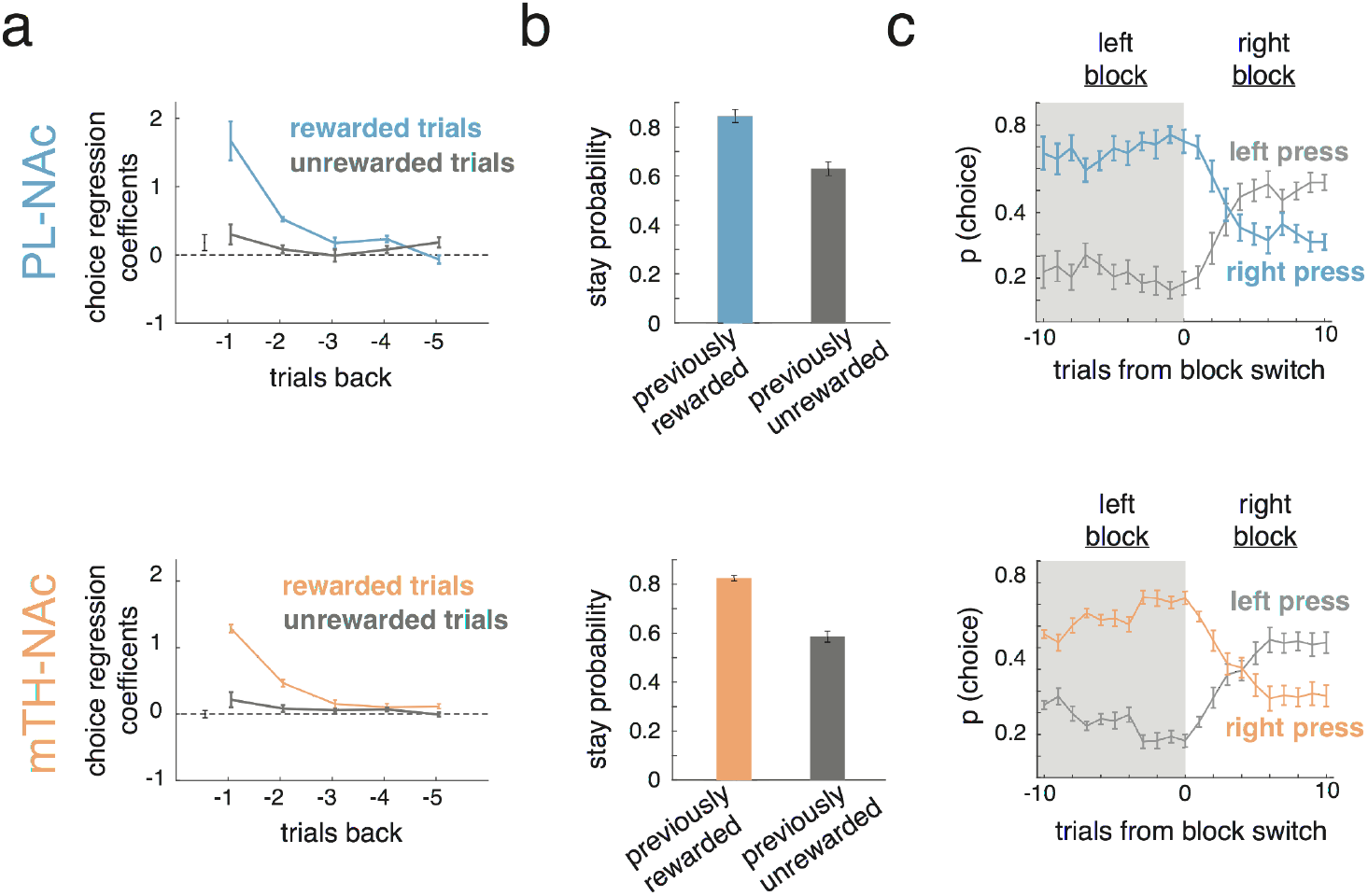
Mice in the PL-NAc and mTH-NAc imaging cohorts have comparable behavior. **(a)** Top, coefficients from logistic regression to predict choice (see **Figure 1**) from PL-NAc cohort (n=7 mice). Bottom, same except coefficients from mTH-NAc cohort (n=9 mice). Both cohorts use choice and outcome information from previous trials to predict the current choice. Regression coefficients between the two cohorts are not significantly different for any trials back for either rewarded or unrewarded trials (P>0.01, unpaired, two-tailed t-test of regression coefficients across mice at each trial back, n=7 and 9 mice for PL-NAc and mTH-NAc, respectively). **(b)** Stay probability following rewarded (blue or orange) and unrewarded (grey) trials for PL-NAc (top) and mTH-NAc (bottom) cohorts. Both cohorts have a significantly higher stay probability following a rewarded trial (PL-NAc: P=0.00008; mTH-NAc: P=0.00003, paired, two-tailed t-test comparing stay probability on rewarded and unrewarded trials across mice, n=7 and 9 mice for PL-NAc and mTH-NAc, respectively). **(c)** Probability of a left or right lever press following a reversal from a left-preferring to right-preferring block of mice from the PL-NAc (top, n=7 mice) and mTH-NAc (bottom, n=9 mice) cohorts. Both cohorts display a qualitatively similar change in choice behavior following a block reversal. In all panels, error bars indicate s.e.m. across mice.

**Supplementary Figure 3.**
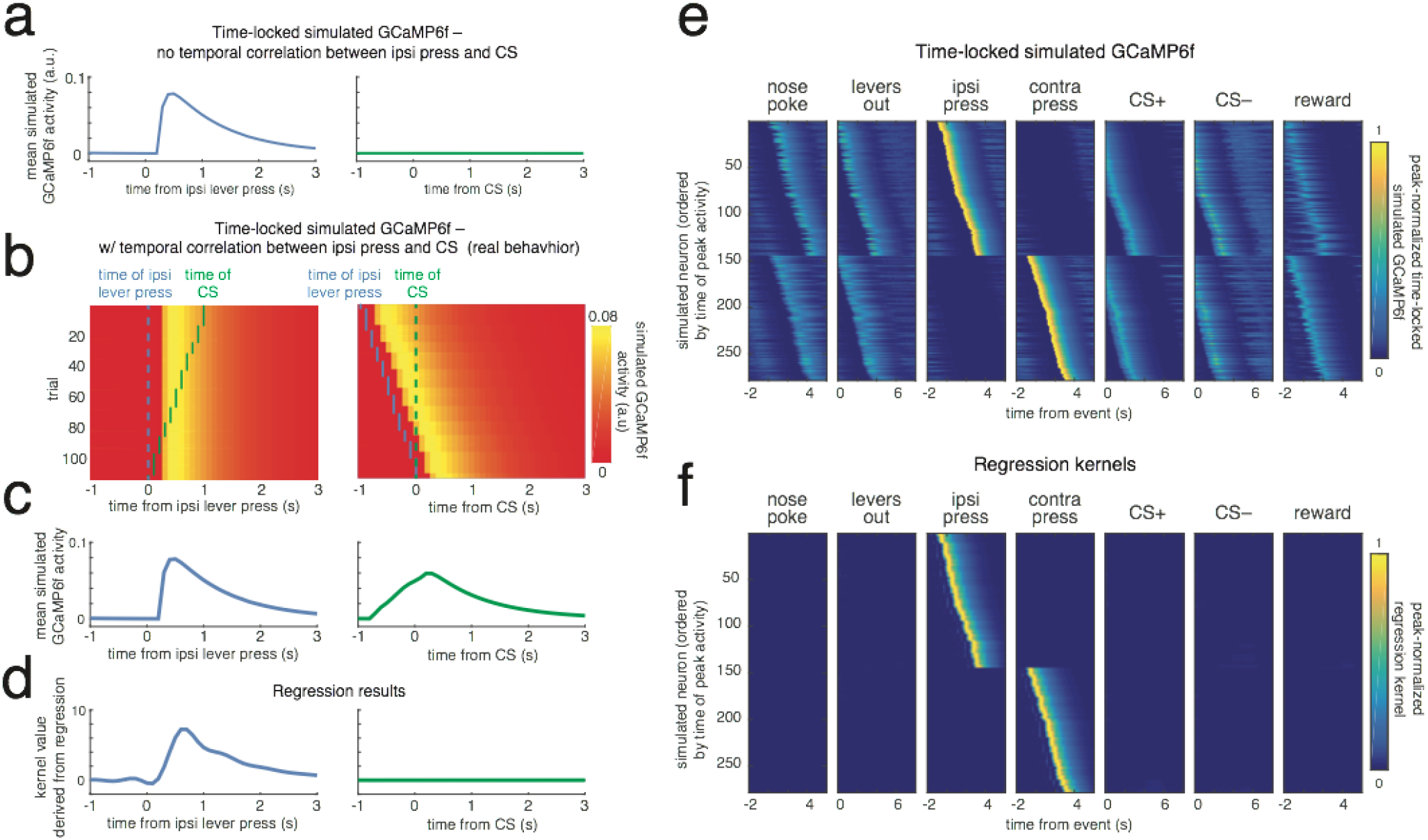
Simulated neural activity to illustrate the ability of the encoding model to successfully relate neural activity to the appropriate behavioral event. **(a)** Simulated neuron that is responsive only to the ipsilateral lever press. **(b)** Trial-by-trial heatmap of a simulated neuron that has increased activity time-locked to the ipsilateral lever press. In the data, there was a correlation between the time of lever press and the time of the CS, which produced a time-locked response to the CS, even though the neuron did not respond to that event. Left, activity heatmap aligned to the time of an ipsilateral lever press (dashed blue line) sorted by the time of the subsequent CS presentation (green dots). Right, activity heatmap is aligned to the time of the CS (dashed green line), ordered by the time of the preceding lever press (blue dots). **(c)** Average activity across trials of the example simulated neuron in **b** aligned to the lever press, left, and CS presentation, right. Unlike the idealized case in **a**, when the timing of task events is maintained from the real behavior, the temporal correlations result in a bump in activity aligned to the CS (right plot). Note that this bump in activity is generated entirely by the correlation in event times, since this simulated neuron only had activity in relation to the lever press (and not the CS presentation). **(d)** Response kernels for lever press, left, and CS, right, derived from the encoding model used to attribute the neural response of individual task events. The model successfully recovers the fact that neural activity in this simulated neuron is related to the lever press and not the CS. **(e)** Heatmap displaying the average activity from a population of 278 simulated neurons that respond to either the ipsilateral or contralateral lever press, but not the other events. Each neuron responds to the lever press, with a randomly assigned response latency from −1 to 3s. While the strongest average time-locked response is to the ipsilateral or contralateral lever presses, there are visible responses to the other task events as a consequence of the correlation between task events resulting from their temporal proximity. **(f)** Same as **e** except heatmap displays the response kernels derived from the encoding model. The model successfully discovers the underlying structure of the data (i.e., that responses are driven by the lever press).

**Supplementary Figure 4.**
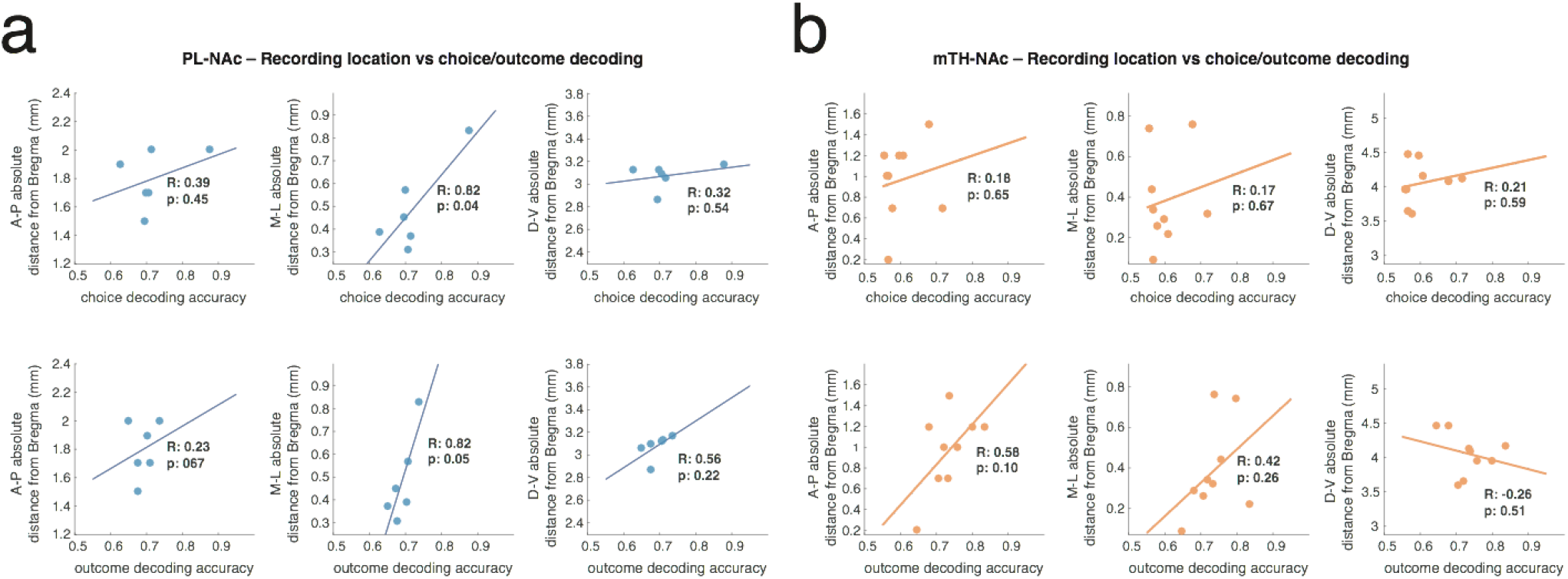
Lack of correlation between recording locations relative to Bregma and choice / outcome decoding. **(a)** Top, correlations between choice decoding accuracy using recorded PL-NAc activity and, in order, the anterior/posterior (A/P), medial/lateral (M/L) and dorsal/ventral (D/V) recording locations relative to Bregma (see **Supplemental Figure 1** for schematic of recording locations; recording locations were aligned to the Allen atlas using the Wholebrain software suite (http://www.wholebrainsoftware.ora/) of Furth et al. (2018); see Methods for details). Bottom, same as top except correlation between recording location and outcome decoding accuracy using PL-NAc activity. **(b)** Same as **a** except decoding accuracy for choice (top) and outcome (bottom) determined using recorded mTH-NAc activity. All p-values are calculated from Pearson’s correlation coefficient; none are significant at the p<.05 level after correction for multiple (6) hypotheses using Bonferroni correction.

**Supplementary Figure 5.**
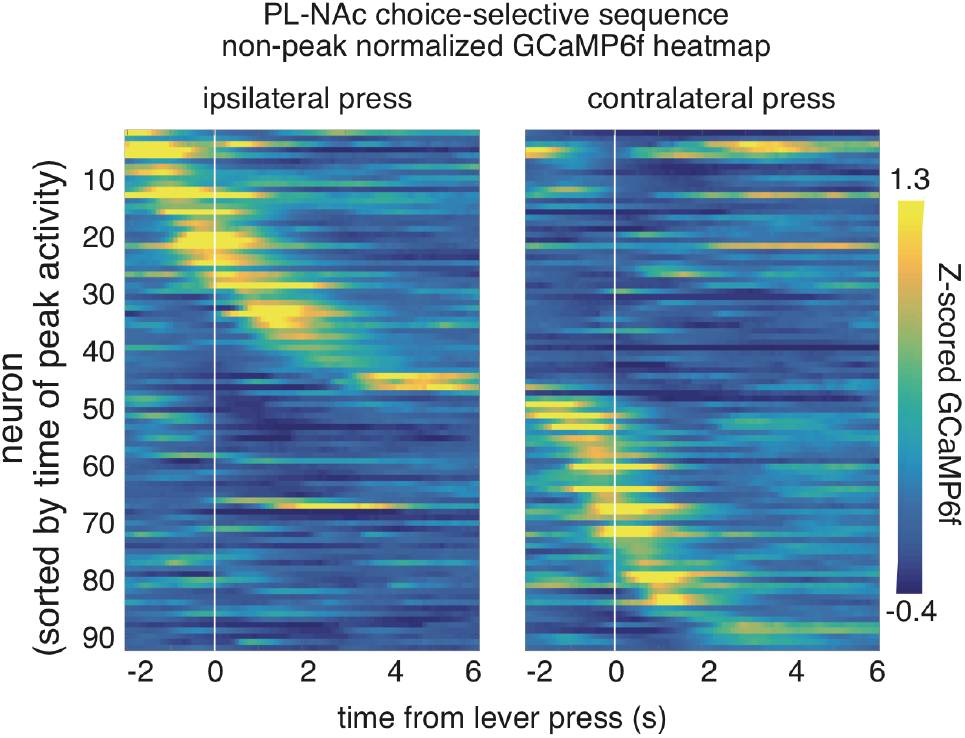
Choice-selective sequences in PL-NAc neurons without peak-normalization. Heatmap demonstrating sequential response of choice-selective PL-NAc neurons to the ipsilateral and contralateral lever press. Similar to **Figure 4b-c**, but time-locked, trial-averaged GCaMP6f fluorescence is not normalized by the peak response to the lever press and is taken from all trials.

**Supplementary Figure 6.**
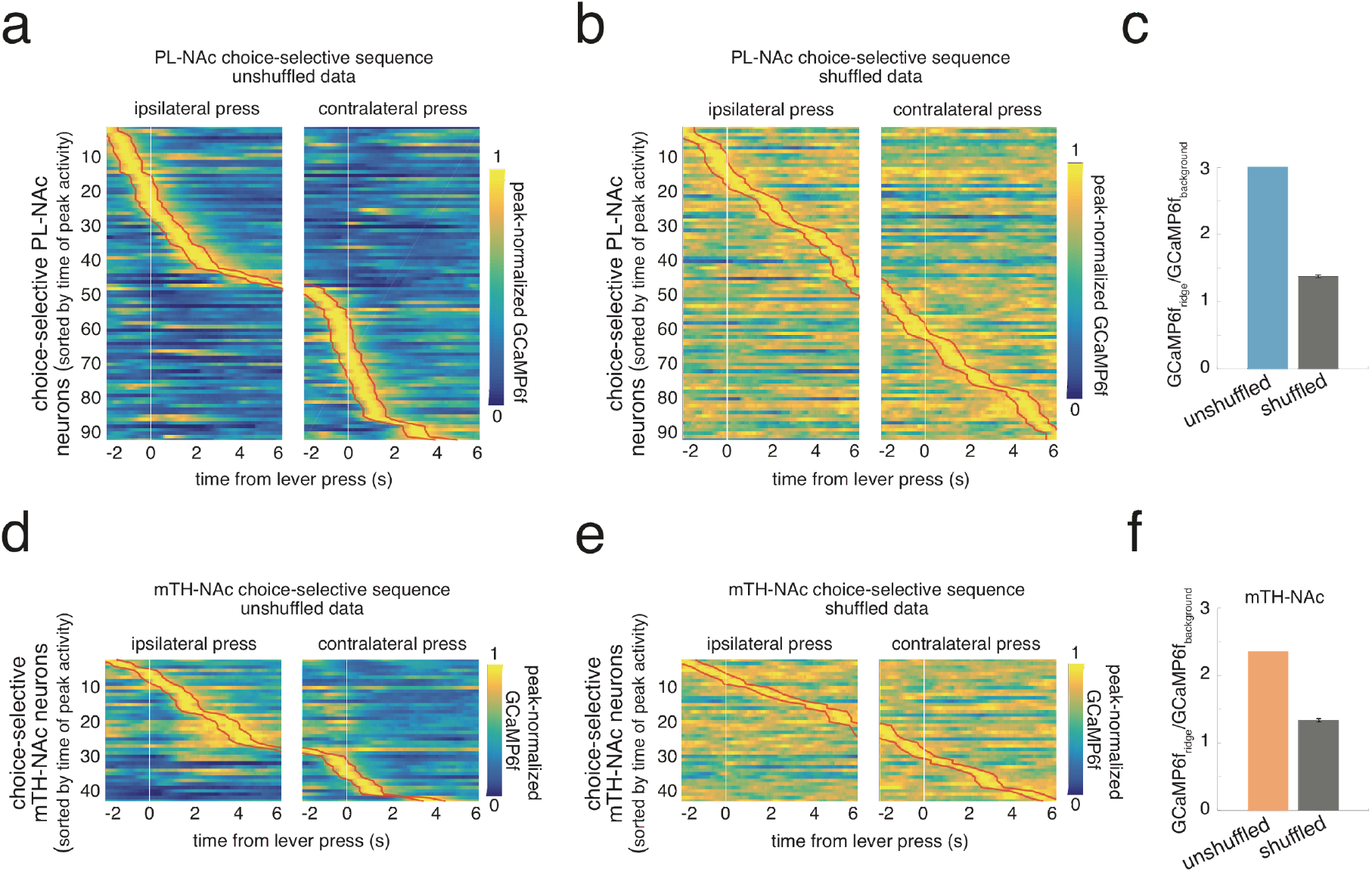
The calculated ridge-to-background ratio of PL-NAc neurons supports the presence of sequences. **(a)** Sequential activity of PL-NAc choice-selective neurons. Similar to **Figure 4b-c**, the heatmap is ordered by the time of peak activity time-locked to the ipsilateral (left column) and contralateral (right column) lever press of each neuron, but instead of cross-validation, activity is averaged across all trials. Red trace represents the borders of the one-second window around the peak defined as the ‘ridge’. Activity at all other surrounding timepoints is considered the ‘background’. **(b)** Same as **a** for data that is shuffled by temporally shifting the GCaMP6f fluorescence trace across a recording session separately for each neuron by a random number of frames, chosen from a uniform distribution. Ordering by the time of peak activity generates spurious sequential activity across the diagonal in shuffled data. **(c)** Calculated ridge-to-background ratio of PL-NAc neurons using unshuffled (blue) and shuffled (grey) data. A ratio is calculated for each individual neuron and the average of these ratios across all neurons displayed in the heatmap is shown. The ratio calculated from unshuffled data is significantly larger than that from the shuffled data (P<0.0001, comparison between unshuffled data and distribution of 500 shuffled iterations). Error bars for shuffled data indicate one standard deviation. **(d-f)** Same as **a-c** but ridge-to-background is calculated using mTH-NAc neural recordings. Similar to PL-NAc, the ratio calculated from unshuffled data was significantly larger than that from the shuffled data (P<0.0001). However, when comparing across the populations, the ridge-to-background calculated using PL-NAc neurons (3.01+/-0.12, mean+/-sem, n=92 neurons) was significantly larger than that using mTH-NAc (2.36+/-0.12, mean+/-sem, n=42 neurons; P=0.001: unpaired, two-tailed t-test comparing ratio between PL-NAc and mTH-NAc neurons, n=92 and 42 for PL-NAc and mTH-NAc populations, respectively).

**Supplementary Figure 7.**
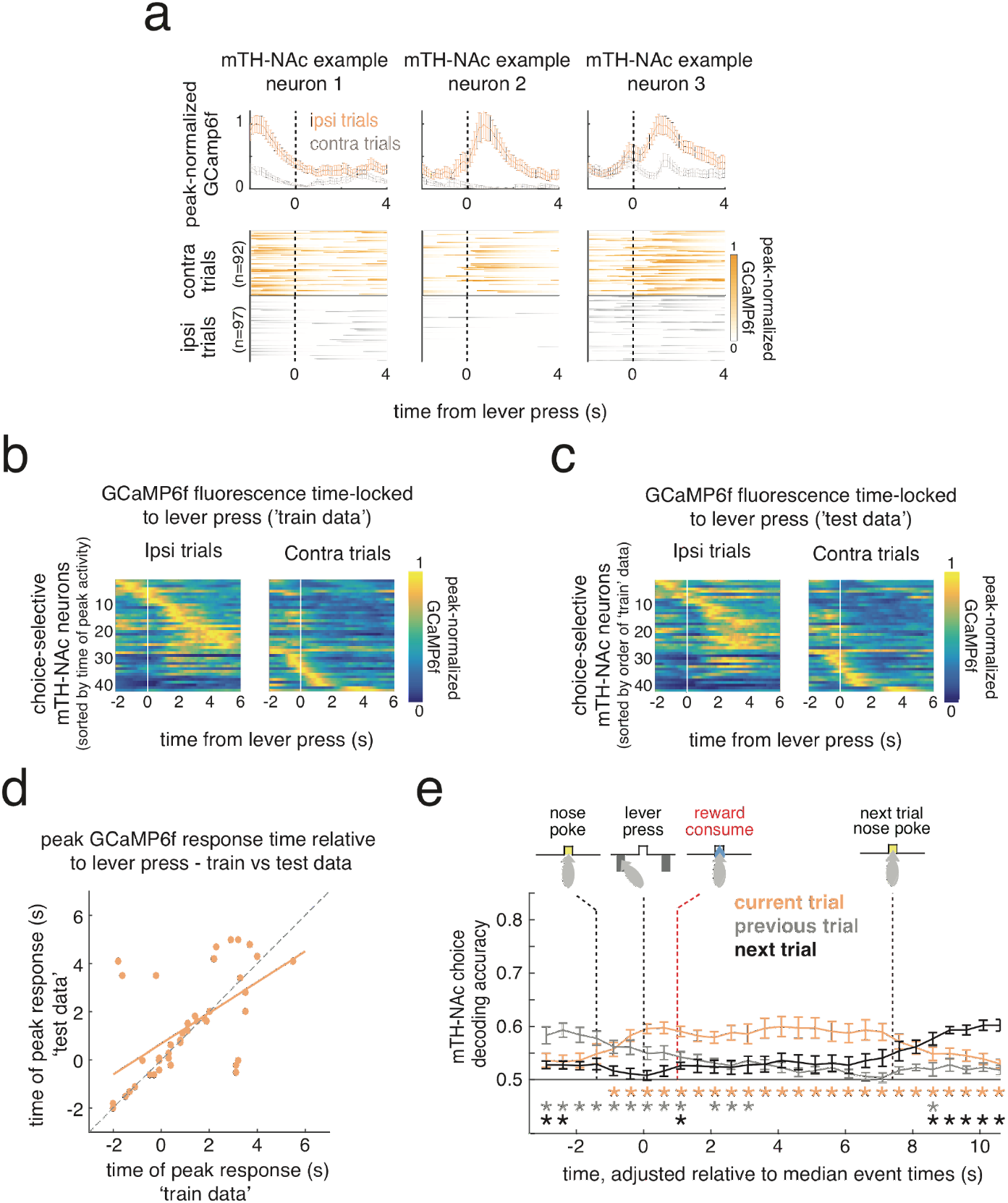
mTH-NAc choice-selective neurons display sequential activity that is less consistent than PL-NAc. **(a)** Top; average GCaMP6f fluorescence of three simultaneously imaged mTH-NAc choice-selective neurons with different response times relative to the lever press. Error bars are s.e.m across trials. Bottom, heatmaps of GCaMP6f fluorescence response across trials to ipsilateral (orange) and contralateral (grey) lever presses. **(b,c)** Heatmaps of choice-selective mTH-NAc neurons’ peak-normalized GCaMP6f responses to lever press (n=42/256 neurons). Each row is the average GCaMP6f fluorescence time-locked to the ipsilateral (left column) and contralateral (right column) lever press for a neuron, normalized by the neuron’s peak average fluorescence. In **b** (‘train data’), heatmap is generated using a randomly selected half of trials and ordered by the time of each neuron’s peak activity. In **c** (‘test data’), the peak-normalized, time-locked GCaMP6f fluorescence from the other half of trials was used while maintaining the order from ‘train data’ in **b**. Compare to PL-NAc data in **Figure 4b-c**. **(d)** Correlation between the time of peak activity using the ‘train’ (horizontal axis) and ‘test’ (vertical axis) trials for choice-selective mTH-NAc neurons. While mTH-NAc choice-selective neurons also show significant correlation between ‘train’ and ‘test’ trials (R^2^ = 0.59, P = 4.2×10^-5^, n=42 neurons from 9 mice), this correlation is significantly lower than that of PL-NAc (comparison with data in **Figure 4d**; P=0.0096, Z=2.34, Fisher’s R-to-Z transformation, comparison of correlation coefficients derived from comparing peak activity between ‘test’ and ‘training’ data from PL-NAc versus mTH-NAc). **(e)** Average choice decoding accuracy of the mice’s choice on the current (orange), previous (grey) and next trial (black) as a function of GCaMP6f fluorescence throughout the current trial. GCaMP6f fluorescence is taken from 100 random selections per mouse of 10 simultaneously imaged mTH-NAc neurons (each trial’s activity is adjusted in a piecewise linear manner relative to the median time of the nose poke, lever press and next trial nose poke, see Methods for details). Error bars are s.e.m. across mice (n=9 mice). Red dashed line indicates median onset of reward consumption. Asterisks indicate significant decoding accuracy above chance, P<0.01, two-tailed, one-sample t-test across mice.

**Supplementary Figure 8.**
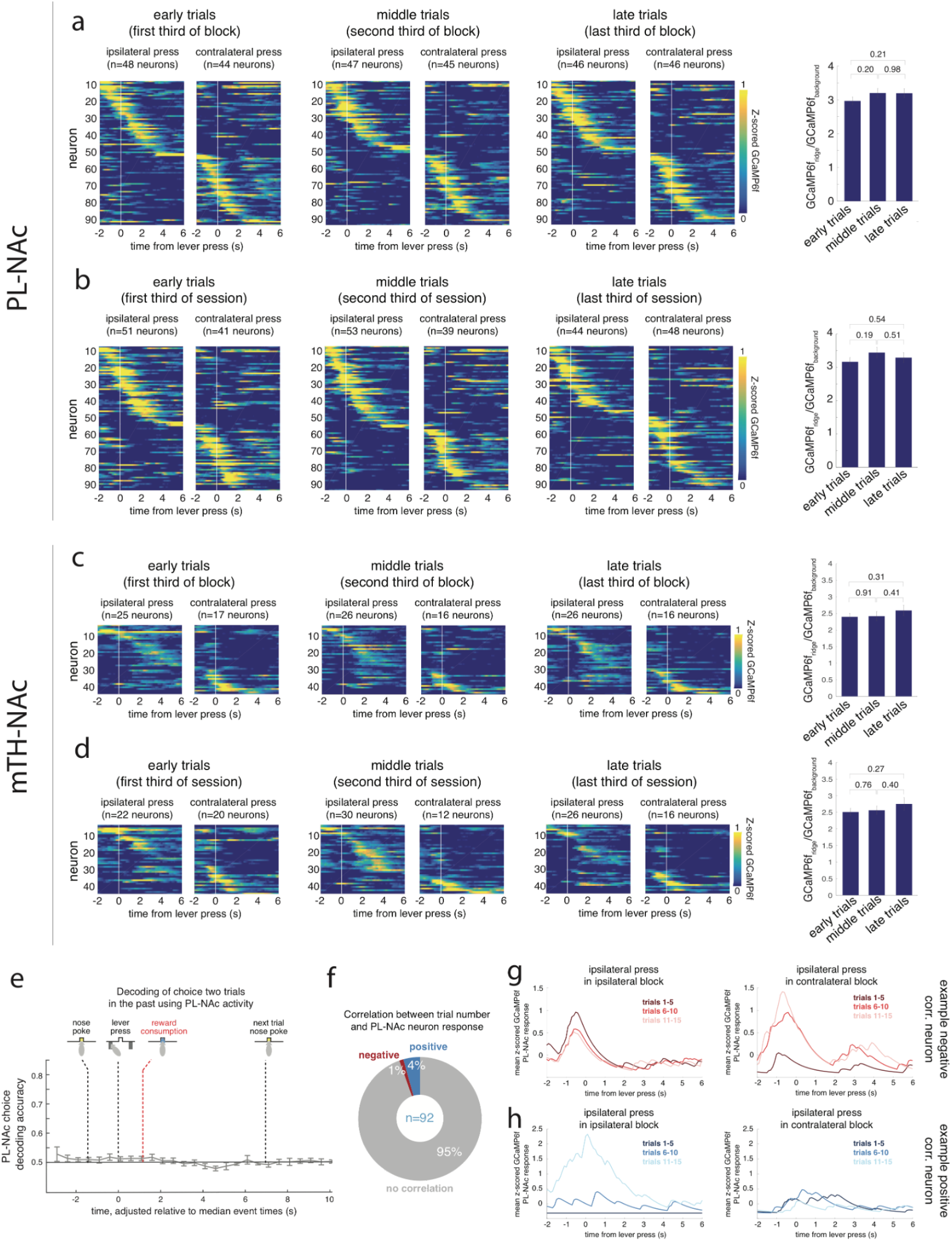
Trial number within a block, and two-trial-back choice, are not strongly encoded by PL-NAc activity. **(a)** Left, heatmaps of Z-scored GCaMP6f activity from 92 choice-selective PL-NAc neurons averaged across the first, middle and last third of trials of each block. Right, calculated ridge-to-background ratio derived from average activity from each third of trials within a block. Error bars represent s.e.m. across neurons (n=92). No significant changes are observed in the ratios calculated across a block, suggesting that the strength of sequences is not modulated by block trial number (p>0.05, paired, two-tailed t-test). **(b)** Same as **a** except data is split into the first, middle and final third of the entire recording session. **(c,d)** Same as **a,b** except activity is of mTH-NAc choice-selective neurons. **(e)** Decoding accuracy for the mice’s choice two trials back using activity from 10 simultaneously recorded PL-NAc neurons. Unlike choice decoding on the current and previous trial (**Figure 4e**; blue and black traces, respectively), PL-NAc activity is notableto accurately decode choice from two trials back after correcting for cross-trial choice correlations (see Methods for details) at any time point in the trial (p>0.05 for all time points: one-sample, two-tailed t-test across mice comparing decoding accuracy with chance rate of 0.5). **(f)** Proportion of PL-NAc choice-selective neurons whose activity is significantly positively (blue, n=4 neurons) or negatively (red, n=1 neuron) correlated with the number of trials into a block (P<0.01). Significance was determined by comparing the calculated correlation coefficient of each neuron to a null distribution of 500 correlation coefficients generated using GCaMP6f signal circularly shifted by a random integer, to control for slow drift in the data. R-values were calculated using the maximum GCaMP6f activity from 2s before to 6s after the time of lever press for the first 15 trials in a block. Using the same criteria, no mTH-NAc neurons were significantly correlated, either positively or negatively with trial number in a block (p>0.05). **(g)** Left, average activity of the negatively correlated PL-NAc neuron in response to an ipsilateral lever press at various trials in an ipsilateral block, where the ipsilateral lever had a higher probability of reward and, thus, the value of the ipsilateral lever increases as a function of trial number. Right, average activity of the negatively correlated PL-NAc neuron in response to an ipsilateral press in a contralateral block. **(h)** Same as **g** except for an example of a positively correlated PL-NAc neuron.

**Supplementary Figure 9.**
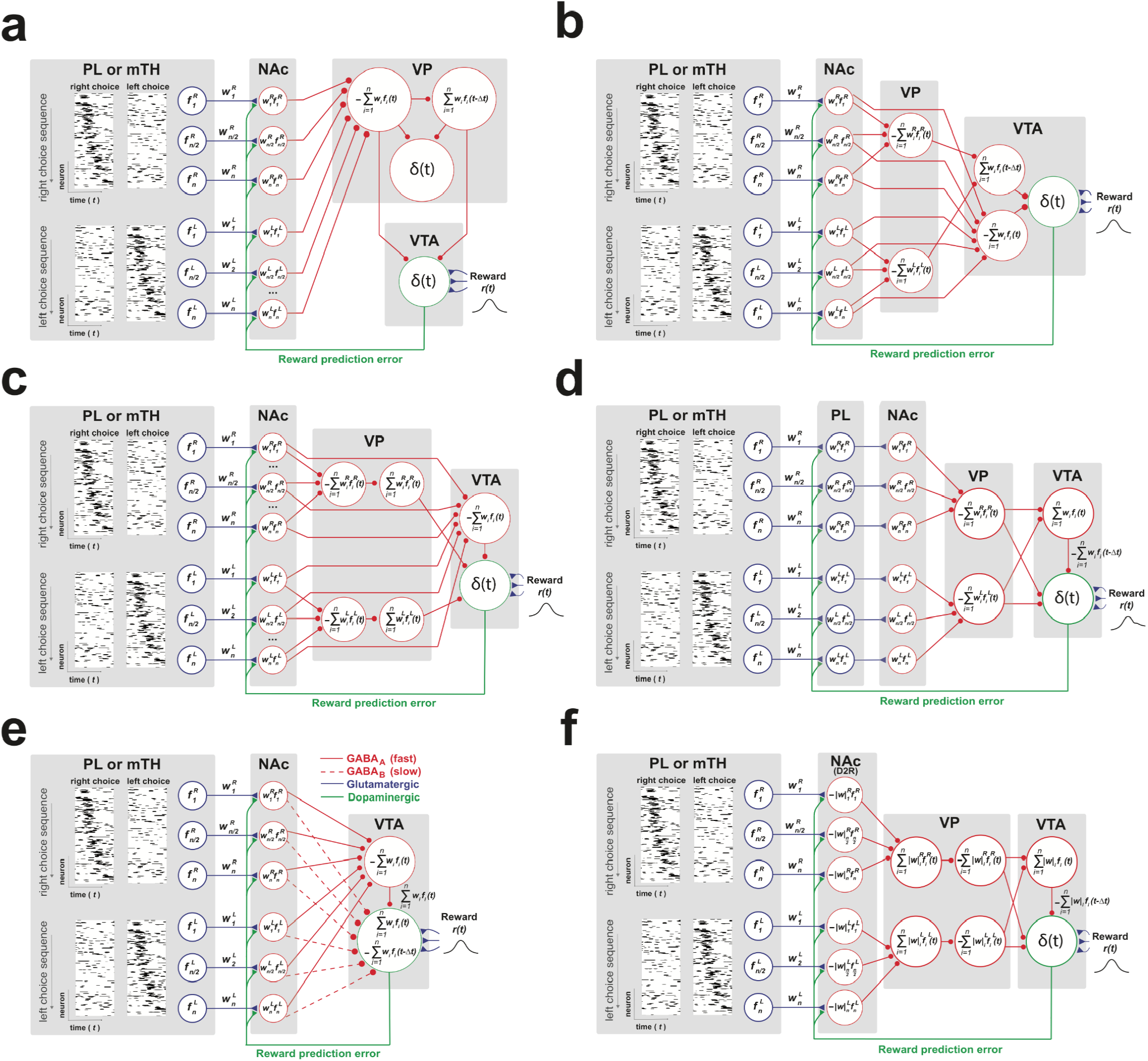
Alternative model architectures used to implement synaptic plasticity model. **(a-f)** Alternative models constructed using known circuit architecture. All models except **a** are able to correctly generate an RPE signal by providing a ‘fast excitatory’ and ‘slow-inhibitory’ value pathway to the VTA dopamine neuron population. Note that all model variants rely on choice-selective sequences in PL-NAc to bridge actions and outcomes across time. For all model variants, GABAergic, glutamatergic and dopaminergic projections are denoted as red, blue and green, respectively. Brain region abbreviations are: prelimbic cortex, PL; nucleus accumbens, NAc; ventral pallidum, VP; ventral tegmental area, VTA. **(a)** In this model, the delay and inversion of the value signal is accomplished through a second VP neuron. These two VP neurons converge onto a third VP interneuron to generate an RPE signal in VP, as has been observed by Ottenheimer et al. (2020). **(b)** In this model, the fast excitatory pathway is generated via a direct projection of NAc neurons onto the VTA GABA neuron while the slow inhibitory pathway passes through the VP before synapsing onto a VTA GABA neuron. **(c)** Similar to **b** except that the slow inhibitory pathway contains an additional VP neuron, which accomplishes the sign inversion and delay assigned to a VTA GABA neuron in **b**. Since the models in **b** prescribe a role for the observed NAc-D1R projections to VTA GABA neurons, they produce negative value signals in VTA GABA neurons, whereas only positive value signals have been observed experimentally in identified GABA interneurons in the VTA (Cohen et al., 2012). **(d)** An alternative method to generate sequential value signals in our model is to modulate the strength of synaptic inputs to PL-NAc, as opposed to PL-NAc outputs, via dopaminergic inputs to the PL. This may be supported by recent studies which demonstrated rapid, DA-modulated synaptic plasticity of PL-PL synapses (Young and Yang, 2005). **(e)** To account for previous work describing direct projections from NAc D1R neurons to the VTA (Beier et al., 2015; Watabe-Uchida et al., 2012; Yang et al., 2018), this alternative model architecture has NAc neurons projecting directly to the VTA, skipping the VP. In this model, the timing difference needed to compute an RPE signal is generated through the activity of fast ionotropic GABA-A receptors (solid red trace), which have been shown to preferentially express in NAc-VTA GABA interneuron projection postsynaptic densities (Edwards et al., 2017), while activity of metabotropic GABA-B receptors (dashed red trace), which are preferentially expressed in the postsynaptic densities of NAc-VTA DA projections (Edwards et al., 2017), generate the slow-inhibitory pathway. Notably, without this differential expression of GABA receptors in the DAergic and GABAergic populations of the VTA, this model architecture would fail to produce an RPE signal, as it would instead generate a fast-inhibitory and slow-excitatory signal in the VTA DA neuron population. **(f)** Multiple studies have implicated D2-R expressing MSNs as playing a critical role in reversal learning in multiple mammalian species (Boulougouris et al., 2009; DeSteno and Schmauss, 2009; Eisenegger, 2016; Kruzich and Grandy, 2004; Kruzich et al., 2006; Kwak et al., 2014; Piray, 2011). Thus, in this model we account for the possibility that the reversal behavior in our task is mediated specifically by changes to synaptic weights from PL to D2-R-expressing NAc MSNs. This model assumes the opposite dopamine-mediated plasticity rule (LTD rather than LTP) than the previous models.

**Supplementary Figure 10.**
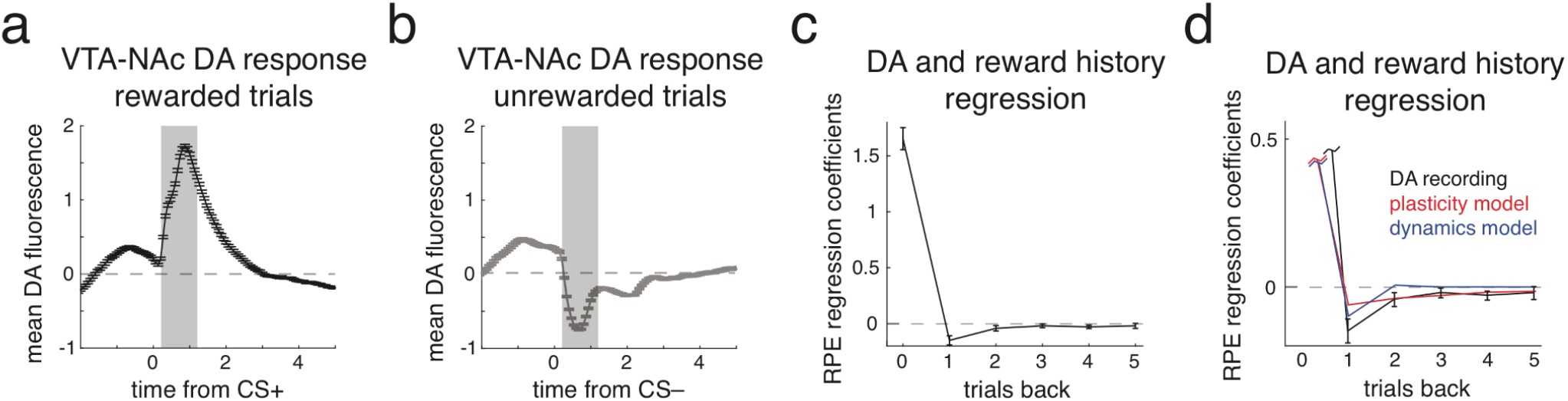
Reward prediction error (RPE) encoding observed in recorded dopamine (DA) activity is similar to that produced by our synaptic plasticity and neural dynamics models. **(a)** Mean bulk GCaMP6f fluorescence from VTA-NAc DA terminals in response to a conditioned stimulus signaling reward (CS+, data taken from Parker et al., 2016). Note that terminal fluorescence recordings are presented here to more accurately reflect the signal that downstream NAc neurons are receiving in our model. **(b)** Same as **a** except DA fluorescence in response to the conditioned stimulus signaling an unrewarded trial (CS-). **(c)** Coefficients from a multiple linear regression in which outcome is predicted using mean DA fluorescence signals from 0.2-1.2s relative to the time of CS presentation across current (“0”) and multiple previous trials (see shaded region in **a,b**), similar to **Figure 5g,l,q** and **Figure 7d**. The positive coefficient for the current trial and negative coefficients for previous trials indicate the encoding of an RPE. Error bars represent s.e.m across 11 recording sites. **(d)** Same as **c**, but also including coefficients from the synaptic plasticity model (red, same coefficients as **Figure 5g**) and the neural dynamics model (black, same coefficients as **Figure 7d**), to allow direct comparison.

**Supplementary Figure 11.**
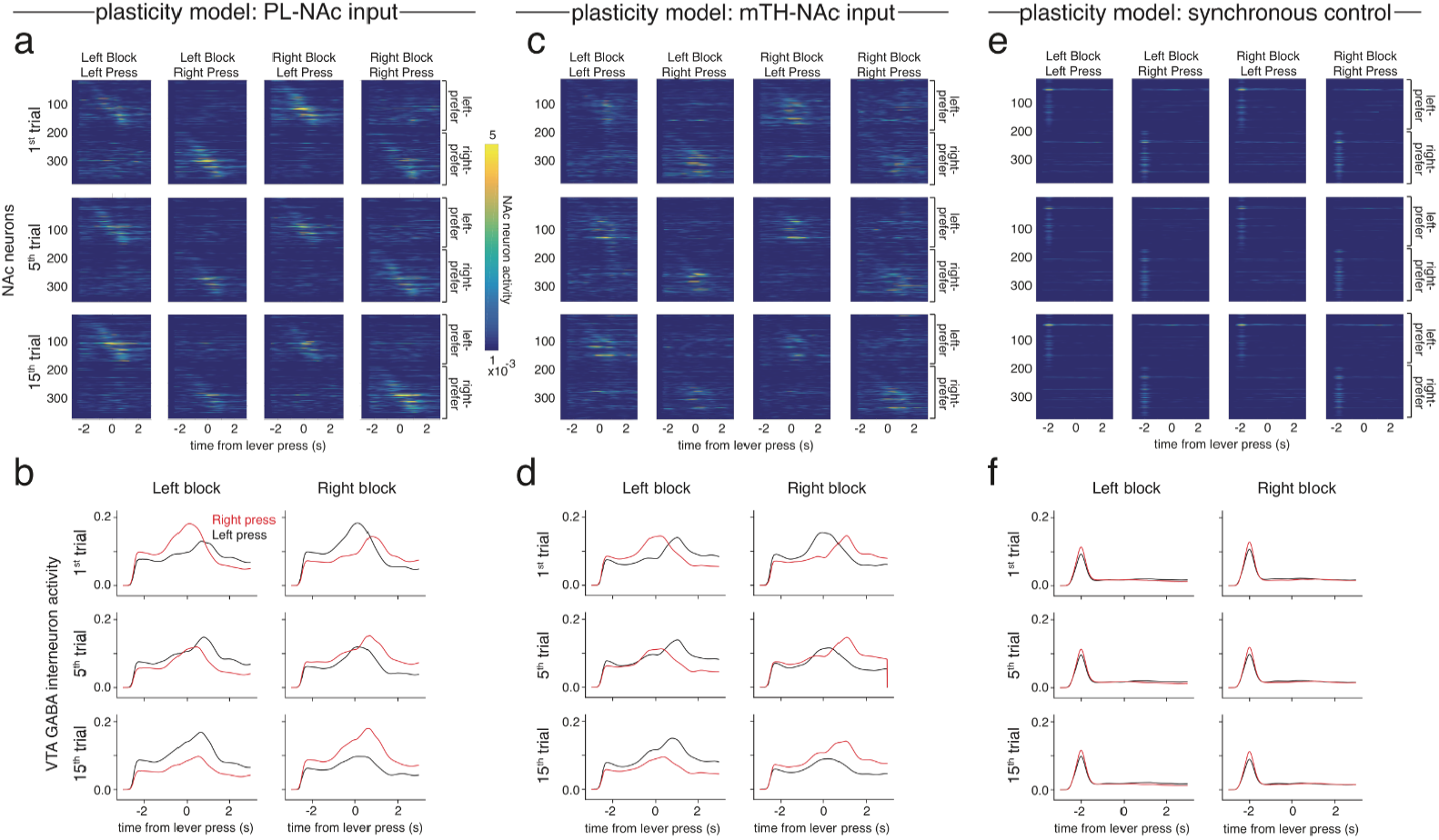
Synaptic plasticity model using sequential but not synchronous PL-NAc activity correctly modulates activity in NAc projection neurons and VTA GABA interneurons. **(a)** Heatmaps of average activity relative to the time of the lever press for NAc projection neurons in the PL-NAc model (**Figure 5c**). Top, middle and bottom heatmaps are the average activity across the first, fifth and fifteenth trial of each block, respectively. Each column is the average activity across trials from different block/press combinations. For each subplot, neurons 1-184 are left-preferring and neurons 185-368 are right-preferring. The activity of these left- and right-preferring NAc neurons increases throughout a block of their respective lever preference. In contrast, their activity decreases throughout a block opposite to their lever preference. **(b)** Average activity of VTA GABA interneuron from synaptic plasticity model using PL-NAc activity as input on left (black) or right (red) trials. Activity is relative to the time of the lever press across the first, fifth and fifteenth trials of a left-preferring (left column) or right-preferring (right column) block. Similar to **a**, throughout a left block (left column), the activity on left press trials increases from the first to fifteenth trial while the activity on right presses decreases. The opposite pattern is seen for left and right press trials throughout a right block (right column). **(c,d)** Same as **a,b** except NAc and VTA GABA interneuron generated using mTH-NAc as input to the synaptic plasticity model. **(e,f)** Same as **a,b** except NAc and VTA GABA interneuron from the PL-NAc synchronous control synaptic plasticity model.

**Supplementary Figure 12.**
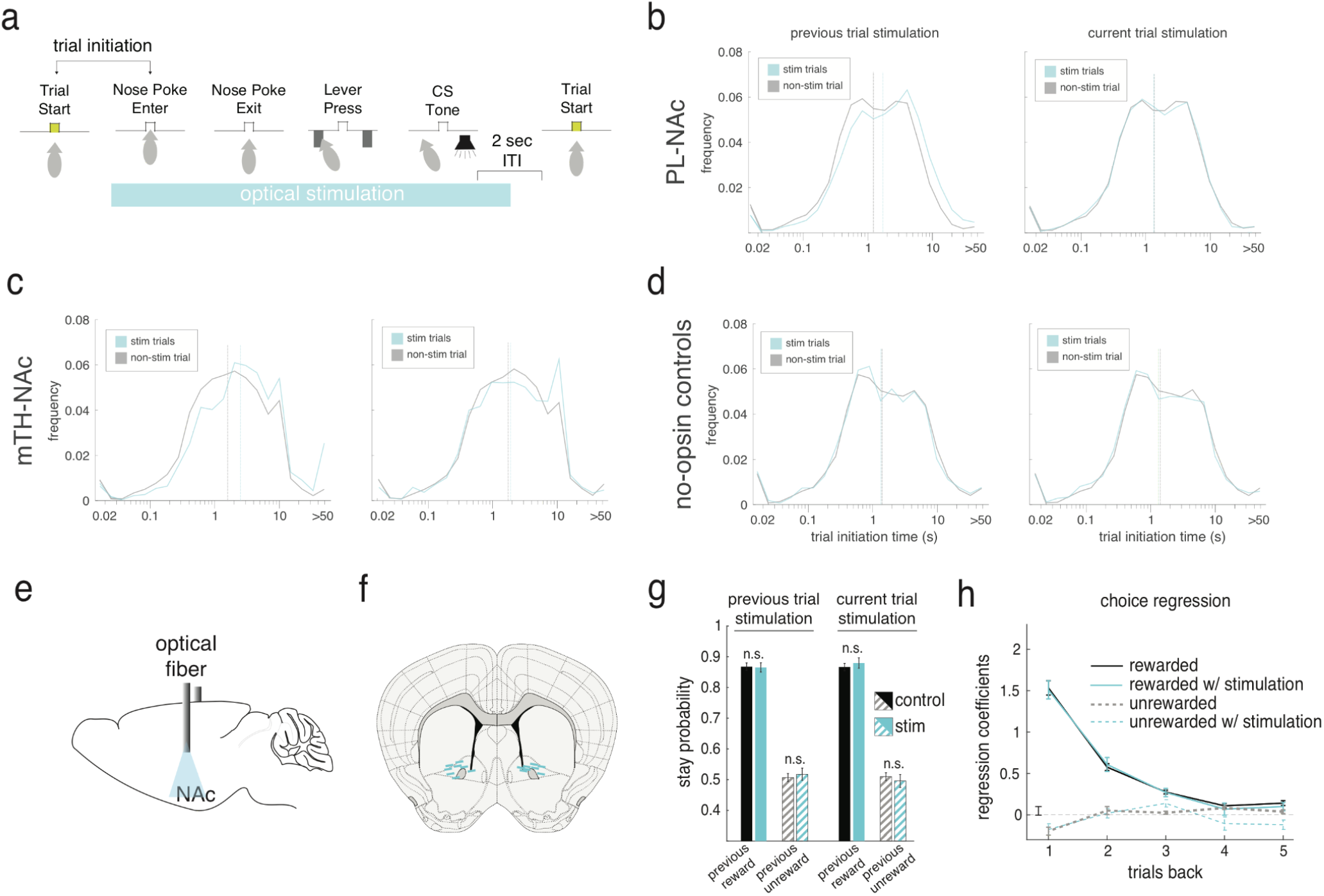
Laser stimulation affects trial initiation times in both PL-NAc and mTH-NAc but does not affect behavior in control mice that do not express opsin. **(a)** Schematic of trial structure and time of optical stimulation. “Trial initiation” time is defined as the latency between the start of the trial and the mouse entering the central nose poke. **(b)** Left, distribution of trial initiation times following stimulation trials (blue) and non-stimulation trials (grey) in PL-NAc ChR2 mice. Blue and grey vertical lines indicate median initiation times for stim and non-stim trials, respectively. Right, same only effect of current trial stimulation on trial initiation times. Previous trial stimulation resulted in significantly longer trial initiations in PL-NAc ChR2 mice than no-opsin control mice (P=3.95×10^-6, p-value from the ‘opsin group X previous trial stimulation’ interaction term of a mixed effects model used to predict latency times of PL-NAc and control mice, fit using the fitglme function in MATLAB; see Methods for additional model details). In contrast, current trial stimulation had no significant effect on initiation times (P=0.16, same test as above except p-value is that of the interaction term of ‘opsin group X current trial stimulation’), an expected result as the start of stimulation was contingent on the mouse performing a nose poke. **(c)** mTH-NAc ChR2 mice had significantly longer trial initiation times following optical stimulation than no-opsin control mice (P=2.74×10^-3, p-value from the ‘opsin group X previous trial stimulation’ interaction term of a mixed effects model used to predict latency times of mTH-NAc and control mice) but no effect of current trial stimulation was observed (P=0.07, same test as above except the p-value is that from the ‘opsin group X current trial stimulation’ interaction term). **(d)** Same as **b,c** except latencies from no-opsin control cohort. **(e)** Surgical schematic of no-opsin control cohort. Optical fibers were implanted into the NAc. **(f)** Optical fiber tip locations of no-opsin control cohort (n=8 mice). **(g)** Unlike PL-NAc ChR2 expressing mice (**Figure 6f,g**), neither current nor previous trial stimulation changed the stay probability in control mice following rewarded (P=0.52: previous trial stimulation; P=0.58: current trial stimulation; paired t-test) or unrewarded trials (P=0.24: previous trial stimulation; P=0.47: current trial stimulation; paired t-test). **(h)** Likewise, stimulation from multiple trials back had no effect on choice (P>0.05 for all trials back, t-test across mice’s laser x choice interaction term coefficients).

**Supplementary Figure 13.**
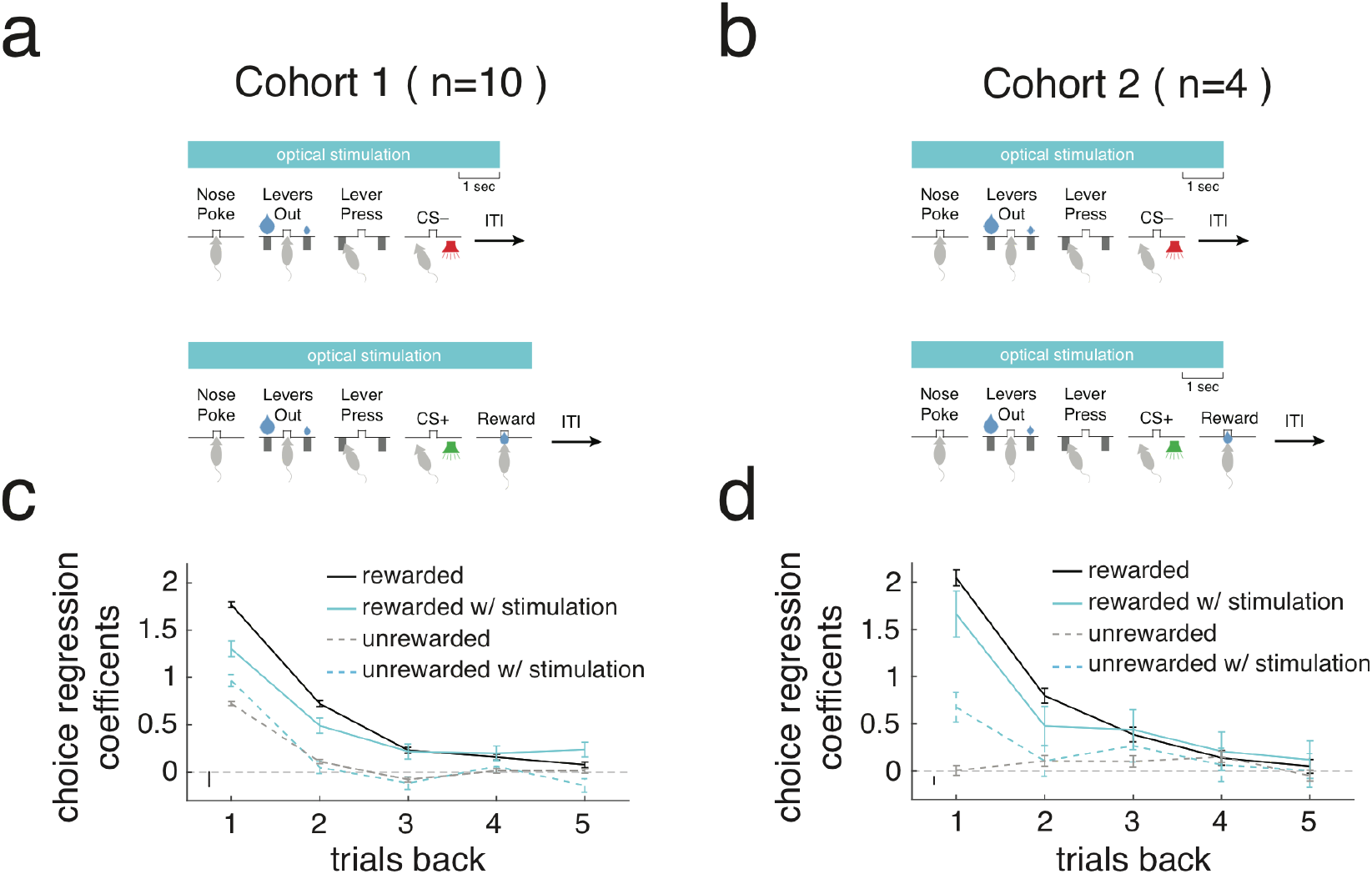
Effect of PL-NAc optogenetic stimulation in two cohorts. **(a)** Schematic of optical stimulation parameters for cohort 1. On 10% of unrewarded trials, optical stimulation began when the mouse entered the central nosepoke and ended 1s into the intertrial interval (ITI), which began at the end of the 500ms CS-tone. On 10% of rewarded trials, stimulation began with nose poke and ended after the mouse left the reward port. **(b)** Schematic for cohort 2. Unlike cohort 1, optical stimulation ended on the same timescale on both rewarded and unrewarded trials, 1s after the end of CS presentation. **(c)** Logistic regression model similar to that in **Figure 1e** demonstrating the effect of PL-NAc stimulation on lever choice in cohort 1 mice (n=10 mice, see Methods for model details). Rewarded trials with stimulation one and two trials back decreased stay probability compared with rewarded trials without stimulation. Stimulation had an opposite effect on unrewarded trials, for which there was an increase in stay probability following stimulation one trial back compared to trials without stimulation. **(d)** Same as **c** except data from cohort 2 (n=4 mice). Effect of optical stimulation of PL-NAc neurons was qualitatively similar across the two cohorts.

**Supplementary Figure 14.**
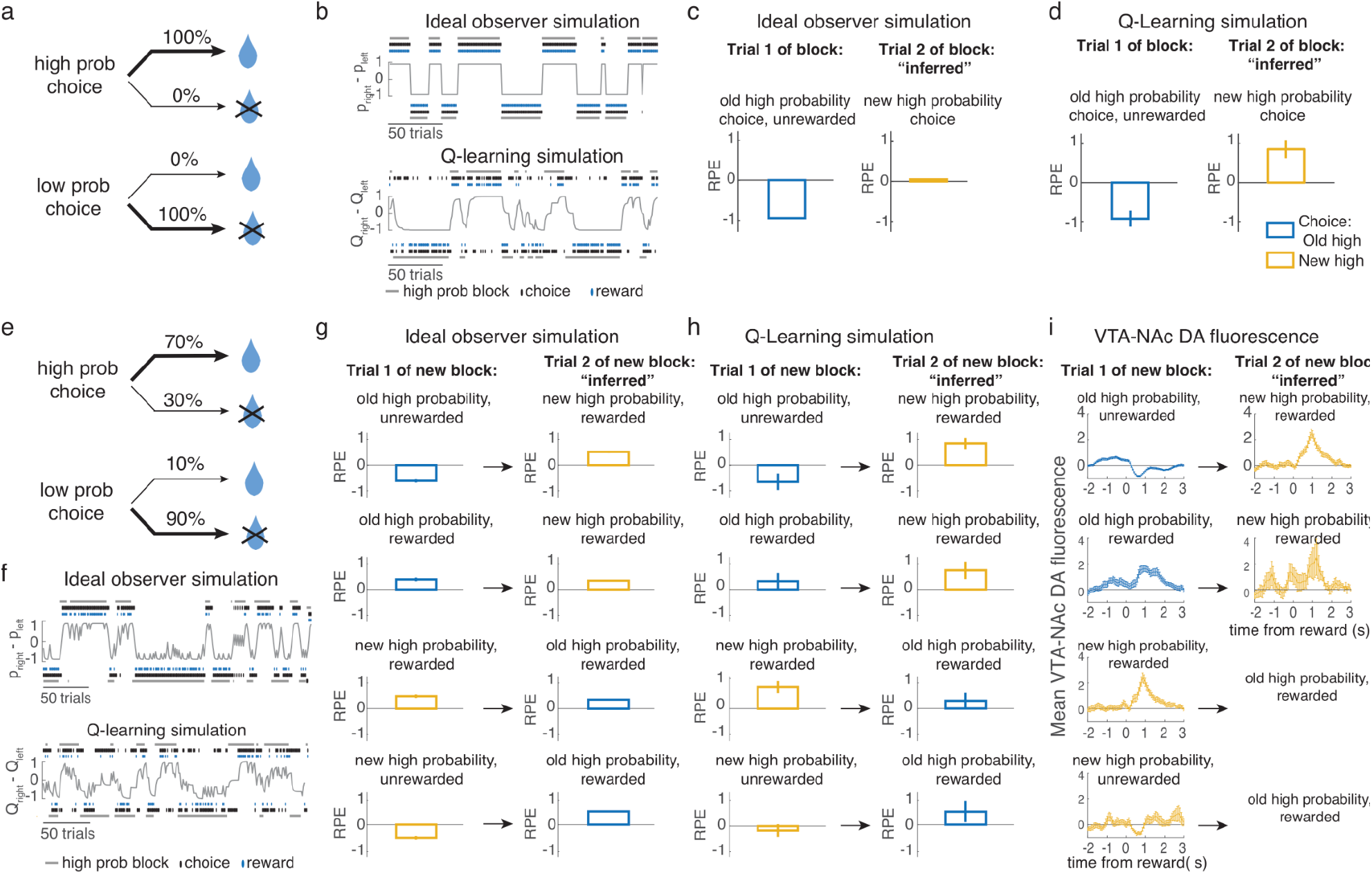
Similar RPE signatures for ideal observer and Q-learning simulation during reversal learning. Given previous evidence that dopamine signals can reflect knowledge of task structure (Bromberg-Martin et al., 2010; Sadacca et al., 2016), we used modeling to gain insight into how clearly RPE in the probabilistic reversal learning task (a bandit task) can indicate the use of model-based inference of block reversals for different reward probability structures. This was done by simulating task performance using an ideal observer model with knowledge of the block structure, and a Q-learning model which did not have information about the block structure (see Methods for more details). **a-d)** To confirm that our ideal observer simulation captured a previously reported RPE signature of model-based block reversal inference, we first simulated behavior of 100,000 trials of a task similar to that used in Bromberg-Martin et al. (2010). **(a)** In this task, one option was rewarded 100% of the time while the other was never rewarded, and the identity of the high probability choice randomly reversed with a probability of 0.05 on each trial. (**b)** Example performance of the ideal observer simulation (top) and the Q-learning simulation (bottom). Choice is determined by the difference between expected reward for the available actions, *ρ*, for the ideal observer and the difference between the action values, *Q*, for the Q-learning simulation (these values are plotted in gray). **(c)** To evaluate RPE signatures of model-based block reversal inference, we compared the estimated RPE (experienced reward minus expected reward for the chosen action) on trial 1 and trial 2 of the new block. The RPE on trial 2 was low for the high probability choice in the new block even without direct experience of that action-outcome pairing. This means that the ideal observer infers the block reversal, so the new, not yet experienced reward contingency is expected and the RPE is low. **(d)** In contrast, because the Q-learning model only updates the value of the chosen action, on trial 2, when the simulation is rewarded for the previously low-probability choice, the reward remains unexpected and the RPE is high. **e-i)** Simulated performance of 100,000 trials using the reward probabilities from this study. **(e)** The high probability action was rewarded 70% of the time while the low probability action was rewarded 10% of the time and the blocks reversed according to the same rule as in **a**. **(f)** Example performance of the ideal observer (top) and the Q-learning (bottom) simulations in this task. **(g)** To determine whether there is a strong qualitative RPE signature of block reversal inference in this task, we compared RPE on the 4 possible trial-1 types to RPE on the subsequent rewarded switch trials (i.e. choice on trial 2 was different than trial 1, meaning that any changes in RPE must be inferred). We focus on rewarded trials to aid comparison with reward responses recorded in dopamine terminals during this task (Parker et al., 2016). In this case, inference of the block reversal is not obviously reflected in the RPE, since the RPE for a given action on trial 1 and trial 2 are similar (comparing the same color bars for rewarded actions on trials 1 and 2). This is because, even though the ideal observer updates the predicted reward for both the chosen and unchosen actions, when reward delivery is probabilistic, predicted reward remains moderate for both actions and RPE changes only subtly. **(h)** Same as in **g** for the Q-learning simulation. As expected, RPE looks very similar on trial 1 and trial 2 for a given rewarded action because the Q-learning simulation does not update the value of the unchosen action on trial 1. **(i)** Consistent with the results from both simulations, GCaMP6f zscored dF/F from dopaminergic axons in the NAc recorded in Parker et al. (2016) is also very similar for a given rewarded action on trial 1 and trial 2. Note that the mice did not make all possible choices in this task, so some trial types are missing. The simulations also rarely made these choices (e.g., switch following a rewarded new high probability choice).

## References

Aggarwal, M., Hyland, B.I., and Wickens, J.R. (2012). Neural control of dopamine neurotransmission: implications for reinforcement learning. Eur. J. Neurosci. 35, 1115–1123.

Akhlaghpour, H., Wiskerke, J., Choi, J.Y., Taliaferro, J.P., Au, J., and Witten, I.B. (2016). Dissociated sequential activity and stimulus encoding in the dorsomedial striatum during spatial working memory. Elife 5, e19507.

Apicella, P., Ljungberg, T., Scarnati, E., and Schultz, W. (1991). Responses to reward in monkey dorsal and ventral striatum. Experimental Brain Research 85.

Asaad, W.F., Lauro, P.M., Perge, J.A., and Eskandar, E.N. (2017). Prefrontal Neurons Encode a Solution to the Credit-Assignment Problem. The Journal of Neuroscience 37, 6995–7007.

Atallah, H.E., Lopez-Paniagua, D., Rudy, J.W., and O’Reilly, R.C. (2007). Separate neural substrates for skill learning and performance in the ventral and dorsal striatum. Nat. Neurosci. 10, 126–131.

Bagot, R.C., Parise, E.M., Peña, C.J., Zhang, H.-X., Maze, I., Chaudhury, D., Persaud, B., Cachope, R., Bolaños-Guzmán, C.A., Cheer, J.F., et al. (2015). Ventral hippocampal afferents to the nucleus accumbens regulate susceptibility to depression. Nat. Commun. 6, 7062.

Bayer, H.M., and Glimcher, P.W. (2005). Midbrain Dopamine Neurons Encode a Quantitative Reward Prediction Error Signal. Neuron 47, 129–141.

Beier, K.T., Steinberg, E.E., DeLoach, K.E., Xie, S., Miyamichi, K., Schwarz, L., Gao, X.J., Kremer, E.J., Malenka, R.C., and Luo, L. (2015). Circuit Architecture of VTA Dopamine Neurons Revealed by Systematic Input-Output Mapping. Cell 162, 622–634.

Botvinick, M., Ritter, S., Wang, J.X., Kurth-Nelson, Z., Blundell, C., and Hassabis, D. (2019). Reinforcement Learning, Fast and Slow. Trends in Cognitive Sciences 23, 408–422.

Botvinick, M., Wang, J.X., Dabney, W., Miller, K.J., and Kurth-Nelson, Z. (2020). Deep Reinforcement Learning and Its Neuroscientific Implications. Neuron 107, 603–616.

Boulougouris, V., Castañé, A., and Robbins, T.W. (2009). Dopamine D2/D3 receptor agonist quinpirole impairs spatial reversal learning in rats: investigation of D3 receptor involvement in persistent behavior. Psychopharmacology 202, 611–620.

Britt, J.P., Benaliouad, F., McDevitt, R.A., Stuber, G.D., Wise, R.A., and Bonci, A. (2012). Synaptic and Behavioral Profile of Multiple Glutamatergic Inputs to the Nucleus Accumbens. Neuron 76, 790–803.

Brog, J.S., Salyapongse, A., Deutch, A.Y., and Zahm, D.S. (1993). The patterns of afferent innervation of the core and shell in the “accumbens” part of the rat ventral striatum: immunohistochemical detection of retrogradely transported fluoro-gold. J. Comp. Neurol. 338, 255–278.

Bromberg-Martin, E.S., Matsumoto, M., Hong, S., and Hikosaka, O. (2010). A pallidus-habenula-dopamine pathway signals inferred stimulus values. J. Neurophysiol. 104, 1068–1076.

Cador, M., Robbins, T.W., and Everitt, B.J. (1989). Involvement of the amygdala in stimulus-reward associations: interaction with the ventral striatum. Neuroscience 30, 77–86.

Cameron, C.M., Murugan, M., Choi, J.Y., Engel, E.A., and Witten, I.B. (2019). Increased Cocaine Motivation Is Associated with Degraded Spatial and Temporal Representations in IL-NAc Neurons. Neuron 103, 80–91.e7.

Campus, P., Covelo, I.R., Kim, Y., Parsegian, A., Kuhn, B.N., Lopez, S.A., Neumaier, J.F., Ferguson, S.M., Solberg Woods, L.C., Sarter, M., et al. (2019). The paraventricular thalamus is a critical mediator of top-down control of cue-motivated behavior in rats. Elife 8.

Cardinal, R.N., and Cheung, T.H.C. (2005). Nucleus accumbens core lesions retard instrumental learning and performance with delayed reinforcement in the rat. BMC Neurosci. 6, 9.

Carelli, R.M., King, V.C., Hampson, R.E., and Deadwyler, S.A. (1993). Firing patterns of nucleus accumbens neurons during cocaine self-administration in rats. Brain Research 626, 14–22.

Carrillo-Reid, L., Tecuapetla, F., Tapia, D., Hernández-Cruz, A., Galarraga, E., Drucker-Colin, R., and Bargas, J. (2008). Encoding network states by striatal cell assemblies. J. Neurophysiol. 99, 1435–1450.

Centonze, D., Picconi, B., Gubellini, P., Bernardi, G., and Calabresi, P. (2001). Dopaminergic control of synaptic plasticity in the dorsal striatum. European Journal of Neuroscience 13, 1071–1077.

Chen, R., Puzerey, P.A., Roeser, A.C., Riccelli, T.E., Podury, A., Maher, K., Farhang, A.R., and Goldberg, J.H. (2019). Songbird ventral pallidum sends diverse performance error signals to dopaminergic midbrain. Neuron 103, 266–276.e4.

Cohen, J.Y., Haesler, S., Vong, L., Lowell, B.B., and Uchida, N. (2012). Neuron-type-specific signals for reward and punishment in the ventral tegmental area. Nature 482, 85–88.

Collins, A.G.E., and Cockburn, J. (2020). Beyond dichotomies in reinforcement learning. Nat. Rev. Neurosci. 21, 576–586.

Collins, A.L., Aitken, T.J., Huang, I.-W., Shieh, C., Greenfield, V.Y., Monbouquette, H.G., Ostlund, S.B., and Wassum, K.M. (2019). Nucleus accumbens cholinergic interneurons oppose cue-motivated behavior. Biol. Psychiatry 86, 388–396.

Cox, J., and Witten, I.B. (2019). Striatal circuits for reward learning and decision-making. Nature Reviews Neuroscience 20, 482–494.

Day, J.J., and Carelli, R.M. (2007). The nucleus accumbens and Pavlovian reward learning. The Neuroscientist 13, 148–159.

Day, J.J., Wheeler, R.A., Roitman, M.F., and Carelli, R.M. (2006). Nucleus accumbens neurons encode Pavlovian approach behaviors: evidence from an autoshaping paradigm. Eur. J. Neurosci. 23, 1341–1351.

Dayan, P., and Niv, Y. (2008). Reinforcement learning: the good, the bad and the ugly. Curr. Opin. Neurobiol. 18, 185–196.

DeSteno, D.A., and Schmauss, C. (2009). A role for dopamine D2 receptors in reversal learning. Neuroscience 162, 118–127.

Di Ciano, P., Cardinal, R.N., Cowell, R.A., Little, S.J., and Everitt, B.J. (2001). Differential involvement of NMDA, AMPA/kainate, and dopamine receptors in the nucleus accumbens core in the acquisition and performance of pavlovian approach behavior. J. Neurosci. 21, 9471–9477.

Ding, X., Qiao, Y., Piao, C., Zheng, X., Liu, Z., and Liang, J. (2014). N-methyl-D-aspartate receptor-mediated glutamate transmission in nucleus accumbens plays a more important role than that in dorsal striatum in cognitive flexibility. Front. Behav. Neurosci. 8, 304.

Doll, B.B., Simon, D.A., and Daw, N.D. (2012). The ubiquity of model-based reinforcement learning. Curr. Opin. Neurobiol. 22, 1075–1081.

Do-Monte, F.H., Minier-Toribio, A., Quiñones-Laracuente, K., Medina-Colón, E.M., and Quirk, G.J. (2017). Thalamic regulation of sucrose seeking during unexpected reward omission. Neuron 94, 388–400.e4.

Doshi-Velez, F., and Konidaris, G. (2016). Hidden Parameter Markov Decision Processes: A *Semiparametric Regression Approach for Discovering Latent Task Parametrizations*. IJCAI 2016, 1432–1440.

Doya, K. (2002). Metalearning and neuromodulation. Neural Netw. 15, 495–506.

Duan, Y., Schulman, J., Chen, X., Bartlett, P.L., Sutskeve, I. and Abbeel, P. (2016). RL^2: Fast reinforcement learning via slow reinforcement learning. arXiv 1611.02779

Edwards, N.J., Tejeda, H.A., Pignatelli, M., Zhang, S., McDevitt, R.A., Wu, J., Bass, C.E., Bettler, B., Morales, M., and Bonci, A. (2017). Circuit specificity in the inhibitory architecture of the VTA regulates cocaine-induced behavior. Nat. Neurosci. 20, 438–448.

Eisenegger, C. (2016). Role of dopamine D2 receptors in human reinforcement learning. Intrinsic Activity 4, A18.61.

Engelhard, B., Finkelstein, J., Cox, J., Fleming, W., Jang, H.J., Ornelas, S., Koay, S.A., Thiberge, S.Y., Daw, N.D., Tank, D.W., et al. (2019). Specialized coding of sensory, motor and cognitive variables in VTA dopamine neurons. Nature 570, 509–513.

Eshel, N., Bukwich, M., Rao, V., Hemmelder, V., Tian, J., and Uchida, N. (2015). Arithmetic and local circuitry underlying dopamine prediction errors. Nature 525, 243–246.

Everitt, B.J., Morris, K.A., O’Brien, A., and Robbins, T.W. (1991). The basolateral amygdala-ventral striatal system and conditioned place preference: further evidence of limbic-striatal interactions underlying reward-related processes. Neuroscience 42, 1–18.

Fee, M.S., and Goldberg, J.H. (2011). A hypothesis for basal ganglia-dependent reinforcement learning in the songbird. Neuroscience 198, 152–170.

Finn C., Abbeel, P. and Levine, S. (2017). RL^2: Model-Agnostic Meta-Learning for Fast Adaptation of Deep Networks. arXiv 1703.03400

Fisher, S.D., Robertson, P.B., Black, M.J., Redgrave, P., Sagar, M.A., Abraham, W.C., and Reynolds, J.N.J. (2017). Reinforcement determines the timing dependence of corticostriatal synaptic plasticity in vivo. Nat. Commun. 8, 334.

French, S.J., and Totterdell, S. (2003). Individual nucleus accumbens-projection neurons receive both basolateral amygdala and ventral subicular afferents in rats. Neuroscience 119, 19–31.

Fürth, D., Vaissière, T., Tzortzi, O., Xuan, Y., Märtin, A., Lazaridis, I., Spigolon, G., Fisone, G., Tomer, R., Deisseroth, K., et al. (2018). An interactive framework for whole-brain maps at cellular resolution. Nat. Neurosci. 21, 139–149.

Genovesio, A., Brasted, P.J., and Wise, S.P. (2006). Representation of future and previous spatial goals by separate neural populations in prefrontal cortex. J. Neurosci. 26, 7305–7316.

Gerfen, C.R., and Surmeier, D.J. (2011). Modulation of striatal projection systems by dopamine. Annu. Rev. Neurosci. 34, 441–466.

Gersch, T.M., Foley, N.C., Eisenberg, I., and Gottlieb, J. (2014). Neural Correlates of Temporal Credit Assignment in the Parietal Lobe.

Gershman, S.J., Moustafa, A.A., and Ludvig, E.A. (2014). Time representation in reinforcement learning models of the basal ganglia. Front. Comput. Neurosci. 7, 194.

Gerstner, W., Lehmann, M., Liakoni, V., Corneil, D., and Brea, J. (2018). Eligibility Traces and Plasticity on Behavioral Time Scales: Experimental Support of NeoHebbian Three-Factor Learning Rules. Front. Neural Circuits 12, 53.

Groenewegen, H.J., Becker, N.E., and Lohman, A.H. (1980). Subcortical afferents of the nucleus accumbens septi in the cat, studied with retrograde axonal transport of horseradish peroxidase and bisbenzimid. Neuroscience 5, 1903–1916.

Hahnloser, R.H.R., Kozhevnikov, A.A., and Fee, M.S. (2002). An ultra-sparse code underliesthe generation of neural sequences in a songbird. Nature 419, 65–70.

Hamid, A.A., Pettibone, J.R., Mabrouk, O.S., Hetrick, V.L., Schmidt, R., Vander Weele, C.M., Kennedy, R.T., Aragona, B.J., and Berke, J.D. (2016). Mesolimbic dopamine signals the value of work. Nat. Neurosci. 19, 117–126.

Harvey, C.D., Coen, P., and Tank, D.W. (2012). Choice-specific sequences in parietal cortex during a virtual-navigation decision task. Nature 484, 62–68.

Hazy, T.E., Frank, M.J., and O’Reilly, R.C. (2010). Neural mechanisms of acquired phasic dopamine responses in learning. Neurosci. Biobehav. Rev. 34, 701–720.

Hernandez, P.J., Sadeghian, K., and Kelley, A.E. (2002). Early consolidation of instrumental learning requires protein synthesis in the nucleus accumbens. Nat. Neurosci. 5, 1327–1331.

Hinton, G., Srivastava, N. and Swersky, K. (2012) Neural networks for machine learning lecture 6A overview of mini-batch gradient descent. Cited vol. 14, no. 8, p. 249.

Howard, M.W., and Eichenbaum, H. (2013). The hippocampus, time, and memory across scales. Journal of Experimental Psychology: General 142, 1211–1230.

Hunnicutt, B.J., Jongbloets, B.C., Birdsong, W.T., Gertz, K.J., Zhong, H., and Mao, T. (2016). A comprehensive excitatory input map of the striatum reveals novel functional organization. Elife 5.

Ito, M., and Doya, K. (2015). Parallel Representation of Value-Based and Finite State-Based Strategies in the Ventral and Dorsal Striatum. PLoS Comput. Biol. 11, e1004540.

Izquierdo, A., Wiedholz, L.M., Millstein, R.A., Yang, R.J., Bussey, T.J., Saksida, L.M., and Holmes, A. (2006). Genetic and dopaminergic modulation of reversal learning in a touchscreen-based operant procedure for mice. Behav. Brain Res. 171, 181–188.

Jin, D.Z., Fujii, N., and Graybiel, A.M. (2009). Neural representation of time in cortico-basal ganglia circuits. Proceedings of the National Academy of Sciences 106, 19156–19161.

Joel, D., Niv, Y., and Ruppin, E. (2002). Actor–critic models of the basal ganglia: new anatomical and computational perspectives. Neural Networks 15, 535–547.

Kalivas, P.W., Churchill, L., and Klitenick, M.A. (1993). GABA and enkephalin projection from the nucleus accumbens and ventral pallidum to the ventral tegmental area. Neuroscience 57, 1047–1060.

Kawai, T., Yamada, H., Sato, N., Takada, M., and Matsumoto, M. (2015). Roles of the lateral habenula and anterior cingulate cortex in negative outcome monitoring and behavioral adjustment in nonhuman primates. Neuron 88, 792–804.

Kelley, A.E., Smith-Roe, S.L., and Holahan, M.R. (1997). Response-reinforcement learning is dependent on N-methyl-D-aspartate receptor activation in the nucleus accumbens core. Proc. Natl. Acad. Sci. U. S. A. 94, 12174–12179.

Kim, C.K., Ye, L., Jennings, J.H., Pichamoorthy, N., Tang, D.D., Yoo, A.-C.W., Ramakrishnan, C., and Deisseroth, K. (2017). Molecular and circuit-dynamical identification of top-down neural mechanisms for restraint of reward seeking. Cell 170, 1013–1027.e14.

Kim, H., Sul, J.H., Huh, N., Lee, D., and Jung, M.W. (2009). Role of striatum in updating values of chosen actions. J. Neurosci. 29, 14701–14712.

Kondo, M., Kobayashi, K., Ohkura, M., Nakai, J., and Matsuzaki, M. (2017). Two-photon calcium imaging of the medial prefrontal cortex and hippocampus without cortical invasion. Elife 6.

Kozhevnikov, A.A., and Fee, M.S. (2007). Singing-related activity of identified HVC neurons in the zebra finch. J. Neurophysiol. 97, 4271–4283.

Krumin, M., Lee, J.J., Harris, K.D., and Carandini, M. (2018). Decision and navigation in mouse parietal cortex. Elife 7.

Kruzich, P.J., and Grandy, D.K. (2004). Dopamine D2 receptors mediate two-odor discrimination and reversal learning in C57BL/6 mice. BMC Neurosci. 5, 12.

Kruzich, P.J., Mitchell, S.H., Younkin, A., and Grandy, D.K. (2006). Dopamine D2 receptors mediate reversal learning in male C57BL/6J mice. Cognitive, Affective, & Behavioral Neuroscience 6, 86–90.

Kwak, S., Huh, N., Seo, J.-S., Lee, J.-E., Han, P.-L., and Jung, M.W. (2014). Role of dopamine D2 receptors in optimizing choice strategy in a dynamic and uncertain environment. Frontiers in Behavioral Neuroscience 8.

Lak, A., Okun, M., Moss, M.M., Gurnani, H., Farrell, K., Wells, M.J., Reddy, C.B., Kepecs, A., Harris, K.D., and Carandini, M. (2020). Dopaminergic and prefrontal basis of learning from sensory confidence and reward value. Neuron 105, 700–711.e6.

Lau, B., and Glimcher, P.W. (2008). Value representations in the primate striatum during matching behavior. Neuron 58, 451–463.

Lee, R.S., Mattar, M.G., Parker, N.F., Witten, I.B., and Daw, N.D. (2019). Reward prediction error does not explain movement selectivity in DMS-projecting dopamine neurons. eLife 8.

Leon, M.I., and Shadlen, M.N. (2003). Representation of Time by Neurons in the Posterior Parietal Cortex of the Macaque. Neuron 38, 317–327.

Long, M.A., Jin, D.Z., and Fee, M.S. (2010). Support for a synaptic chain model of neuronal sequence generation. Nature 468, 394–399.

Lovett-Barron, M., Chen, R., Bradbury, S., Andalman, A.S., Wagle, M., Guo, S., and Deisseroth, K. (2019). Multiple overlapping hypothalamus-brainstem circuits drive rapid threat avoidance.

Luk, C.-H., and Wallis, J.D. (2013). Choice coding in frontal cortex during stimulus-guided or action-guided decision-making. J. Neurosci. 33, 1864–1871.

MacAskill, A.F., Cassel, J.M., and Carter, A.G. (2014). Cocaine exposure reorganizes cell type- and input-specific connectivity in the nucleus accumbens. Nature Neuroscience 17, 1198–1207.

Maggi, S., and Humphries, M.D. (2019). Independent population coding of the present and the past in prefrontal cortex during learning.

Maggi, S., Peyrache, A., and Humphries, M.D. (2018). An ensemble code in medial prefrontal cortex links prior events to outcomes during learning. Nat. Commun. 9, 2204.

Matsumoto, M., and Hikosaka, O. (2009). Two types of dopamine neuron distinctly convey positive and negative motivational signals. Nature 459, 837–841.

Matsumoto, N., Minamimoto, T., Graybiel, A.M., and Kimura, M. (2001). Neurons in the Thalamic CM-Pf Complex Supply Striatal Neurons With Information About Behaviorally Significant Sensory Events. Journal of Neurophysiology 85, 960–976.

Montague, P.R., Dayan, P., and Sejnowski, T.J. (1996). A framework for mesencephalic dopamine systems based on predictive Hebbian learning. J. Neurosci. 16, 1936–1947.

Musall, S., Kaufman, M.T., Juavinett, A.L., Gluf, S., and Churchland, A.K. (2019). Single-trial neural dynamics are dominated by richly varied movements. Nat. Neurosci. 22, 1677–1686.

Nagabandi, A., Kahn, G., Fearing, R.S., and Levine, S. (2018). Neural Network Dynamics for Model-Based Deep Reinforcement Learning with Model-Free Fine-Tuning. 2018 IEEE International Conference on Robotics and Automation (ICRA).

Nestler, E.J. (2001). Molecular basis of long-term plasticity underlying addiction. Nat. Rev. Neurosci. 2, 119–128.

Nicola, S.M., Surmeier, J., and Malenka, R.C. (2000). Dopaminergic modulation of neuronal excitability in the striatum and nucleus accumbens. Annu. Rev. Neurosci. 23, 185–215.

O’Doherty, J., Dayan, P., Schultz, J., Deichmann, R., Friston, K., and Dolan, R.J. (2004). Dissociable roles of ventral and dorsal striatum in instrumental conditioning. Science 304, 452–454.

O’Doherty, J.P., Dayan, P., Friston, K., Critchley, H., and Dolan, R.J. (2003). Temporal difference models and reward-related learning in the human brain. Neuron 38, 329–337.

Ölveczky, B.P., Otchy, T.M., Goldberg, J.H., Aronov, D., and Fee, M.S. (2011). Changes in the neural control of a complex motor sequence during learning. J. Neurophysiol. 106, 386–397.

O’Neill, M., and Brown, V.J. (2007). The effect of striatal dopamine depletion and the adenosine A2A antagonist KW-6002 on reversal learning in rats. Neurobiol. Learn. Mem. 88, 75–81.

Otis, J.M., Namboodiri, V.M.K., Matan, A.M., Voets, E.S., Mohorn, E.P., Kosyk, O., McHenry, J.A., Robinson, J.E., Resendez, S.L., Rossi, M.A., et al. (2017). Prefrontal cortex output circuits guide reward seeking through divergent cue encoding. Nature 543, 103–107.

Otis, J.M., Zhu, M., Namboodiri, V.M.K., Cook, C.A., Kosyk, O., Matan, A.M., Ying, R., Hashikawa, Y., Hashikawa, K., Trujillo-Pisanty, I., et al. (2019). Paraventricular thalamus projection neurons integrate cortical and hypothalamic signals for cue-reward processing. Neuron 103, 277–290.e6.

Ottenheimer, D., Richard, J.M., and Janak, P.H. (2018). Ventral pallidum encodes relative reward value earlier and more robustly than nucleus accumbens. Nat. Commun. 9, 4350.

Ottenheimer, D.J., Wang, K., Haimbaugh, A., Janak, P.H., and Richard, J.M. (2019). Recruitment and disruption of ventral pallidal cue encoding during alcohol seeking. Eur. J. Neurosci. 50, 3428–3444.

Ottenheimer, D.J., Bari, B.A., Sutlief, E., Fraser, K.M., Kim, T.H., Richard, J.M., Cohen, J.Y., and Janak, P.H. (2020). A quantitative reward prediction error signal in the ventral pallidum. Nat. Neurosci. 23, 1267–1276.

Pan, W.-X., Schmidt, R., Wickens, J.R., and Hyland, B.I. (2005). Dopamine cells respond to predicted events during classical conditioning: evidence for eligibility traces in the reward-learning network. J. Neurosci. 25, 6235–6242.

Park, I.M., Meister, M.L.R., Huk, A.C., and Pillow, J.W. (2014). Encoding and decoding in parietal cortex during sensorimotor decision-making. Nature Neuroscience 17, 1395–1403.

Parker, N.F., Cameron, C.M., Taliaferro, J.P., Lee, J., Choi, J.Y., Davidson, T.J., Daw, N.D., and Witten, I.B. (2016). Reward and choice encoding in terminals of midbrain dopamine neurons depends on striatal target. Nat. Neurosci. 19, 845–854.

Parkinson, J.A., Olmstead, M.C., Burns, L.H., Robbins, T.W., and Everitt, B.J. (1999). Dissociation in effects of lesions of the nucleus accumbens core and shell on appetitive pavlovian approach behavior and the potentiation of conditioned reinforcement and locomotor activity by D-amphetamine. J. Neurosci. 19, 2401–2411.

Pastalkova, E., Itskov, V., Amarasingham, A., and Buzsáki, G. (2008). Internally generated cell assembly sequences in the rat hippocampus. Science 321, 1322–1327.

Paxinos, G., and Franklin, K.B.J. (2004). The Mouse Brain in Stereotaxic Coordinates (Gulf Professional Publishing).

Phillips, G.D., Le Noury, J., Wolterink, G., Donselaar-Wolterink, I., Robbins, T.W., and Everitt, B.J. (1993). Cholecystokinin-dopamine interactions within the nucleus accumbens in the control over behaviour by conditioned reinforcement. Behav. Brain Res. 55, 223–231.

Phillips, G.D., Robbins, T.W., and Everitt, B.J. (1994). Mesoaccumbens dopamine-opiate interactions in the control over behaviour by a conditioned reinforcer. Psychopharmacology 114, 345–359.

Phillipson, O.T., and Griffiths, A.C. (1985). The topographic order of inputs to nucleus accumbens in the rat. Neuroscience 16, 275–296.

Picardo, M.A., Merel, J., Katlowitz, K.A., Vallentin, D., Okobi, D.E., Benezra, S.E., Clary, R.C., Pnevmatikakis, E.A., Paninski, L., and Long, M.A. (2016). Population-level representation of a temporal sequence underlying song production in the zebra finch. Neuron 90, 866–876.

Pinto, L., and Dan, Y. (2015). Cell-type-specific activity in prefrontal cortex during goal-directed behavior. Neuron 87, 437–450.

Piray, P. (2011). The role of dorsal striatal D2-like receptors in reversal learning: a reinforcement learning viewpoint. J. Neurosci. 31, 14049–14050.

Pnevmatikakis, E.A., and Giovannucci, A. (2017). NoRMCorre: An online algorithm for piecewise rigid motion correction of calcium imaging data. J. Neurosci. Methods 291, 83–94.

Ponzi, A., and Wickens, J. (2010). Sequentially switching cell assemblies in random inhibitory networks of spiking neurons in the striatum. J. Neurosci. 30, 5894–5911.

Poulin, J.-F., Caronia, G., Hofer, C., Cui, Q., Helm, B., Ramakrishnan, C., Chan, C.S., Dombeck, D.A., Deisseroth, K., and Awatramani, R. (2018). Mapping projections of molecularly defined dopamine neuron subtypes using intersectional genetic approaches. Nat. Neurosci. 21, 1260–1271.

Rakelly, K., Zhou, A., Quillen, D., Finn, D. and Levine, D. (2019). Efficient Off-Policy Meta-Reinforcement Learning via Probabilistic Context Variables. arXiv 1903.08254

Reed, S.J., Lafferty, C.K., Mendoza, J.A., Yang, A.K., Davidson, T.J., Grosenick, L., Deisseroth, K., and Britt, J.P. (2018). Coordinated reductions in excitatory input to the nucleus accumbens underlie food consumption. Neuron 99, 1260–1273.e4.

Reynolds, J.N.J., and Wickens, J.R. (2002). Dopamine-dependent plasticity of corticostriatal synapses. Neural Netw. 15, 507–521.

Richard, J.M., Ambroggi, F., Janak, P.H., and Fields, H.L. (2016). Ventral Pallidum Neurons Encode Incentive Value and Promote Cue-Elicited Instrumental Actions. Neuron 90, 1165–1173.

Robbins, T.W., Cador, M., Taylor, J.R., and Everitt, B.J. (1989). Limbic-striatal interactions in reward-related processes. Neuroscience & Biobehavioral Reviews 13, 155–162.

Roitman, M.F., Wheeler, R.A., and Carelli, R.M. (2005). Nucleus accumbens neurons are innately tuned for rewarding and aversive taste stimuli, encode their predictors, and are linked to motor output. Neuron 45, 587–597.

Russo, S.J., Dietz, D.M., Dumitriu, D., Morrison, J.H., Malenka, R.C., and Nestler, E.J. (2010). The addicted synapse: mechanisms of synaptic and structural plasticity in nucleus accumbens. Trends in Neurosciences 33, 267–276.

Rutledge, R.B., Lazzaro, S.C., Lau, B., Myers, C.E., Gluck, M.A., and Glimcher, P.W. (2009). Dopaminergic drugs modulate learning rates and perseveration in Parkinson’s patients in a dynamic foraging task. Journal of Neuroscience 29, 15104–15114.

Sæmundsson, S., Hofmann, K. and Deisenroth, M.P. (2018). Meta Reinforcement Learning with Latent Variable Gaussian Processes. arXiv 1803.07551

Sabatini, B.L. (2019). The impact of reporter kinetics on the interpretation of data gathered with fluorescent reporters.

Sadacca, B.F., Jones, J.L., and Schoenbaum, G. (2016). Midbrain dopamine neurons compute inferred and cached value prediction errors in a common framework. Elife 5.

Sakata, J.T., Hampton, C.M., and Brainard, M.S. (2008). Social modulation of sequence and syllable variability in adult birdsong. J. Neurophysiol. 99, 1700–1711.

Salamone, J.D., Steinpreis, R.E., McCullough, L.D., Smith, P., Grebel, D., and Mahan, K. (1991). Haloperidol and nucleus accumbens dopamine depletion suppress lever pressing for food but increase free food consumption in a novel food choice procedure. Psychopharmacology 104, 515–521.

Saunders, B.T., Richard, J.M., Margolis, E.B., and Janak, P.H. (2018). Dopamine neurons create Pavlovian conditioned stimuli with circuit-defined motivational properties. Nat. Neurosci. 21, 1072–1083.

Schultz, W. (1998). Predictive reward signal of dopamine neurons. J. Neurophysiol. 80, 1–27.

Schultz, W., Dayan, P., and Montague, P.R. (1997). A neural substrate of prediction and reward. Science 275, 1593–1599.

Seo, M., Lee, E., and Averbeck, B.B. (2012). Action selection and action value in frontal-striatal circuits. Neuron 74, 947–960.

Setlow, B., Schoenbaum, G., and Gallagher, M. (2003). Neural encoding in ventral striatum during olfactory discrimination learning. Neuron 38, 625–636.

Shen, W., Flajolet, M., Greengard, P., and Surmeier, D.J. (2008). Dichotomous dopaminergic control of striatal synaptic plasticity. Science 321, 848–851.

Shonesy, B.C., Stephenson, J.R., Marks, C.R., and Colbran, R.J. (2020). Cyclic AMP-dependent protein kinase and D1 dopamine receptors regulate diacylglycerol lipase-α and synaptic 2-arachidonoyl glycerol signaling. J. Neurochem. 153, 334–345.

Siniscalchi, M.J., Wang, H., and Kwan, A.C. (2019). Enhanced population coding for rewarded choices in the medial frontal cortex of the mouse. Cereb. Cortex 29, 4090–4106.

Song, H.F., Yang, G.R., and Wang, X.-J. (2017). Reward-based training of recurrent neural networks for cognitive and value-based tasks. Elife 6.

Steinberg, E.E., Keiflin, R., Boivin, J.R., Witten, I.B., Deisseroth, K., and Janak, P.H. (2013). A causal link between prediction errors, dopamine neurons and learning. Nat. Neurosci. 16, 966–973.

Steinmetz, N.A., Zatka-Haas, P., Carandini, M., and Harris, K.D. (2019). Distributed coding of choice, action and engagement across the mouse brain. Nature 576, 266–273.

Stephenson-Jones, M., Bravo-Rivera, C., Ahrens, S., Furlan, A., Xiao, X., Fernandes-Henriques, C., and Li, B. (2020). Opposing contributions of GABAergic and glutamatergic ventral pallidal neurons to motivational behaviors. Neuron 105, 921–933.e5.

Stuber, G.D., Sparta, D.R., Stamatakis, A.M., van Leeuwen, W.A., Hardjoprajitno, J.E., Cho, S., Tye, K.M., Kempadoo, K.A., Zhang, F., Deisseroth, K., et al. (2011). Excitatory transmission from the amygdala to nucleus accumbens facilitates reward seeking. Nature 475, 377.

Sul, J.H., Kim, H., Huh, N., Lee, D., and Jung, M.W. (2010). Distinct roles of rodent orbitofrontal and medial prefrontal cortex in decision making. Neuron 66, 449–460.

Suri, R.E., and Schultz, W. (1998). Learning of sequential movements by neural network model with dopamine-like reinforcement signal. Experimental Brain Research 121, 350–354.

Suri, R.E., and Schultz, W. (1999). A neural network model with dopamine-like reinforcement signal that learns a spatial delayed response task. Neuroscience 91, 871–890.

Sutton, R.S. (1988). Learning to predict by the methods of temporal differences. Mach. Learn. 3, 9–44.

Sutton, R.S., and Barto, A.G. (1998). Reinforcement Learning: An Introduction (MIT Press).

Swanson, L.W. (1982). The projections of the ventral tegmental area and adjacent regions: a combined fluorescent retrograde tracer and immunofluorescence study in the rat. Brain Res. Bull. 9, 321–353.

Taghzouti, K., Louilot, A., Herman, J.P., Le Moal, M., and Simon, H. (1985). Alternation behavior, spatial discrimination, and reversal disturbances following 6-hydroxydopamine lesions in the nucleus accumbens of the rat. Behav. Neural Biol. 44, 354–363.

Tai, L.-H., Lee, A.M., Benavidez, N., Bonci, A., and Wilbrecht, L. (2012). Transient stimulation of distinct subpopulations of striatal neurons mimics changes in action value. Nat. Neurosci. 15, 1281–1289.

Taylor, J., and Robbins, T. (1986). 6-Hydroxydopamine lesions of the nucleus accumbens, but not of the caudate nucleus, attenuate enhanced responding with reward-related stimuli produced by intra-accumbens d-amphetamine. Psychopharmacology 90.

Terada, S., Sakurai, Y., Nakahara, H., and Fujisawa, S. (2017). Temporal and Rate Coding for Discrete Event Sequences in the Hippocampus. Neuron 94, 1248–1262.e4.

Tesauro, G. (1992). Practical issues in temporal difference learning. Reinforcement Learning 33–53.

Thévenaz, P., Ruttimann, U.E., and Unser, M. (1998). A pyramid approach to subpixel registration based on intensity. IEEE Trans. Image Process. 7, 27–41.

Tian, J., Huang, R., Cohen, J.Y., Osakada, F., Kobak, D., Machens, C.K., Callaway, E.M., Uchida, N., and Watabe-Uchida, M. (2016). Distributed and mixed information in monosynaptic inputs to dopamine neurons. Neuron 91, 1374–1389.

Tindell, A.J., Berridge, K.C., and Aldridge, J.W. (2004). Ventral pallidal representation of pavlovian cues and reward: population and rate codes. J. Neurosci. 24, 1058–1069.

Tsai, H.-C., -C. Tsai, H., Zhang, F., Adamantidis, A., Stuber, G.D., Bonci, A., de Lecea, L., and Deisseroth, K. (2009). Phasic firing in dopaminergic neurons Is sufficient for behavioral conditioning. Science 324, 1080–1084.

Tsitsiklis, J.N., and Van Roy, B. (1997). An analysis of temporal-difference learning with function approximation. IEEE Transactions on Automatic Control 42, 674–690.

Tsutsui, K.-I., Grabenhorst, F., Kobayashi, S., and Schultz, W. (2016). A dynamic code for economic object valuation in prefrontal cortex neurons. Nat. Commun. 7, 12554.

Wan, X., and Peoples, L.L. (2006). Firing patterns of accumbal neurons during a pavlovian-conditioned approach task. J. Neurophysiol. 96, 652–660.

Wang, J.X., Kurth-Nelson, Z., Kumaran, D., Tirumala, D., Soyer, H., Leibo, J.Z., Hassabis, D., and Botvinick, M. (2018). Prefrontal cortex as a meta-reinforcement learning system. Nat. Neurosci. 21, 860–868.

Watabe-Uchida, M., Zhu, L., Ogawa, S.K., Vamanrao, A., and Uchida, N. (2012). Whole-brain mapping of direct inputs to midbrain dopamine neurons. Neuron 74, 858–873.

Wei, Wei, W., and Wang, X.-J. (2016). Inhibitory control in the cortico-basal ganglia-thalamocortical loop: complex regulation and interplay with memory and decision processes. Neuron 92, 1093–1105.

Wilson, C.J. (2004). Basal Ganglia. The Synaptic Organization of the Brain 361–414.

Witten, I.B., Steinberg, E.E., Lee, S.Y., Davidson, T.J., Zalocusky, K.A., Brodsky, M., Yizhar, O., Cho, S.L., Gong, S., Ramakrishnan, C., et al. (2011). Recombinase-driver rat lines: tools, techniques, and optogenetic application to dopamine-mediated reinforcement. Neuron 72, 721–733.

Wolff, S.B.E., Ko, R. and Ölvezky, B.P. (2019) Distinct roles for motor cortical and thalamic inputs to striatum during motor learning and execution. bioRxiv https://doi.org/10.1101/825810.

Wörgötter, F., and Porr, B. (2005). Temporal sequence learning, prediction, and control: a review of different models and their relation to biological mechanisms. Neural Comput. 17, 245–319.

Wright, C.I., and Groenewegen, H.J. (1995). Patterns of convergence and segregation in the medial nucleus accumbens of the rat: relationships of prefrontal cortical, midline thalamic, and basal amygdaloid afferents. J. Comp. Neurol. 361, 383–403.

Xiong, Q., Znamenskiy, P., and Zador, A. (2015). Selective corticostriatal plasticity during acquisition of an auditory discrimination task. Nature 521, 348–351.

Yagishita, S., Hayashi-Takagi, A., Ellis-Davies, G.C.R., Urakubo, H., Ishii, S., and Kasai, H. (2014). A critical time window for dopamine actions on the structural plasticity of dendritic spines. Science 345, 1616–1620.

Yang, H., de Jong, J.W., Tak, Y., Peck, J., Bateup, H.S., and Lammel, S. (2018). Nucleus accumbens subnuclei regulate motivated behavior via direct inhibition and disinhibition of VTA dopamine subpopulations. Neuron 97, 434–449.e4.

Young, C.E., and Yang, C.R. (2005). Dopamine D1-like receptor modulates layer- and frequency-specific short-term synaptic plasticity in rat prefrontal cortical neurons. European Journal of Neuroscience 21, 3310–3320.

Zhou, P., Resendez, S.L., Rodriguez-Romaguera, J., Jimenez, J.C., Neufeld, S.Q., Giovannucci, A., Friedrich, J., Pnevmatikakis, E.A., Stuber, G.D., Hen, R., et al. (2018). Efficient and accurate extraction of in vivo calcium signals from microendoscopic video data. Elife 7.

Zhou, S., Masmanidis, S.C., and Buonomano, D.V. (2020). Neural Sequences as an Optimal Dynamical Regime for the Readout of Time. Neuron 108, 651–658.e5.

Zhu, Y., Wienecke, C.F.R., Nachtrab, G., and Chen, X. (2016). A thalamic input to the nucleus accumbens mediates opiate dependence. Nature 530, 219–222.

Zhu, Y., Nachtrab, G., Keyes, P.C., Allen, W.E., Luo, L., and Chen, X. (2018). Dynamic salience processing in paraventricular thalamus gates associative learning. Science 362, 423–429.

